# When Can Brain Connectivity Track the Working Mind? A Large-Scale Benchmark of Dynamic Functional Connectivity Across Cognitive Paradigms

**DOI:** 10.64898/2026.06.28.735101

**Authors:** Mohammad Torabi, Jean-Baptiste Poline, Georgios D. Mitsis

## Abstract

Dynamic functional connectivity (dFC) — the time-varying re-configuration of brain network interactions — has become a widely adopted method for studying how neural dynamics reflect ongoing cognition. Yet a fundamental question remains unresolved: can dFC reliably track when a person is cognitively engaged, and if not, why does it fail? Here, we address this question through a large-scale benchmark of seven widely used dFC methods, evaluating how well each predicts task presence across 16 fMRI datasets encompassing over 1,500 participants and 28 distinct experimental settings, complemented by realistic simulated data.

Across experimental data, dFC-based tracking of cognitive engagement was unreliable in many cases: most method–experiment combinations performed near chance, and no single method succeeded across all contexts. This failure, however, was not uniform. Both experimental and simulated data showed that decoding performance varied systematically with three interacting factors — experimental design, data quality, and the choice of dFC method — rather than depending on dFC features alone.

Critically, we identify specific experimental design conditions associated with more reliable tracking: paradigms with longer, more regular task blocks and fewer task–rest transitions were substantially more decodable, while data quality independently influenced performance across methods. These findings offer actionable principles for when dFC can — and cannot — be expected to serve as a reliable marker of underlying cognitive states.

## Introduction

A central goal of cognitive neuroscience is to understand how the brain dynamically encodes and transitions between mental states. Dynamic functional connectivity (dFC) — the time-varying pattern of coordinated activity across brain networks — has emerged as a promising window into this process [1–5]. Unlike static measures of connectivity, dFC aims to track the reconfiguration of neural networks as cognition unfolds in real time, offering potential as both a scientific tool for studying cognition and biomarker for neurological and psychiatric conditions.

Despite widespread adoption, the validity of dFC as a measure of ongoing cognitive states remains poorly established [6–8]. Because most dFC studies use resting-state data — where no ground truth exists for what the brain “should” be doing — it has proven difficult to determine whether observed fluctuations in connectivity reflect genuine cognitive dynamics or merely motion/physiological noise and measurement artefacts [6, 9]. This ambiguity is compounded by the substantial “analytical flexibility” related to how dFC is computed: different methodologies applied to the same dataset may yield dramatically variable results, with intermethod variability often comparable in magnitude to the temporal fluctuations being studied [10]. The practical implication is alarming — a large body of published findings may critically depend on - sometimes arbitrary-methodological choices, with no principled basis for knowing which methods, if any, are reliably tracking mental state.

This problem cannot be resolved by studying resting-state data alone. Pioneering work established that task-based paradigms can be used to this end: Gonzalez-Castillo et al. [11] showed that dFC states can track distinct mental tasks (e.g., math vs. memory) at the block level, and Xie et al. [8] used multitask datasets to compare methods in terms of their ability to segment time series into cognitively homogeneous periods. These contributions validated the logic of using tasks as ground truth — but 1) they were implemented at the timescale of sustained, minute-long task blocks, where cognitive states are stable and contrasts between conditions are large, and 2) validated on a single dataset. A more demanding and ultimately more informative test is whether dFC features, assigned to individual time points, can track cognitive engagement at a finer temporal scale than sustained block-level contrasts — regardless of the temporal resolution intrinsic to each method. This requires detecting not only the broad contrast between two sustained cognitive states — as in block-level comparisons — but the moment-to-moment shift between being actively engaged in a task and being at rest. Second, dFC validation should be performed across idiosyn-crasies of datasets and cognitive paradigms as dFC methods’ performance may depend on data characteristics. Last, we propose to frame dFC validation in a predictive framework, making results easily interpretable and statistically sound.

Here, we address this question directly at an unprecedented scale. We treat the binary distinction between task engagement and rest not merely as a classification problem, but as a principled validity criterion: if a dFC method is truly sensitive to moment-to-moment cognitive states, it must reliably predict task-present from rest periods even within a single paradigm. We benchmark seven widely used dFC methodologies — spanning state-free approaches (Sliding Window, Time-Frequency) and state-based approaches (Co-Activation Patterns, Hidden Markov Models, Clustering, Windowless) — across 16 independent fMRI datasets comprising over 1,500 participants and 28 experimental settings drawn from the OpenNeuro repository, using our open-source `PydFC` toolbox [12].

Our results deliver a clear and consequential message. In experimental data, dFC-based tracking of cognitive engagement was unreliable in many cases; most method–experiment combinations performed near chance, and no dFC assessment method was universally effective. For most combinations, decoding accuracy scarcely exceeded chance level. Nevertheless, some methods were indeed effective at predicting experimental conditions on some specific datasets.

This suggests that the limiting factors are not solely intrinsic to the dFC approach, but also include experimental design and methodological choice. By comparing the empirically acquired (experimental) data with simulated data, and analysing what separated high- from low-performing experiments for each method, we identified three interacting determinants of success: experimental design (particularly task block length and transition frequency), data quality (the degree to which motion, physiological artefacts, and other sources of signal degradation compromise the fMRI signal), and the choice of dFC method. Together, these factors significantly influence the likelihood of reliable dFC-based cognitive state tracking — though how they interact and generalize across broader contexts remains an open question.

These findings have broad implications for how cognitive neuroscience uses connectivity dynamics as a window into mental states. They provide a framework for addressing the uncertainty about whether dFC fluctuations reflect genuine cognitive dynamics or merely noise and measurement artefacts, examine whether widely adopted dFC methods are actually valid for tracking cognitive engagement in real time, and offer researchers actionable design principles that may increase the likelihood of reliable dFC-based cognitive state tracking. More fundamentally, this work provides a multi datasets and time resolved predictive validation framework to evaluate dFC methods and their interactions with paradigms across 28 experiments.

## Results

To determine whether dFC can reliably track cognitive engagement as it unfolds, and identify when and why it fails, we benchmarked seven dFC methodologies across 28 task-fMRI experiments (`EXP.1`–`EXP.28`), each treated as an independent evaluation setting. We used the binary transition between task and rest as a ground-truth validity criterion, assessing each method’s ability to decode cognitive state at the resolution of individual fMRI time points (TRs). Decoding performance was quantified using balanced accuracy (chance level = 50%), and intrinsic task-rest separability was assessed using the Silhouette Index (SI). Together, these metrics reveal both the reliability of dFC as a cognitive state tracker and the structure of the underlying feature space. For an illustration of the full prediction pipeline, see Figure 1.

**Fig. 1.**
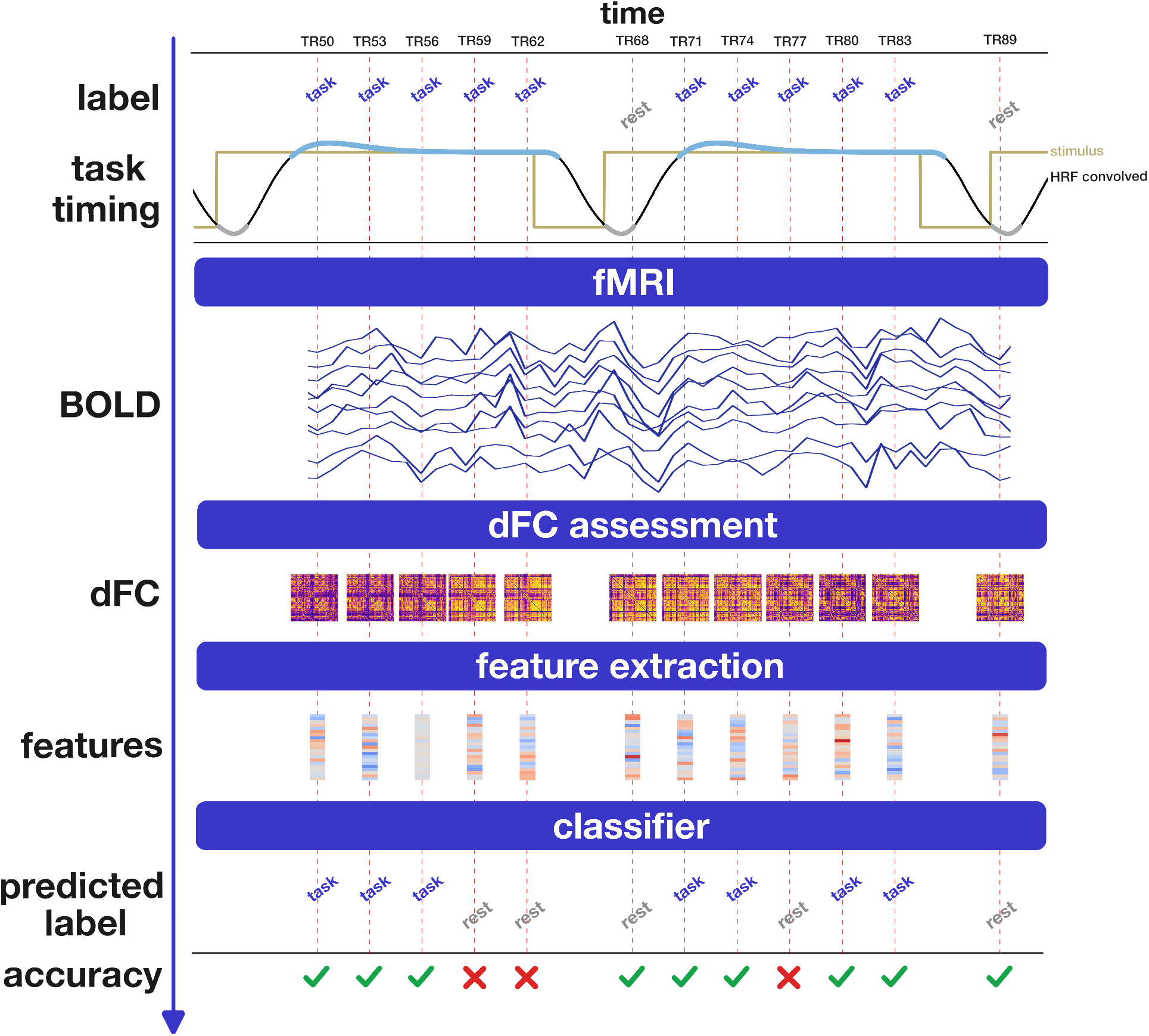
From stimuli to predicted task presence: a step-by-step illustration of how dFC features are extracted, reduced, and classified at the level of individual TRs. This figure illustrates the stages of analysis for `sub-101` from `EXP.27` (`ds005038`) performing the *Task-Localizer* paradigm (TR 50-89; selected TRs: 50, 53, 56, 59, 62, 68, 71, 74, 77, 80, 83, 89). From top to bottom, the workflow shows: **(1) Task timing:** original stimulus block timing extracted from the `events.tsv` file is shown in gold, with the HRF-convolved time course in black. Portions of the convolved signal identified as task by Gaussian Mixture Model (GMM) are highlighted in light blue, and rest periods in grey. Only TRs corresponding to these selected areas are retained for prediction. The true labels for each displayed TR are displayed as text (“rest” in grey, “task” in blue). **(2) BOLD time series:** TR-aligned BOLD signals from the first 10 regions of the Schaefer atlas. **(3) dFC patterns:** dFC patterns represented as connectivity matrices for each TR generated by the specified dFC method. The dFC patterns shown here were obtained using the Sliding Window method. **(4) Feature extraction:** for state-based methods (ContinuousHMM, DiscreteHMM, CAP, Clustering), features are based on state probabilities, distances from states, or mixing weights; for state-free methods (Sliding Window, Time-Frequency), features are extracted using Partial Least Squares (PLS) to reduce dimensionality. The features shown here were obtained using the Sliding Window method. **(5) Prediction:** extracted features on left out data are fed as inputs to the classifier, producing predicted labels for each TR (“rest” in grey, “task” in blue). **(6) Accuracy assessment:** predicted labels are compared to true labels; correct predictions are indicated by a green checkmark, incorrect predictions by a red cross (X). Across stages, elements corresponding to each selected TR are aligned vertically, with the vertical red dashed line marking the TR being displayed. TR indices are shown at the top to indicate temporal progression from left to right. This visualization demonstrates the temporal alignment and data flow from stimuli to predicted task labels in the pipeline for a single subject and task segment.

### DFC Patterns Do not Always Track Cognitive Engagement in Experimental Data

In experimental fMRI data, dFC-based decoding of ongoing cognitive engagement was unreliable across the majority of method–experiment combinations (Figure 2). Crucially, this near-chance decoding cannot be attributed to an absence of task-evoked signal in the underlying data: across experiments, the task-rest BOLD contrast (Cohen’s *d*, computed per region and taken as the maximum across regions) ranged from 0.19 to 2.06 (median 0.46), with nearly all experiments (27 of 28, 96%) exceed-ing 0.25 and about half (13 of 28, 46%) exceeding 0.5 (Supplementary Figure 9). A measurable task-evoked response was therefore present in essentially every experiment, yet this did not translate into reliable single-TR decoding from dFC — and a strong contrast was not sufficient on its own (e.g., `EXP.2`, with the largest contrast, was decoded near chance). Most combinations yielded accuracies near chance (~50%), with only 3 out of 196 combinations exceeding 70% and 18 out of 196 significantly above chance. This near-chance performance is not a marginal effect: it reflects a systematic failure of the examined dFC methods to detect the cognitive transition between engagement and rest in experimental data, even when a task-evoked signal is demonstrably present.

**Fig. 2.**
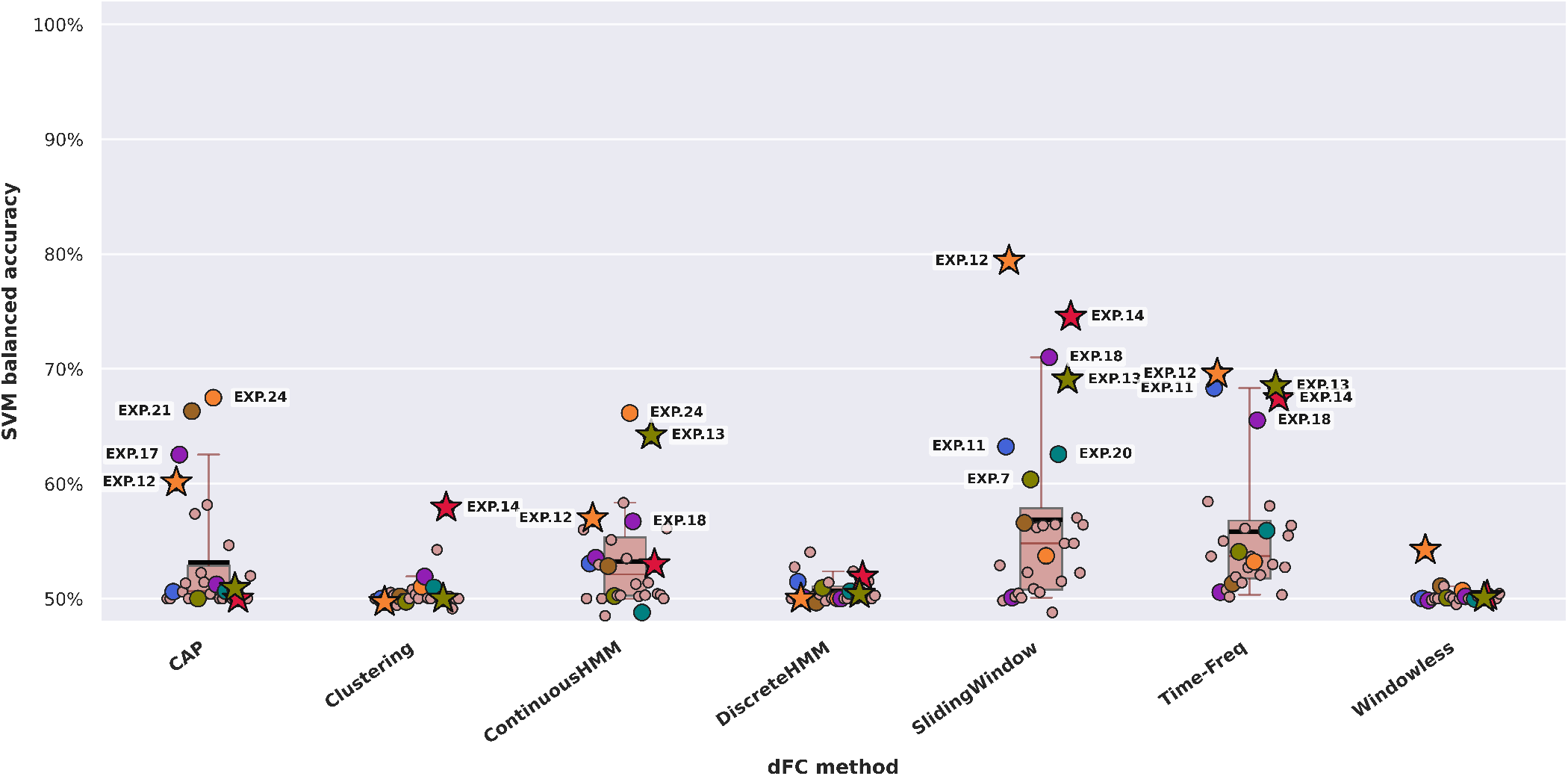
Single-TR task prediction performance varies substantially across dFC methods and experiments. Test-set balanced accuracy for task vs. rest classification is shown for each dFC method across 28 experiments (`EXP.1`-`EXP.28`), using PLS-transformed features and a Support Vector Machine (SVM) classifier. Each boxplot summarizes performance across experiments, with individual points representing experiments and median (red line) and mean (black line) balanced accuracy values indicated. Experiments achieving >60% accuracy for at least one method are highlighted across all methods, and the three highest-performing experiments overall (`EXP.12`, `EXP.13`, `EXP.14`) are marked with a star. Selected high-performing experiments are annotated to aid comparison. Overall performance is modest, with many experiment-method combinations yielding near chance performance (50%). Only 3 out of 196 experiment-method combinations exceed 70% accuracy, and relatively few exceed 60% (18/196), suggesting that reliable single-TR decoding is not the norm. Performance is strongly experiment-dependent. `EXP.12`, `EXP.13`, and `EXP.14` consistently rank among the top performers for Sliding Window and Time-Frequency methods, but their performance is not uniform across methods, dropping to near chance in some cases (e.g., Clustering and Discrete HMM). In contrast, `EXP.24` achieves the highest accuracy for CAP and Continuous HMM, while remaining in the lower half for Sliding Window and Time-Frequency. Method-dependent trends are also evident. Sliding Window achieves the highest central tendency and yields the largest number of above chance decoding performance occurrences, including three cases above 70%, but with substantial variability across experiments. Time-Frequency and Continuous HMM exhibit an overall more consistent performance, with most experiments falling between 50% and 60% and median accuracy around 55%, although Time-Frequency results in a higher number of higher-performing cases (~65-70%) and a higher mean. CAP displays a different pattern, with median performance close to chance but a small subset of experiments exceeding 60%. In contrast, Clustering, Discrete HMM, and Windowless methods remain predominantly near chance across almost all experiments. Together, these results highlight two key observations: (i) the detectability of task engagement varies markedly across experiments, and (ii) the relative performance of dFC methods is highly context-dependent, reinforcing the need for systematic benchmarking. For exact balanced accuracy values across all experiment-method pairs, see Supplementary Figure 38, and for detailed across-run variability, see Supplementary Figure 39.

A small cluster of experiments (`EXP.12`, `EXP.13`, `EXP.14`), all drawn from the same dataset (`ds003465`), achieved peak accuracies of 70–79% when using the Sliding Window and Time-Frequency methods. Even these relatively favorable cases, however, were not robust across methods: performance dropped to near chance for Clustering and Discrete HMM applied to the same experiments. This indicates that high performance is method-dependent rather than a stable property of cognitive engagement itself, and may arise from a specific alignment between experimental structure and methodological assumptions.

These favorable cases, however, were the exception. The majority of experiments (18 out of 28) were not reliably decoded by any method, with accuracies below 60% and in many cases close to chance. Reliable single-TR decoding thus emerged only for specific combinations of experiment and method, and was rare across the benchmark as a whole. The examined dFC assessment methods also differed in their overall performance profiles (Figure 2). Sliding Window and Time-Frequency showed the highest central tendency in experimental data. CAP and Continuous HMM exhibited occasional above-chance results — exceeding 65% in a few experiments — but lower overall reliability in experimental data. Clustering, Discrete HMM, and Windowless methods remained near chance across virtually all experiments. Notably, no single method exhibited consistently reliable performance across all the examined experiments.

These findings establish a sobering baseline: reliable single-TR decoding of cognitive engagement from dFC is the exception, not the norm, across a diverse landscape of cognitive paradigms and methods.

### Simulated Data Suggest That dFC Can Track Cognition Under Favorable Conditions

The near-chance performance observed in experimental data raises a fundamental question: does this reflect an intrinsic limitation of dFC as a cognitive state tracker, or does it reflect the compounding influence of experimental design constraints and data quality in experimental fMRI data? To probe this, we generated simulated fMRI data using a biophysical network model (The Virtual Brain), in which task engagement was modeled by stimulating a fixed set of brain regions during task-on periods. Inter-subject variability was introduced by adding noise to the model’s structural connectivity and the strength of interactions between regions. In this setting, the task signal is known and the motion, physiological artefacts, and other sources of signal degradation present in experimental BOLD data are absent, providing controlled conditions in which the effects of methodology and experimental design can be examined in relative isolation from the experimental data artefacts. By construction, these simulations also carried a strong task-evoked signal (Supplementary Figure 12): the task-rest BOLD contrast (Cohen’s d, maximum across regions) ranged from 0.26 to 8.66 (median 5.44), with 11 of 13 simulated experiments exceeding 2.0 — substantially larger than in the experimental data (median 0.46). These simulations were therefore designed as a favorable test: a setting in which, if a method is capable of tracking task engagement at all, a strong and clean signal should allow it to do so.

Under these controlled conditions, decoding performance improved substantially across all high-performing methods, with many experiment–method combinations achieving high or near-perfect accuracy (Figure 3). CAP exhibited the highest improvement (mean *>* 90%, median *>* 95%), fre-quently reaching perfect accuracy. Sliding Window and Time-Frequency also performed strongly (mean ~ 80%, median ~ 85%), while Continuous HMM exhibited moderateto-high performance with substantial variability. In sharp contrast, Clustering, Discrete HMM, and Windowless methods remained near chance even under these controlled conditions (median ~ 55%), failing to track task engagement despite the strong simulated signal (Cohen’s *d* up to 8.66) — a clear indication that their limitations are primarily methodological rather than driven by data quality or design.

**Fig. 3.**
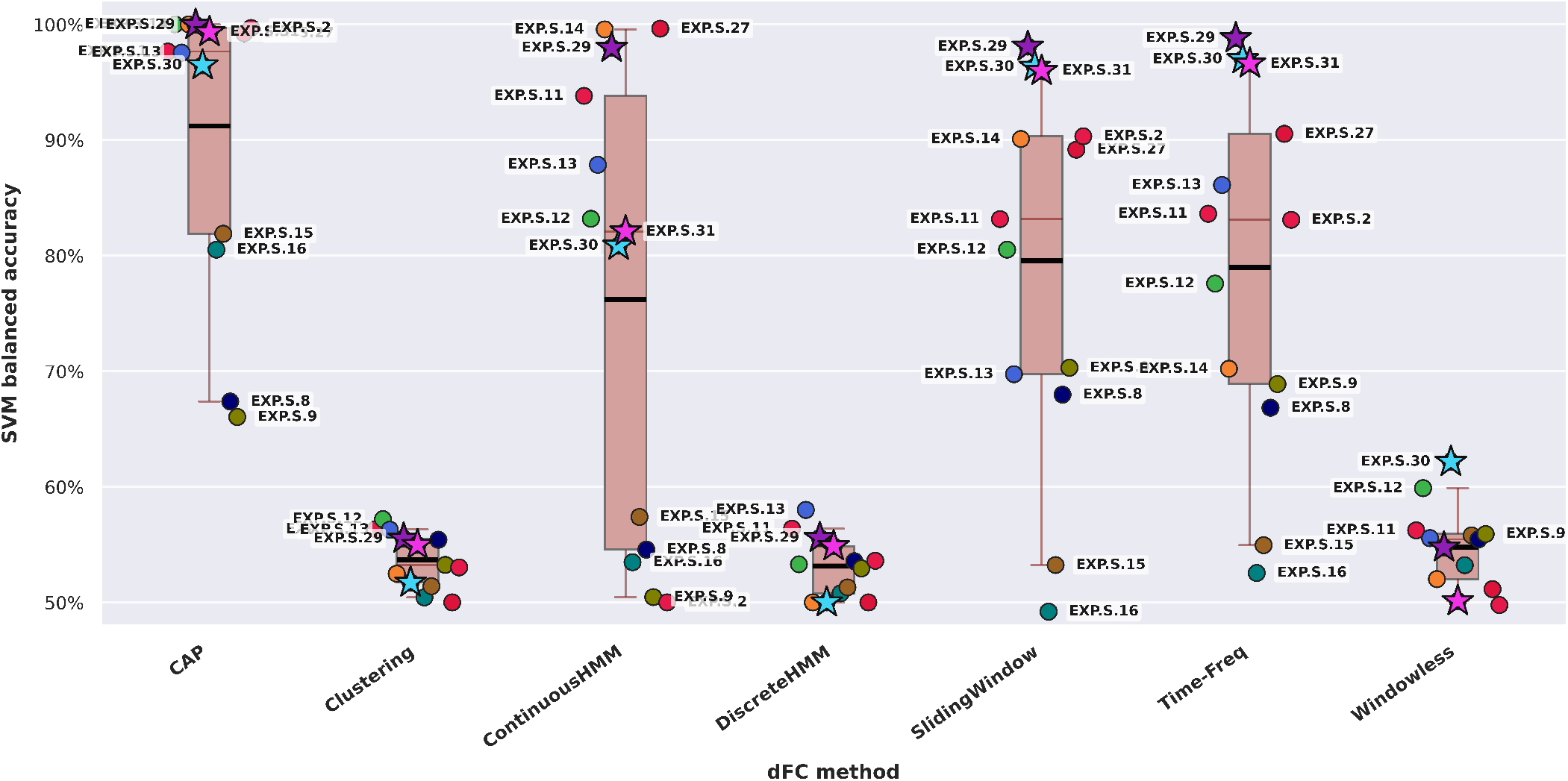
Single-TR task prediction performance improves substantially under simulated conditions. Test-set balanced accuracy for task vs. rest classification is shown for each dFC method across simulated experiments (`EXP.S.2`-`EXP.S.31`), using PLS-transformed features and an SVM classifier. Each boxplot summarizes performance across experiments, with individual points representing experiments and median (red line) and mean (black line) balanced accuracy values indicated. Simulated experiments include both counterparts of some real-world experiments and fully synthetic designs (`EXP.S.29`-`EXP.S.31`). The highest-performing experiments across all experimentmethod combinations (`EXP.S.29`-`EXP.S.31`) are marked with a star. Overall performance is markedly higher than in experimental data, with many experiment-method combinations achieving high or near-perfect accuracy. CAP consistently achieves the strongest performance, with mean and median above 90% (median >95%) and 9 out of 13 experiments reaching near-perfect or perfect accuracy, including multiple cases of 100% accuracy. Sliding Window and Time-Frequency also show strong performance, with mean accuracy around 80% and median close to 85%, and largely similar behavior across experiments. Sliding Window, Time-Frequency, and Continuous HMM all exhibit substantial variability, ranging from near chance to near-perfect accuracy. In contrast, Clustering, Discrete HMM, and Windowless methods remain close to chance level, with mean and median around 55%, indicating limited discriminative power even under simulated conditions. Performance remains experiment-dependent, although to a lesser extent than in experimental data. Fully synthetic experiments (`EXP.S.29`-`EXP.S.31`) consistently achieve high performance across multiple methods, with `EXP.S.29` reaching near-perfect or perfect accuracy across CAP, Continuous HMM, Sliding Window, and Time-Frequency. Conversely, a small subset of experiments (`EXP.S.8`, `EXP.S.9`, `EXP.S.15`, `EXP.S.16`) exhibit lower performance across methods. Aside from these cases, most experiments achieve near-perfect accuracy in at least one method, typically CAP. These results indicate that under simulated conditions, dFC features can encode task engagement, suggesting that the limitations observed in experimental data are not attributable to the dFC features alone, but also reflect experimental design, motion and physiological noise, and overall data complexity. For exact balanced accuracy values across all experiment-method pairs, see Supplementary Figure 46.

The gap between experimental and simulated performance suggests that the failure observed in experimental data is driven substantially by factors that are absent or reduced in the simulation — including arousal fluctuations, motion and physiological artefacts, variability in the hemodynamic and physiological response, and other sources of signal degradation — rather than being attributable solely to the dFC features themselves [8, 13–15]. We note, however, that the generative model does not capture all properties of real-world fMRI, so this interpretation holds to the extent that the simulation reflects the relevant structure of experimental data.

Stratifying simulated experiments by category further il-luminates how experimental design and data quality contribute independently to performance (Figure 4). We grouped the simulated experiments into three types: fully synthetic periodic designs, and simulations that reproduce the task timing of experiments that had performed either well (`EXP.11`–`EXP.14`) or poorly (e.g., `EXP.2`, `EXP.8`, `EXP.9`, `EXP.15`, `EXP.16`, `EXP.27`) in our experimental data analyses. Synthetic periodic experiments yielded the highest accuracy across methods, while simulations derived from poorly-performing experiments resulted in substantially reduced accuracy — even under controlled conditions. Comparing performance across these categories helps dissociate the two factors: Continuous HMM performance drops to near chance for simulations derived from poorly-performing experiments, suggesting that in these cases experimental design is the primary limiting factor — a constraint shared between experimental and simulated data. In contrast, CAP, Sliding Window, and Time-Frequency maintain substantially higher performance in simulated than in experimental data for both the strongand weak-performing categories, suggesting that their reduced performance in experimental datasets reflects not only experimental design effects but also the additional influence of data quality. More generally, performance differences that persist between experimental and simulated data point to the role of data quality — i.e., the influence of motion, physiological artefacts, and other sources of signal degradation in real-world fMRI — whereas performance patterns that remain similar across both conditions indicate constraints imposed by experimental design.

**Fig. 4.**
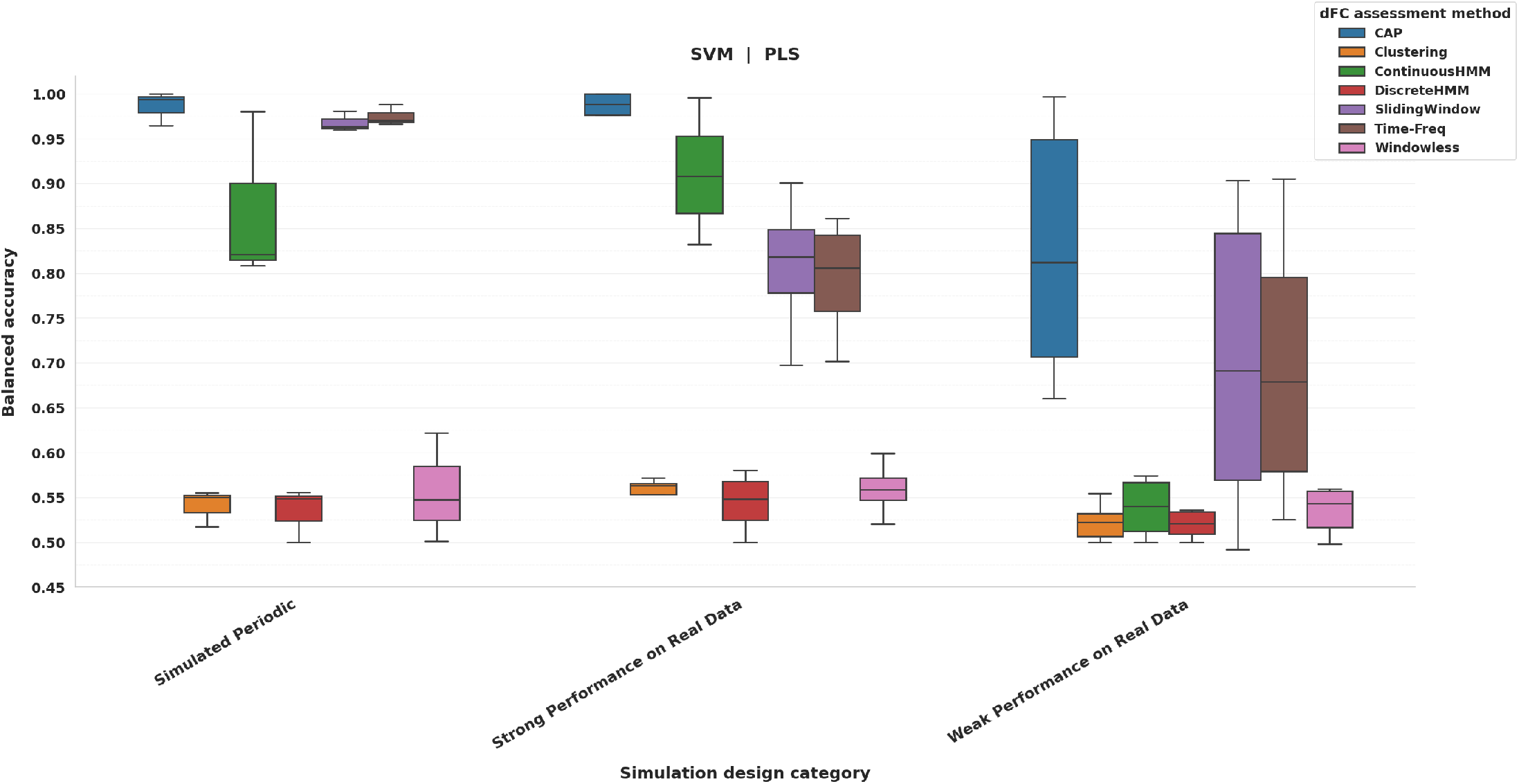
Simulated experiment categories reveal the relative contributions of experimental design, data quality, and methodology to dFC-based decoding performance. Test-set balanced accuracy for task vs. rest classification is shown across simulated experiment categories, with boxplots summarizing performance across experiments and colors indicating dFC methods. Categories include fully synthetic periodic designs (Simulated Periodic), simulations based on task timings of experiments with strong performance in experimental data, and simulations based on task timings of experiments with weak performance in experimental data. Across all methods, performance is highest for Simulated Periodic experiments, followed by simulations derived from strong-performing experiments, and lowest for those derived from weak-performing experiments. For Simulated Periodic experiments, CAP, Sliding Window, and Time-Frequency achieve near-perfect or perfect accuracy across all cases, indicating that under idealized conditions, dFC features can reliably encode task engagement and methodological limitations are not the primary constraint. Differences across categories reveal the role of experimental design. Continuous HMM shows a substantial drop in performance for simulations derived from weak-performing experiments, often approaching chance levels, suggesting that limitations in these cases are largely driven by experimental design rather than data quality. In contrast, CAP, Sliding Window, and Time-Frequency maintain relatively high performance for these same experiments compared to experimental data, indicating that while experimental design imposes constraints, reduced performance in experimental datasets is also substantially influenced by data quality. The performance achieved by Clustering, Discrete HMM, and Windowless methods remain near chance across all categories, suggesting that their limitations are primarily methodological. Continuous HMM does not reach near-perfect performance even in Simulated Periodic experiments, indicating potential methodological constraints relative to other approaches. Together, these results demonstrate that dFC decoding performance is jointly shaped by experimental design, data quality, and methodological choice, with their relative contributions varying across approaches.

Notably, Sliding Window and Time-Frequency reached nearperfect accuracy for synthetic periodic designs but not for simulations based on experimental paradigms. Because both sets of simulations share the same reduced data complexity and differ only in their task design, this contrast points to experimental design as a factor that can limit performance on its own — independent of data quality or methodological choice.

### Three Factors Predict When dFC Successfully Tracks Cognitive Engagement

Having established that dFC sometimes succeeds and often fails, we next address the following question: what specifically determines success? We conducted a top-vs-bottom profile analysis, stratifying runs of each experiment into the top and bottom 20% of decoding performance and computing standardized effect sizes (Cohen’s *d*) for a set of interpretable task design, signal-strength, and data-quality factors (Figures 5 and 6). This analysis transforms the benchmarking results into a predictive framework: a set of empirically grounded conditions under which dFC can be expected to reliably track cognitive state.

**Fig. 5.**
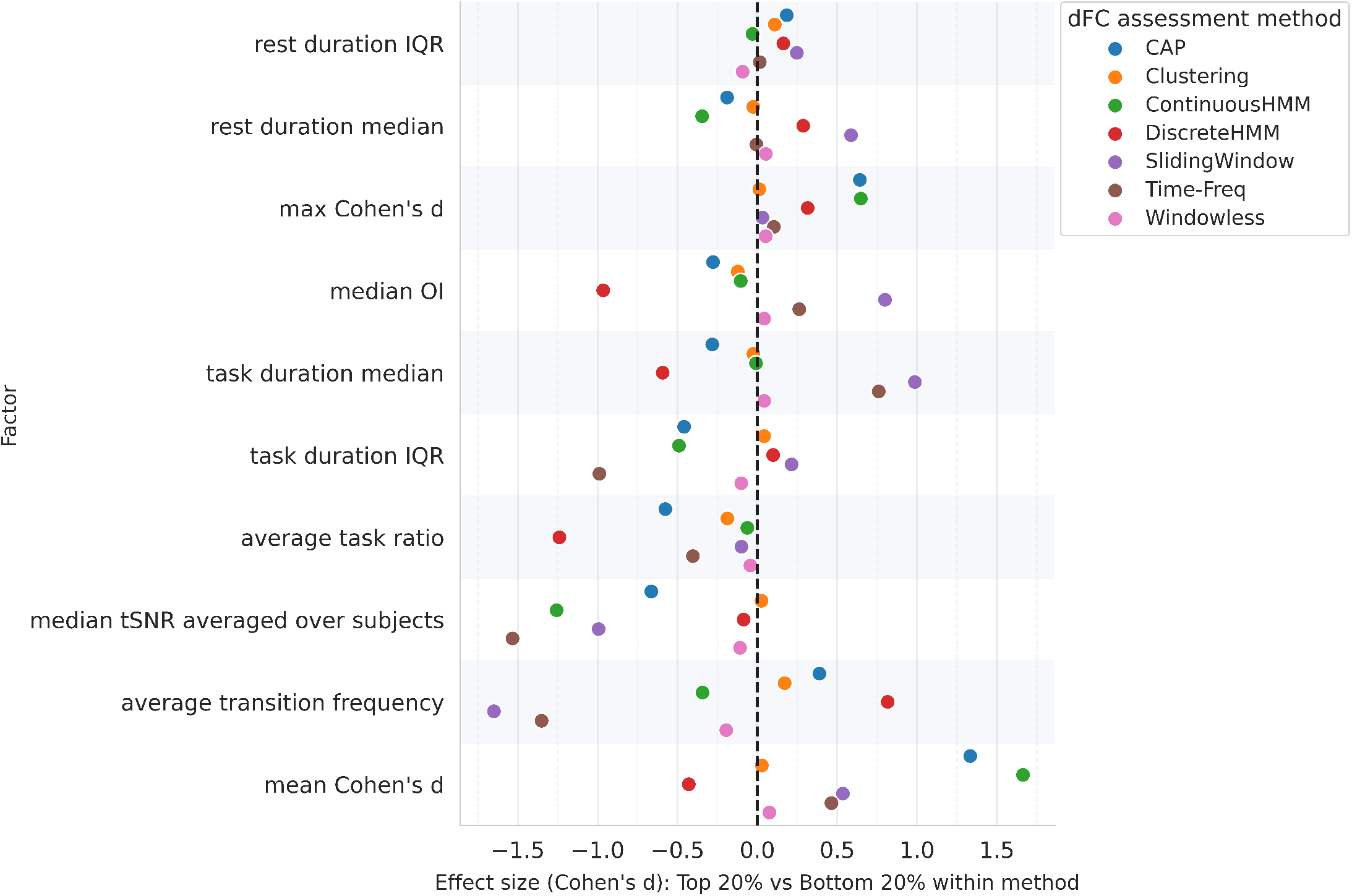
Top-vs-bottom performance profile analysis reveals “experimental task design” and “task-rest signal contrast” related factors associated with dFC decoding performance in experimental data. Within each dFC method, the top 20% and bottom 20% experiments were selected based on the achieved balanced accuracy, and standardized effect sizes (Cohen’s d) were computed for each candidate factor. Positive values indicate higher factor values in high-performing runs, while negative values indicate higher values in low-performing runs. Across methods, transition frequency exhibits a strong negative effect size, particularly for Sliding Window and Time-Frequency, indicating that frequent task-rest transitions are associated with reduced decoding performance. The mean Cohen’s d score (task-rest signal contrast) — the task-vs-rest BOLD effect size averaged across regions — shows a positive effect across methods, with stronger effects for CAP and Continuous HMM compared to Sliding Window and Time-Frequency, suggesting that larger task-rest signal contrast facilitate decoding. Task duration statistics also show method-specific effects, with longer median task durations associated with improved performance for Sliding Window and Time-Frequency, and higher variability (IQR) in task durations associated with reduced performance for Time-Frequency, CAP, and Continuous HMM. Interestingly, tSNR exhibits a negative effect size, indicating that lower tSNR is not the primary limiting factor for decoding performance. The optimality index (OI) — which compares the frequency content of the task design to that of an ideal design, with higher values indicating better-aligned, more optimal task timing [16] — shows a positive effect for Sliding Window, suggesting sensitivity to design efficiency. Given that Sliding Window and Time-Frequency exhibit the strongest and most reliable decoding performance in experimental data, their factor relationships provide the most informative characterization of performance drivers, whereas near-chance methods (Clustering, Discrete HMM, Windowless) offer limited interpretability.

**Fig. 6.**
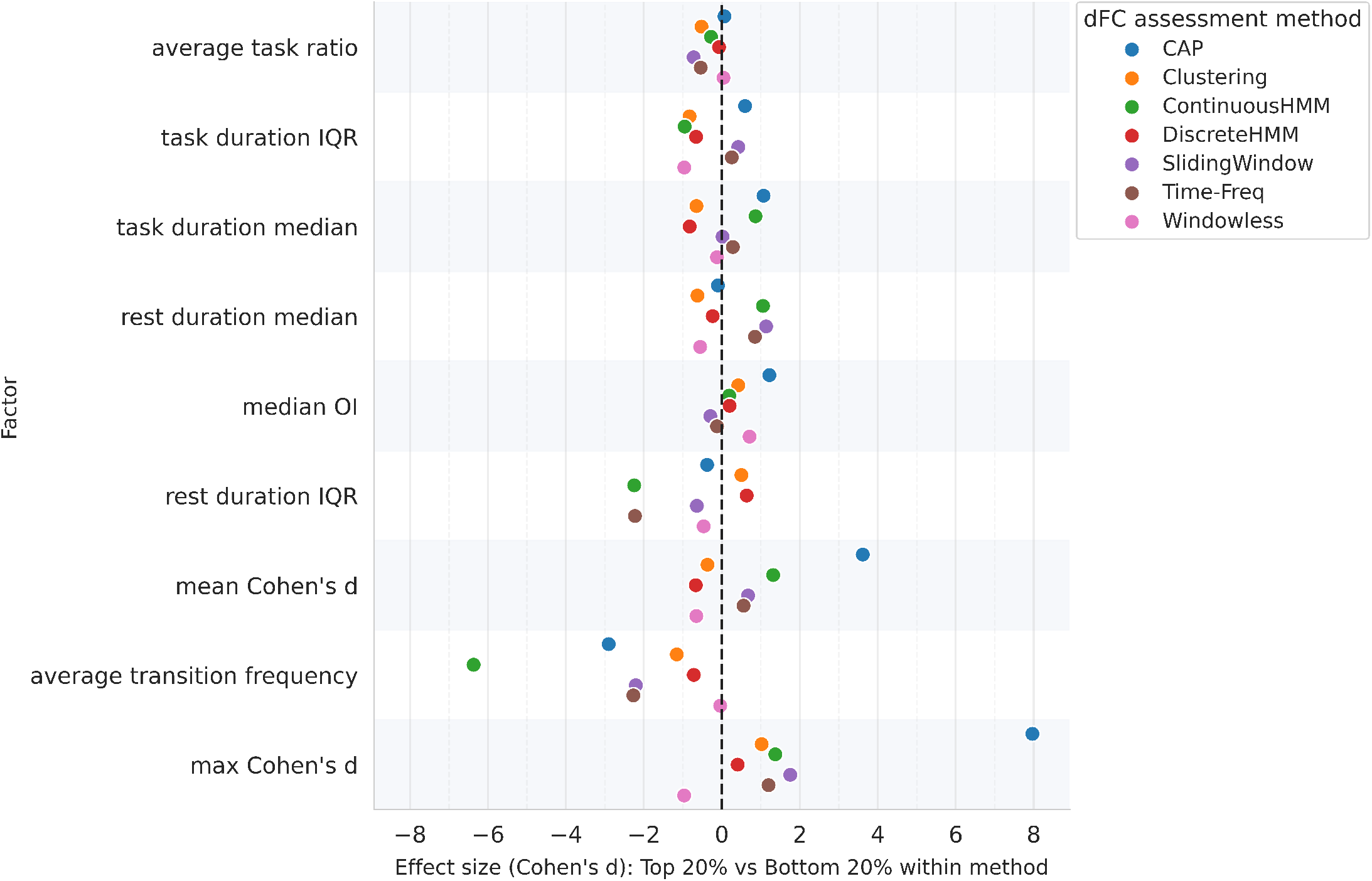
Top-vs-bottom profile analysis in simulated data highlights the dominant role of experimental design and signal strength under idealized conditions. Using the same top 20% vs. bottom 20% stratification approach as in Figure 5, effect sizes were computed for each factor across simulated runs. Transition frequency shows a consistently strong negative effect across all high-performing methods (CAP, Continuous HMM, Sliding Window, Time-Frequency), indicating that frequent task transitions strongly limit decoding performance even under idealized conditions. Mean Cohen’s d exhibits a positive effect across methods, with a markedly stronger effect for CAP compared to experimental data, suggesting increased sensitivity of state-based approaches to the task-rest signal contrast in simulated settings. Task structure variability (e.g., rest duration IQR) also shows negative effects, indicating that irregular timing reduces detectability. Methods that yield near chance performance across all conditions (Clustering, Discrete HMM, Windowless) show weak or inconsistent factor associations, supporting the interpretation that their limitations are primarily methodological rather than driven by task design or signal properties.

Three factors emerged as consistent and interpretable predictors of performance. First, **task-rest transition frequency** was the strongest and most consistent negative predictor across methods, particularly for Sliding Window and Time-

Frequency (Cohen’s *d ≈ −* 1.5 in experimental data, *d< −* 2 in simulation). Experiments in which participants frequently switched between task and rest were systematically harder to decode, likely because rapid transitions demand temporal resolution that these dFC methods cannot achieve. This effect was present even under simulated, noise-free conditions, confirming that transition frequency imposes a design-level ceiling on cognitive state tracking that cannot be overcome by methodological refinement alone.

Second, **task-rest signal contrast** (Cohen’s *d* of BOLD differences between task and rest averaged across regions) was a consistent positive predictor: experiments with larger neural response differences were more decodable. This effect was stronger for state-based methods (CAP, Continuous HMM) than for state-free approaches, suggesting that methods that model discrete brain states may be especially sensitive to the magnitude of task-evoked responses.

Third, **task block structure** — specifically the length and regularity of task epochs — was associated with performance in a method-dependent manner. Longer median task durations facilitated decoding for Sliding Window and TimeFrequency, while greater variability in block durations impaired performance for Time-Frequency, CAP, and Continuous HMM. These findings indicate that the temporal regularity of cognitive demands, not just their presence, shapes whether dFC can faithfully decode them.

Crucially, the same factors predicted performance in both experimental and simulated data, but their effects were stronger in simulation. Transition frequency remained the dominant negative predictor, and the task-rest signal contrast effects became more pronounced — particularly for CAP — in the absence of real-world artefacts such as motion and physiological noise. This consistency across conditions supports these factors as robust design-level determinants of cognitive state trackability, rather than features specific to either the noisy experimental data or the controlled simulated setting.

Methods that remained near chance across all conditions (Clustering, Discrete HMM, Windowless) showed weak and inconsistent factor associations, consistent with the interpretation that their limitations are methodological rather than driven by task design or signal properties.

Together, these three factors — transition frequency, task-rest signal contrast, and block regularity — constitute a practical and empirically validated framework for predicting when dFC will more likely reliably track ongoing cognition.

Although not explicitly captured by these factors, the cognitive domain of an experiment may modulate performance indirectly through its influence on task structure or signal amplitude. A preliminary analysis across cognitive domains is provided in Supplementary materials Section, but does not support strong domain-specific conclusions at this stage.

### Cognitive States Are Weakly Separated in dFC Feature Space

To complement the supervised decoding analyses, we assessed the intrinsic structure of dFC feature spaces using the Silhouette Index (SI) — a measure of how naturally task and rest time points cluster without any classifier. Across experiments from experimental datasets and all dFC methods, SI values were consistently low, with most combinations falling between 0 and 0.2 (Figure 7). Task-present and rest states do not form naturally separable clusters in the connectivity feature space as currently defined by these methods.

**Fig. 7.**
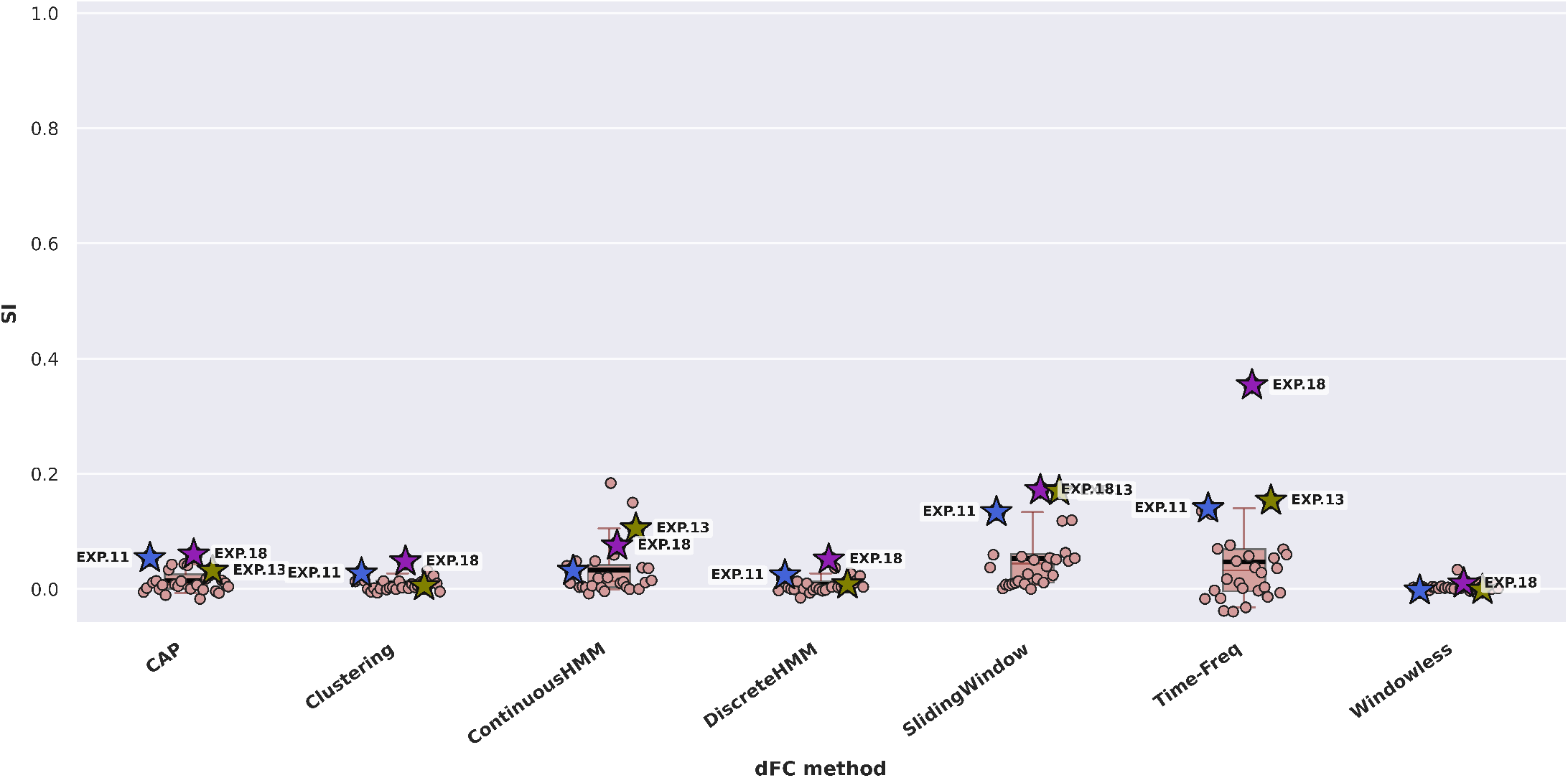
Intrinsic task-rest separability across dFC methods in experimental datasets, quantified by the Silhouette Index (SI). Each boxplot shows the distribution of SI scores across experiments (`EXP.1`-`EXP.28`) for a given dFC method, with median (red line) and mean (black line) values indicated. Individual points represent experiments, with selected experiments exhibiting higher separability (`EXP.11`, `EXP.13`, `EXP.18`) marked with a star. Across methods, SI values are generally low, with most experiment-method combinations falling between 0 and 0.2, indicating weak intrinsic separability between task and rest states in the dFC feature space. The highest separability is observed for `EXP.18` using Time-Frequency (SI = 0.35), with a handful of additional experiments (e.g., `EXP.11`, `EXP.12`, `EXP.13`, `EXP.27`, `EXP.28`) reaching modest values around 0.13-0.18 depending on the method. Sliding Window and Time-Frequency exhibit a slightly higher central tendency (mean SI *≥* 0.1) compared to other methods, although separability remains limited overall. Continuous HMM shows moderate separability in a subset of experiments, while CAP, Clustering, Discrete HMM, and Windowless methods remain near zero across most cases. Overall, these results indicate that rest and task-present states are only weakly separated in the dFC feature space in experimental data, and that reliable cognitive state decoding requires supervised rather than unsupervised analytic strategies. For exact SI values across all experiment-method pairs, see Supplementary Figure 53, and for detailed across-run variability, see Supplementary Figure 54.

Only isolated experiments showed modest separability. The highest value observed was SI = 0.35, for `EXP.18` using Time-Frequency, with a handful of additional experiments (`EXP.11`, `EXP.12`, `EXP.13`, `EXP.27`, `EXP.28`) reach-ing SI values of 0.13–0.18 depending on the method. Sliding Window and Time-Frequency exhibited slightly higher average separability than other methods.

The observed low separability may be viewed as being at odds with the above-chance classification performance observed for several experiment–method combinations (e.g. top performers such as `EXP.12`, `EXP.13`, and `EXP.14`). This apparent discrepancy likely arises because task-related information is present in the dFC features but is weak and spread across many dimensions, rather than concentrated in a way that separates task and rest into distinct clusters. Supervised models, which are guided by the known labels, can pick up these faint and distributed differences, whereas unsupervised clustering — with no access to the labels — cannot. This has practical implications for how dFC should be used as a cognitive state tracker — methods that rely on unsupervised state identification (as many standard dFC pipelines do) may systematically miss task-related dynamics that are in principle detectable. Reliable tracking of cognitive engagement from dFC likely requires supervised or hypothesis-driven analytic approaches.

*Note: For experiments with multiple runs, all reported values correspond to the best-performing run. Supplementary figures provide the range (min-max) across all runs for completeness*.

## Discussion

A central aspiration of cognitive neuroscience is to decode the dynamics of cognition using brain dynamics. DFC has attracted wide interest as a tool for achieving this goal by tracking the moment-to-moment reconfiguration of brain networks as cognitive states unfold in real time. However, the empirical foundation for this ambition is stil not sufficiently elucidated. Our results provide the most systematic evaluation to date of whether dFC can achieve this overall goal, and under which conditions.

Overall, our results strongly suggest that dFC-based tracking of cognitive engagement is unreliable in many experimental contexts — but this failure does not appear to be solely a limitation of the dFC approach itself. Across both experimental and simulated data, we observed that decoding performance varied systematically with the experimental design, the data quality, and the dFC method used: some combinations yielded reliable task decoding, while many others did not. This suggests that the unreliability seen in experimental data also depends on the conditions under which dFC is applied. The key question is therefore not simply whether dFC can track cognition, but under what conditions it is more likely to do so — and our results provide a principled, empirically grounded answer structured around three interacting factors that shape success: **experimental design, data quality**, and **methodological choice**. Within experimental design, three specific features are particularly consequential: task-rest transition frequency, task-rest signal contrast, and task block regularity.

### The Gap Between Simulated and Experimental Performance

One of the most striking findings of the present study is the contrast between simulation and experimental performance. When simulation data were used, several methods achieved near-perfect single-TR decoding of task engagement; in experimental data, many experiment–method combinations did not exceed chance levels. This gap helps identify where the real difficulty lies: even if the information needed to track cognitive engagement is captured by connectivity dynamics, our results suggest it can be systematically obscured in experimental data by the complex, aggregate nature of the BOLD fMRI signal (motion and physiological artefacts, neurovascular coupling) and by suboptimal experimental design.

This distinction has implications regarding how we may interpret the results of dFC studies. If the drivers identified here generalize, they would predict that published findings of meaningful dFC dynamics during cognition are more likely to emerge from studies with favorable experimental designs and high data quality, while null results may disproportionately reflect unfavorable conditions rather than the absence of real cognitive dynamics. We have not verified this directly across the existing literature as it is outside the scope of the current study, and doing so would require the kind of systematic meta-analytic coding we identify as a future direction. Nevertheless, our framework provides a principled basis for adjudicating between these possibilities post hoc when evaluating individual studies, and for designing future studies that optimize the conditions under which dFC can reliably detect cognitive dynamics.

### Experimental Design as the Primary Determinant

Of the three main drivers, experimental design exerts the most fundamental influence on dFC performance — fundamental in the sense that its effects persist even under noise-free simulated conditions, beyond the reach of methodological refinement or data-quality improvements. Our factor analysis identifies three specific design features that jointly determine how detectable task engagement will be.

**Task-rest transition frequency** is the dominant limiting factor. Frequent transitions between cognitive engagement and rest reduce decoding performance strongly and consistently across methods, and this effect persists even under noise-free simulated conditions. This indicates a fundamental temporal resolution constraint: current dFC methods — even the best-performing ones — cannot faithfully track cognitive state when the brain is asked to switch rapidly between engagement and rest. This constraint is design-level, not methodological, and cannot be overcome by choosing a better algorithm. It implies that event-related and rapid mixed designs, which are standard in much of cognitive neuroscience, are intrinsically poorly suited for dFC-based state tracking, regardless of the method applied.

**Task-rest signal contrast** is the second key design-level determinant. Experiments in which task engagement produces a larger BOLD contrast relative to rest are systematically more decodable. This effect is stronger for state-based methods (CAP, Continuous HMM) compared to state-free approaches, suggesting that methods which model discrete brain states may be particularly sensitive to the magnitude of task-evoked network reconfigurations. From a cognitive neuroscience perspective, this suggests that dFC is more likely to track cognitive engagement when this engagement produces a substantial and sustained reorganization of network architecture — a property that may vary across cognitive domains and task demands.

**Task block regularity** is the third design-level factor. Longer and more temporally regular task epochs facilitate decoding, while irregular or variable block structures impair it. This is consistent with the temporal smoothing inherent in most dFC estimation approaches: methods that integrate connectivity information over a window or state assignment will naturally perform better when the cognitive signal is sustained and predictable. This also helps explain why earlier studies that classified entire sustained task blocks — rather than individual time points — achieved substantially higher performance [8, 11]: long, stable blocks represent the most favorable end of this design dimension, whereas single-TR decoding across shorter and more variable epochs is considerably more demanding. The implication is that experimental designs optimized for mass-univariate GLM analyses — which benefit from variance in timing — are not necessarily optimal for dFC analyses, and that dFC-focused studies may require different design principles.

Together, these three design features constitute a practical checklist for prospective study design: paradigms with low transition frequency, high task-evoked contrast, and regular block structure are systematically more amenable to dFC-based cognitive state tracking.

### Data Quality as a Modulating Factor

Beyond experimental design, the quality of the data plays an independent and substantial role. By data quality, we mean the extent to which the measured BOLD signal faithfully reflects the underlying task-related neural dynamics — which is degraded by motion, physiological fluctuations, and the indirect, aggregate nature of the BOLD signal. The large performance gap between simulated and experimental data — present even for experiments with favorable designs — indicates that these factors attenuate task-related dynamics in ways that experimental design alone cannot compensate for. Many of these factors are intrinsic to fMRI and only partly controllable, which sets a practical ceiling on what design optimization alone can achieve.

Importantly, the contributions of experimental design and data quality are separable in our framework. Simulations that preserve task timings of experimental data, but eliminate the influence of factors such as motion and physiological artefacts, allow us to isolate what design alone constraints; com-parisons between simulated and experimental performance for the same experiment reveal the additional attenuation attributable to data quality. This decomposition suggests that for some experiments, design is the primary bottleneck; for others, data quality is the dominant limitation. Both must be considered when interpreting why a given study succeeded or failed in terms of using dFC to track cognitive engagement.

### Methodological Choice Determines What Can Be Extracted

Given favorable experimental design and sufficient data quality, methodological choice determines whether the available task-related signal can actually be detected. Across all conditions, methods fell into two clearly distinct groups. Sliding Window and Time-Frequency methods showed the most consistent sensitivity to task-related dynamics in experimental data, with some state-based approaches (CAP, Continuous HMM) exhibiting strong performance under idealized conditions but greater sensitivity to data quality. In contrast, Clustering, Discrete HMM, and Windowless methods achieved near chance performance across virtually all conditions — including simulated data — indicating that their limitations are primarily methodological and cannot be attributed to design or data quality.

This methodological stratification has direct implications for the field. Encouragingly, Sliding Window — among the most widely used dFC methods [10] — was also among the better-performing methods for decoding task engagement in our benchmark, suggesting that the field’s most popular choice performed comparatively well under the conditions tested here. This alignment was not consistent, however: Clustering, also widely used, showed limited validity within our framework, while Time-Frequency performed comparably to Sliding Window; yet, it remains substantially underrepresented in the literature. These discrepancies suggest that current practice is not always well aligned with the available evidence — a gap that systematic benchmarking like ours aims to address.

An important nuance is that methods are not consistent in terms of which experiments are most tractable. This method– experiment interaction implies that different dFC approaches are sensitive to distinct properties of the underlying signal, and should be treated as complementary rather than interchangeable. The choice of method is not merely a technical preference — it encodes assumptions about the temporal and spatial structure of cognitive dynamics that may or may not match the paradigm at hand.

### State-Based and State-Free Methods Represent Subject Variability Differently

A structural difference between the two method families warrants consideration when interpreting their relative performance. State-free methods (Sliding Window, Time-Frequency) retain subject-specific connectivity information at every time point, whereas state-based methods (CAP, Clustering, Continuous and Discrete HMM, Windowless) summarize connectivity using a fixed set of group-level states — here, five — shared across all subjects. This discretization necessarily discards some of the subject-specific variability that state-free methods preserve, and this variability may carry information relevant to tracking ongoing cognition. To mitigate this, we did not classify directly on the discretized group-level patterns but instead used the time-resolved, subject-specific state-related dFC quantities (state probabilities, distances to state centroids, or state weights), which retain how each individual engages the shared state space over time.

This representational difference is not in itself a limitation of either family, but a consequence of their design. The discretization performed by state-based methods is deliberate: by reducing connectivity to a small set of recurring states, these methods are intended to be more robust to noise, and, ideally, the recovered states would correspond to distinct, cognitively meaningful configurations that track engagement and rest. Our results suggest that, under the conditions tested here, this is largely not the case: in experimental data, the state-based methods did not reliably distinguish task from rest even when subject-specific state expression was used as the feature. CAP and Continuous HMM did show improved performance under simulated conditions, indicating that their discretized representations can encode task engagement when data quality and design are favorable, but this improvement was not observed in experimental data. The low intrinsic separability observed across methods (Figure 7) is consistent with this interpretation: the group-level states recovered by these methods do not appear to align cleanly with the task–rest distinction. Whether the discretized representations used by state-based methods could be configured to better capture cognitively relevant dynamics remains an open question that our study does not address.

### Cognitive Information Is Latent, Not Geometrically Compact

The consistently low Silhouette Index values across experiments reveal that task and rest states do not form naturally separable clusters in dFC feature space. However, supervised classifiers can still achieve meaningful above-chance decoding in a subset of experiments. This apparent contradiction resolves to an important insight: task-related information is present in dFC features, but it is distributed and latent rather than geometrically compact. It can be detected by hypothesis-driven supervised approaches but not by the unsupervised clustering methods.

This has a direct practical implication that spans all three drivers. Even when experimental design and data quality are favorable and a sensitive dFC assessment method is chosen, studies that rely on unsupervised state identification for linking dFC patterns to cognition may systematically fail to detect the signal. The states identified by unsupervised methods are unlikely to align with cognitively meaningful transitions because that information is not expressed as compact, separable clusters. Reliable cognitive state tracking from dFC therefore requires not only the right design, data quality, and method, but also a supervised or hypothesis-driven analytic strategy that can recover distributed patterns.

### A Framework for Contextualizing dFC Results

The three main drivers identified here — experimental design, data quality, and methodological choice — are rarely reported systematically in published dFC studies, yet our data show they predict a substantial portion of the variance in decoding performance across 28 independent experiments. If these drivers generalize beyond our benchmark dataset, they would predict the following specific pattern: studies that combine favorable experimental designs (low transition frequency, high signal amplitude, regular block structure) with good data quality and sensitive dFC assessment methods would succeed in using dFC to track cognition, while those that do not fulfill these conditions would not. Whether this prediction holds across the broader literature remains to be tested — doing so would require a systematic meta-analysis that codes published studies along these dimensions and asks whether performance outcomes co-vary accordingly. We view this as an important direction for future work.

In the meantime, our results provide a practical basis for contextualization: Rather than asking “does dFC track cognition?” as if it were a binary question with a universal answer, studies should report the design and data-quality factors identified here to help answer the more tractable question of *when* dFC tracks cognition, allowing readers to situate results within the broader performance landscape that our benchmark defines. Our open-source `PydFC` toolbox [12] provides a comprehensive and reproducible framework for computing, comparing, and benchmarking dFC methods across datasets — equipping researchers with the tools needed to characterize the design and data-quality factors that our results identify as critical.

### Limitations and Future Directions

Several limitations of the current study point toward important directions for future work. Although our factor analysis identifies variables that co-vary with performance, it does not establish causal relationships among the three main drivers. Controlled experimental manipulations that independently vary transition frequency, block length, signal contrast, and noise characteristics within the same paradigm would allow more direct causal inference about their relative contributions.

Our conclusions about data quality also rest on a generative model that, by design, is simpler than real-world fMRI. The simulation reproduces task timing and introduces intersubject variability, but it does not reproduce the specific sources of signal degradation that affect experimental data — such as head motion, respiratory and cardiac fluctuations, fluctuations in arousal, and variability in the hemodynamic response across regions and individuals [15, 17, 18]. This makes the simulation a useful controlled reference, but it also suggests that it treats these degrading factors as a single block: it shows what happens when they are all removed, without resolving how much each one individually contributes. A natural extension would be to incorporate specific data-quality factors directly into the simulation — for example, by adding controlled levels of motion, physiological noise, or hemodynamic variability — and observing how each degrades decoding performance [14, 19]. This would move the design–data-quality decomposition from a comparison between two conditions toward a controlled, parametric account of how individual sources of signal degradation limit dFC-based cognitive tracking.

A further limitation concerns the role of cognitive domain. Our cognitive domain analyses (Supplementary materials) suggest that some task categories may be associated with better performance, but these patterns are confounded by design differences across paradigms — disentangling cognitive domain from experimental design requires purpose-built studies in which cognitive demands vary while design parameters are held constant.

Finally, and most fundamentally, the present work tests only the binary distinction between task engagement and rest. This is a deliberately stringent validity criterion, but it is only a first step: extending the framework to finer-grained cognitive distinctions — between different task types, or different levels of engagement — will test whether the conditions identified here generalize to the more nuanced cognitive tracking that dFC is ultimately meant to support.

### Conclusion

Dynamic functional connectivity may be used to track ongoing cognitive engagement, but its success depends on three interacting drivers: experimental design, data quality, and methodological choice. Of these, experimental design is the most fundamental — it sets a performance ceiling that neither better methods nor higher-quality data can overcome. Within design, three features are critical: low task-rest transition frequency, strong task-rest signal contrast, and regular block structure. Data quality determines how much of that design-level potential is expressed in the signal; methodological choice determines how much of the expressed signal can be detected. When all three drivers are favorable, dFC is most likely to track cognitive engagement reliably at single-TR resolution. When they are not — and in most real-world fMRI studies they are not simultaneously optimized — dFC fails, not necessarily because of an intrinsic limitation of the approach, but often because the conditions for success have not been met. Our work reframes dFC not as a universally valid or universally limited tool, but as a context-sensitive measure whose reliability is predictable. For cognitive neuroscience, this suggests that designing studies to reliably track cognitive dynamics with dFC requires treating experimental design, data quality, and methodological choice as jointly optimized components of a coherent analytic strategy.

## Methods and Materials

We developed a unified framework to evaluate and interpret dFC methods for predicting task presence at the level of individual time points. The pipeline operates on BIDS-formatted fMRI datasets with associated task timing information and supports multiple datasets and experimental paradigms, enabling large-scale benchmarking. Time-resolved dFC features were extracted from preprocessed BOLD fMRI signals and used to classify task presence (task vs. rest) at each time point. Performance was assessed using balanced accuracy across methods and experiments.

### fMRI Datasets Selected from OpenNeuro

Functional MRI data were obtained from the OpenNeuro repository. To ensure suitable data quality and coverage for our study, we applied the following inclusion criteria: (i) the dataset contains functional scans (`func` datatype); (ii) includes at least 50 participants; (iii) has at least one block-designed task paradigm with sufficiently long task and rest periods; (iv) is BIDS-compliant; and (v) includes `events.tsv` files with task/stimulus timing information. Out of the entire repository, only 22 datasets met these criteria, yielding a total of approximately 50 experiments. After extracting task presence and rest/task labels, experiments with sub-optimal design those producing extreme class imbalances or unreasonable task presence time courses - were excluded. The final selection included 16 datasets comprising 28 experiments. A detailed overview of these datasets, including number of subjects and experiments, is provided in Table 1.

**Table 1.**
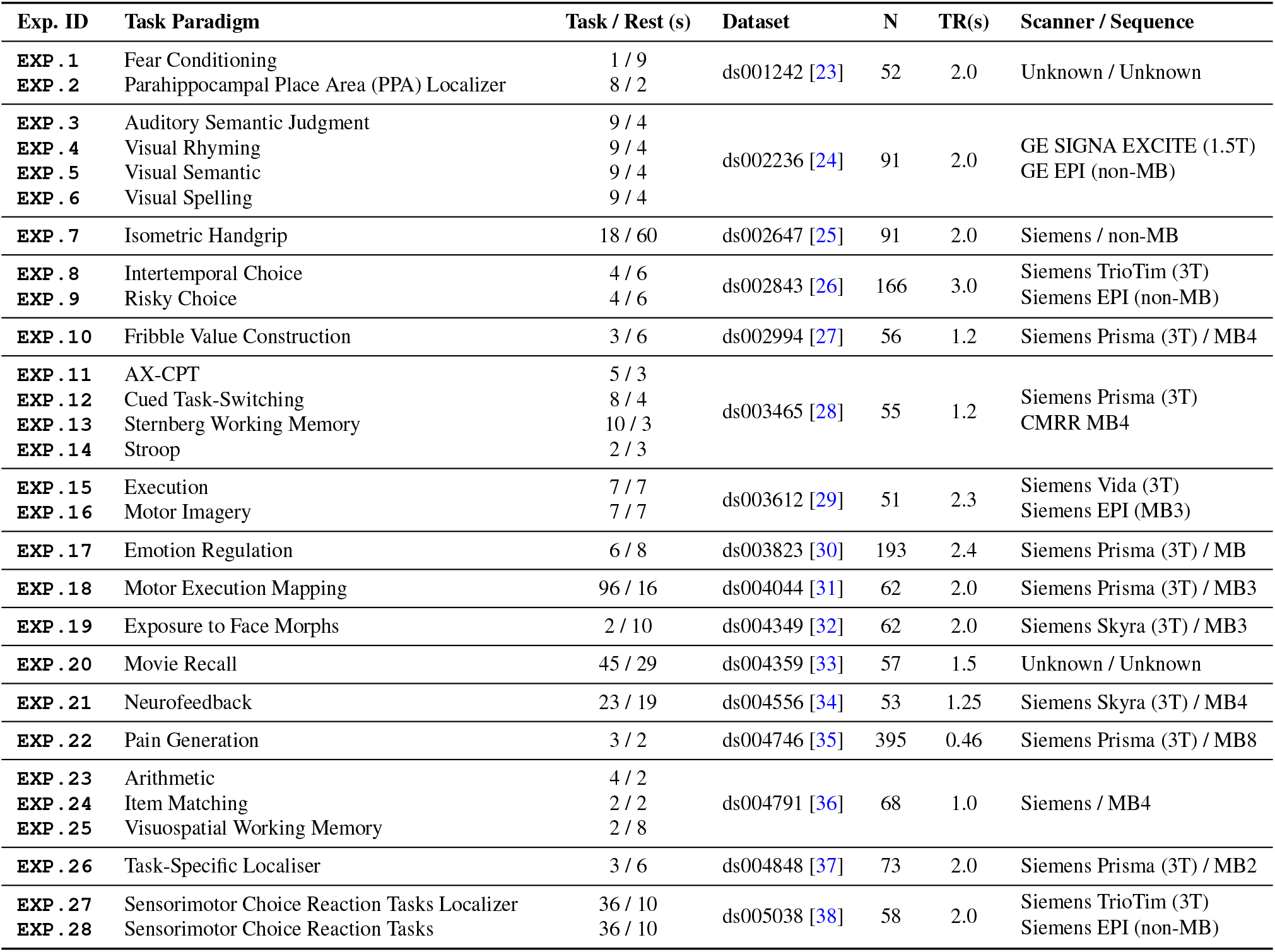
Characteristics of the 28 Experimental Settings. The table details the 28 distinct experimental settings (IDs `EXP.1` - `EXP.28`) and their corresponding task paradigms. For each experiment, the respective OpenNeuro dataset and shared acquisition parameters (N subjects, TR, Scanner, Sequence) are given. Abbreviations: MB = Multiband; TR = Repetition Time

All datasets were processed starting from raw fMRI `NIfTI` images. First, we applied `fMRIprep` through `nipoppy`, a lightweight framework that streamlines preprocessing for large, heterogeneous neuroimaging datasets by managing parameters via `JSON` configuration files. `fMRIprep` applied standard minimal preprocessing, including motion correction, slice-timing correction, and spatial normalization. The confound variables provided by `fMRIprep` were used to further denoise each dataset using `nilearn’s` simple denoising strategy, which includes high-pass filtering, tissue signal regression (CSF and white matter), full motion correction, and demeaning [20].

Following denoising, voxel-wise BOLD signals were parcellated using the Schaefer 100-region atlas [21], yielding regional time series for each experiment. These regional time series formed the basis for all subsequent dFC analyses.

### Task domains and codes

We assigned each task paradigm to a *Research Domain Criteria* (RDoC) [22] *domain* and to its corresponding *construct* (e.g., Cognitive Systems → Working Memory; Positive Valence Systems → Reward Valuation). Our primary domains were: *Cognitive Systems* (language, cognitive control, working memory, attention, perception), *Positive Valence Systems* (reward valuation, reward responsiveness), *Negative Valence Systems* (acute threat (“fear”)), *Arousal and Regulatory Systems* (arousal), and *Sensorimotor Systems* (motor actions) (see Table2). This RDoC-aligned categorization clarifies the targeted processes and supports principled comparisons across heterogeneous tasks.

**Table 2.** Task domains and associated paradigms based on Research Domain Criteria (RDoC).

| Domain | Task Paradigm |
| --- | --- |
| Arousal and Regulatory Systems | Emotion Regulation |
| Cognitive Systems | Arithmetic, AX-CPT, Auditory Semantic Judgment, Cued Task-Switching, Exposure to Face Morphs, Neurofeedback, Task-Specific Localiser, Sensorimotor Choice Reaction Tasks Localizer, Item Matching, Parahippocampal Place Area (PPA) Localizer, Movie Recall, Sensorimotor Choice Reaction Tasks, Sternberg Working Memory, Stroop, Visual Rhyming, Visual Semantic , Visual Spelling , Visuospatial Working Memory |
| Negative Valence System | Fear Conditioning, Pain Generation |
| Positive Valence System | Fribble Value Construction, Intertemporal Choice, Risky Choice |
| Sensorimotor Systems | Execution, Isometric Handgrip, Motor Imagery, Motor Execution Mapping |

### Simulated Experiments

To complement experimental fMRI datasets, we generated simulated data using The Virtual Brain (TVB) framework [39], which allows flexible modeling of neural dynamics and network connectivity. TVB enabled us to define task timing and systematically vary key parameters across subjects, including structural connectivity, the global coupling coefficient (which modulates functional connectivity strength between regions), and connection speed.

We generated a total of 13 simulated experiments, organized into three categories of increasing expected difficulty. The first consists of idealized periodic designs, which have the regular, predictable structure that should be easiest to decode. The second and third reproduce the task timing of real-world experiments — those that performed well in our experimental data analyses, and those that performed poorly and are expected to be the hardest to decode:

1. **Simulated Periodic** fully synthetic periodic designs, including: *task-lowFreqLongRest, task-lowFreqShortRest*, and *task-lowFreqShortTask* (`EXP.S.29`, `EXP.S.30`, and `EXP.S.31`). These experiments include 200 subjects, a TR of 500 ms, and a scan length of 250 s. They vary systematically in task/rest durations:
  1. *task-lowFreqLongRest*: task = 8 s, rest = 12 s
  2. *task-lowFreqShortRest*: task = 12 s, rest = 8 s
  3. *task-lowFreqShortTask*: task = 1 s, rest = 19 s
2. **Strong Performance on Experimental Data** simulations based on task timings from experimental datasets that yielded better performance in our analyses, including `EXP.11`, `EXP.12`, `EXP.13`, and `EXP.14` (from `ds003465`), named `EXP.S.11` - `EXP.S.14`, same numbers as their experimental counterparts.
3. **Weak Performance on Experimental Data** - simulations based on task timings of experiments yielding worse performance in experimental data, including `EXP.2`, `EXP.8`, `EXP.9`, `EXP.15`, `EXP.16`, and `EXP.27` (from ds001242, ds002843, ds003612, and ds005038), named `EXP.S.2`, `8`, `9`, `15`, `16`, and `27`, same numbers as their experimental counterparts.

Simulated experiments derived from experimental datasets matched the original dataset in terms of number of subjects, TR, scan length, and task timing. These simulations served as a reference for performance under idealized conditions, enabling direct comparison with corresponding real-world experiments.

All simulated data were generated using TVB’s default structural connectivity, corresponding to a macaque brain with 76 regions. During task-on blocks, we stimulated five regions (0, 7, 13, 33, 42). To introduce variability across subjects, we slightly randomized the structural connectivity, global coupling coefficient, and connection speeds by adding Gaussian noise to each parameter. Neural dynamics were integrated using the stochastic Heun method with additive Gaussian noise (dispersion D = 0.1). As the appropriate noise level in TVB is model-dependent and not fixed by a single standard value, we set D = 0.1 based on visual inspection of the simulated BOLD signals, and dFC patterns across all examined methods, selecting a value that produced realistic dynamics without making the simulated signals unrealistically simple.

### Deriving Task Presence Time Courses

For each experiment, task presence was determined separately for each individual scan (run), ensuring subject- and run-specific labeling. Initial labels were assigned using the onset and duration values provided in the `events.tsv` files: time points were marked as task-present (1) or rest (0). To account for the hemodynamic delay in fMRI signals due to neurovascular coupling mechanisms, the binary time course was convolved with a canonical HRF implemented using `SPM’s` double-gamma function implemented in the `nilearn` toolbox. The resulting HRF-convolved time course, sometimes termed a block regressor [7] or expected task response, was then down-sampled to the MRI TR and binarized again to produce the final task presence time course.

Binarization was performed using a Gaussian Mixture Model (GMM), which assigns probabilities to each time point belonging to each cluster, i.e., rest or task. This probabilistic framework allows identification and removal of ambiguous “gray zone” time points corresponding to transitions between rest and task. Excluding these uncertain samples improves classifier performance and avoids confusing the model with unclear labels.

The GMM-based procedure was implemented as follows:

1. **Initial clustering:** Fit a two-component GMM to all time points.
2. **Downsampling:** Cluster probabilities were downsampled to MRI TR resolution. For sliding window dFC methods, probabilities were further aligned to the time points of the resulting dFC matrices.
3. **Discarding ambiguous points:** Time points with probabilities < 0.99 for both clusters were removed. This threshold worked well for most experiments. In cases where a class became empty, the threshold was iteratively increased until both classes contained at least one sample. For a small number of experiments, even this adjustment did not resolve the issue; visual inspection revealed sub-optimal or unclear block designs that made reliable task/rest assignment impossible. These experiments (*N* = 6) were excluded at this stage and are not among the 28 experiments reported in this study.
4. **Handling transitional clusters:** If the two clusters did not clearly correspond to rest and task (e.g., the higher-mean cluster < 0.75 or the lower-mean cluster > 0.25), the GMM was refitted with three clusters, with the middle cluster representing transition periods.

Finally, all task presence time courses were visually inspected. Experiments with unclear rest/task assignments or extreme class imbalance were discarded from further analyses. Illustrations of the GMM-based clustering and label assignment are shown in the Task Timing stage of Figure1, as well as in Figures 14 and 15 (Supplementary materials), including examples of experiments with optimal and sub-optimal design.

### Dynamic Functional Connectivity Assessment

DFC was computed using seven widely used methodologies, grouped into state-free and state-based approaches:

- **State-free:** Sliding Window [40] and Time-Frequency [3].
- **State-based:** Co-Activation Patterns (CAP) [41], Clustering [1], Continuous Hidden Markov Models (HMM) [42], Discrete HMM [43], and Windowless [44].

All methods were implemented using the `PydFC` toolbox [12]. Detailed implementation, assumptions, and hyperparameters are described in [10]. Minor adjustments applied here include: setting the overlap ratio for sliding window methods to 1.0 (full overlap minus one sample) to maximize temporal coverage, and changing the clustering distance metric for clustering and discrete HMM from Euclidean to Manhattan. The number of brain states for all state-based methods was set to five. The reason for choosing five was both for computational reasons, and also due to the assumption that the brain during a task based scan transitions between more limited number of states. Supplementary Figures 25 and 26 show representative dFC patterns obtained with each method across two example experiments.

### Feature Extraction and Dimensionality Reduction

After computing dFC, features were extracted for classification of task presence, with state-free and state-based methods handled separately.

#### State-free methods

Time-resolved dFC matrices (time × region × region) were vectorized into edge-wise feature vectors (time × connection, with 100 (100 − 1)*/*2 = 4950 connections).

Samples corresponding to transition periods identified by the task presence procedure were excluded. To reduce the effects of inter-subject variability, features were then centered for each subject by subtracting the subject-specific mean. This step removes subject-specific offsets while preserving within-subject dynamics. Following subject centering, features were standardized using `StandardScaler` (zero mean, unit variance), fitted on training data and applied to validation and test sets when performing cross-validation. Dimensionality reduction was then performed using supervised Partial Least Squares (PLS), which learns latent components that maximize covariance between dFC features and task labels [45]. PLS was applied within a cross-validated pipeline, where the number of components was treated as a hyperparameter and selected using group-aware cross-validation on the training data. Candidate component numbers ranged from low-dimensional (e.g., 2-5) to moderately high-dimensional (e.g., up to 50), and selection was based on balanced accuracy. All transformations were fitted on training folds only and applied to validation and test data to prevent information leakage.

As a comparison, Principal Component Analysis (PCA) was also implemented, with the number of components selected via cross-validation; PCA results are reported in the Supplementary materials.

#### State-based methods

For state-based approaches (CAP, Clustering, Continuous/Discrete HMM, Windowless), features were derived from the temporal expression of connectivity states rather than full edge-wise matrices. This is because, in these methods, the connectivity matrices correspond to a fixed set of states estimated at the group level and shared across time points; the time-resolved, subject-specific information instead resides in how each time point expresses those states. Accordingly, we extracted state probabilities (HMM-based methods), distances to state centroids (CAP/Clustering), or state weights (Windowless), resulting in feature matrices of shape (time,*K*), where *K* is the number of states.

Because these features are compositional (non-negative and summing to one), we applied an isometric log-ratio (ILR) transformation, mapping them to Euclidean space with dimensionality *K −*1. Samples corresponding to transition periods were removed prior to analysis. Subject centering and standard scaling were then applied as described above. The resulting normalized ILR-transformed features were used directly without additional PCA or PLS, as their dimensionality is already low.

For each experiment and dFC method combination, features from all subjects were concatenated into a final matrix of shape (subject × time, features). These concatenated, normalized, and transformed features serve as the inputs for task presence prediction stage.

#### Remarks on manifold learning approaches

We also evaluated Laplacian Eigenmaps (LE) with Procrustes alignment, following prior work [46]. While LE can provide meaningful low-dimensional representations at the individual-subject level, its embeddings are not uniquely defined and are subject to arbitrary orthogonal transformations. Aligning these embeddings across subjects using Procrustes analysis requires an implicit assumption of point-wise correspondence between time points, which does not generally hold in dFC data.

In experiments with identical task timing across subjects, Procrustes alignment can enforce correspondence between samples that share the same temporal position and label, introducing a form of information leakage and artificially inflating classification performance. In more realistic settings with variable timing across subjects, this assumption breaks down, and alignment becomes unreliable or requires discarding large portions of the data.

Due to these limitations, and the risk of spurious alignment driven by temporal structure rather than neural features, we did not consider LE followed by Procrustes to be a reliable approach for cross-subject representation learning in this study. Notably, this approach yielded higher decoding performance than the methods reported here — but we interpret this apparent advantage as a consequence of the information leakage described above rather than as evidence of genuinely better representation learning.

### Prediction of Task Presence Using dFC Features

To evaluate the predictive power of dFC features, we performed supervised classification of task presence versus rest at the single-time-point (TR) level. Each time point in the fMRI scan was treated as a separate sample, with labels derived from the task presence time course (0 = rest, 1 = task). This approach allows us to quantify how well dFC captures moment-to-moment task presence transitions.

A Support Vector Machine (SVM) with RBF kernel was chosen due to its ability to model nonlinear relationships in high-dimensional feature spaces, maintain computational efficiency, and handle class imbalance more robustly than alternatives such as k-nearest neighbors. Initial tests confirmed that replacing SVM with other algorithms (e.g., Random Forests or Gradient Boosting) had minimal effect on the results, but substantially increased computation time. A logistic regression model with L1 penalty was also tested for comparison, with results provided in Supplementary materials.

Train/test splitting was performed at the subject level: 80% of subjects were randomly assigned to the training set and 20% to the held-out test set. All time points from a given subject were kept together in either the training or test set, never split across both, to prevent leakage. Samples were then shuffled within the training set to reduce temporal or subject-order biases. Hyperparameters were tuned using stratified group cross-validation (`StratifiedGroupKFold`), ensuring that class ratios and subject-level dependencies were preserved in each fold. All preprocessing steps, including subject centering, feature scaling, and dimensionality reduction (for state-free methods), were fitted within each training fold and applied to the corresponding validation/test data to prevent information leakage.

The primary evaluation metric was balanced accuracy, which accounts for class imbalance and ensures that chance performance corresponds to 50%. All implementations were conducted using the `scikitlearn` toolbox. This framework allows direct comparison of the predictive efficacy of features obtained using each dFC method and provides a systematic benchmark of their ability to capture task-related dynamics at single-TR resolution.

### Task-Rest Separability

In addition to supervised classification, we evaluated whether dFC features exhibit intrinsic separability of rest and task-present states—that is, whether task and rest time points form distinct structures in feature space independent of any trained model. We refer to this property as task-rest separability.

Following [46], task-rest separability was quantified using the Silhouette Index (SI), which measures how well samples with different labels are separated in the feature space based on pairwise distances. For each dFC method, the SI was computed using the true task labels (task vs. rest) and the corresponding feature vectors. Higher SI values indicate that rest and task-present samples are more distinctly organized in feature space.

Importantly, SI does not involve training a classifier; instead, it evaluates the geometric structure of the feature space with respect to known labels. As such, it provides a complementary, model-independent perspective on whether task-related information is reflected as a globally separable structure, or instead requires supervised methods to be detected.

## Data Availability

All datasets analysed in this study are openly available from the OpenNeuro repository (https://openneuro.org). The 16 datasets, comprising the 28 experimental settings reported here, are accessible under the following accession codes: ds001242 [23], ds002236 [24], ds002647 [25], ds002843 [26], ds002994 [27], ds003465 [28], ds003612 [29], ds003823 [30], ds004044 [31], ds004349 [32], ds004359 [33], ds004556 [34], ds004746 [35], ds004791 [36], ds004848 [37], and ds005038 [38]. Detailed acquisition parameters and the task paradigms associated with each dataset are provided in Table 1. All data were obtained in their raw `NIfTI` form and preprocessed as described in the Methods; the Schaefer 100-region atlas [21] used for region definition was obtained via the `nilearn` toolbox (https://nilearn.github.io). The simulated fMRI data were generated using The Virtual Brain framework (https://thevirtualbrain.org) with the parameters specified in the Methods and can be regenerated us-ing the simulation scripts in the `PydFC` repository (see Code Availability).

## Code Availability

The `PydFC` toolbox [12] used for all dynamic functional connectivity assessment, feature extraction, and analysis in this study is openly available and released under the MIT License. The source code is hosted on GitHub (https://github.com/neurodatascience/dFC) and is installable from PyPI (`pip install pydfc`); the version used here is archived on Zenodo (https://doi.org/10.5281/zenodo.10161176). The toolbox is registered with the SciCrunch Registry (RRID: SCR_024729) and bio.tools (biotoolsID: pydfc). The end-to-end analysis pipeline used in this study — covering conversion of preprocessed NIfTI inputs to parcellated time series, group-level state estimation, subject-level dFC assessment, machine-learning feature extraction and classification, and report generation — is implemented as the modular, BIDS-compatible scripts in the toolbox’s `task_dFC` directory, designed to run sequentially and exchange outputs as saved artefacts for transparent, reproducible execution (including on Slurm cluster environment). The simulation analyses were carried out with the corresponding scripts in the `simul_dFC` directory, which build on The Virtual Brain framework. All method-specific hyperparameter values needed to reproduce the reported results are stated in the Methods.

## Abbreviations

AD: Alzheimer’s Disease
ADHD: Attention-Deficit/Hyperactivity Disorder
ASD: Autism Spectrum Disorders
BOLD: Blood-Oxygen-Level-Dependent
CAP: Co-Activation Patterns
CHMM: Continuous Hidden Markov Model
dFC: dynamic Functional Connectivity
DHMM: Discrete Hidden Markov Model
DOC: Disorders Of Consciousness
FC: Functional Connectivity
fMRI: functional Magnetic Resonance Imaging
FO: Fractional Occupancy
HMM: Hidden Markov Model
LE: Laplacian Eigenmaps
MCI: Mild Cognitive Impairment
PCA: Principal Component Analysis
PD: Parkinson’s Disease
PTSD: Post-Traumatic Stress Disorder
RBF: Radial Basis Function
ROI: Region Of Interest
RSN: Resting State Network
SD: Standard Deviation
SI: Silhouette Index
SVM: Support Vector Machines (SVM)
SW: Sliding Window
SWC: Sliding Window + Clustering
SZ: SchiZophrenia
TF: Time-Frequency
TVB: The Virtual Brain
WL: Window-Less

## Author’s Contributions

J.-B.P. conceived the original idea for the study. M.T. and G.D.M. contributed to refining the study conception. M.T. curated and preprocessed the datasets, developed the `PydFC` toolbox, designed and implemented the analyses and simulations, and wrote the manuscript. J.-B.P. and G.D.M. supervised the study, contributed to the interpretation of the results, and reviewed and edited the manuscript. All authors read and approved the final manuscript.

