## Supplementary materials for "When Can Brain Connectivity Track the Working Mind? A Large-Scale Benchmark of Dynamic Functional Connectivity Across Cognitive Paradigms"

In this document, we present supplementary data that offers additional insights and results.

### Task Paradigm Descriptions

We organized task paradigms using the Research Domain Criteria (RDoC) framework: each subsection corresponds to an RDoC *domain* and each subsubsection to a core *construct* within that domain (e.g., Cognitive Systems → Working Memory). This structure clarifies which cognitive, affective, or sensorimotor processes each paradigm is intended to manipulate or measure, and facilitates comparisons across datasets. For transparency and reproducibility, we retained the study's canonical BIDS-style task codes in parentheses (e.g., *task-Stroop*). Some paradigms (e.g., Fribble Value Construction) may support multiple constructs; we included them in the closest domain.

#### A. Arousal and Regulatory Systems.

##### *A.1. Arousal.*

**Emotion Regulation / HRV Intervention (*task-emotionRegulation*).** Participants practiced slow-paced breathing or individualized biofeedback strategies to modulate heart-rate variability. The task probes how physiological self-regulation alters emotion-related network connectivity.

#### B. Cognitive Systems.

##### *B.1. Attention.*

**Neurofeedback (*task-feedback*).** Participants viewed moving bars linked to real-time brain activity. Depending on condition, they attempted to control the bars or monitored them passively, probing self-regulation and attentional control.

##### *B.2. Cognitive Control.*

**AX-CPT (*task-Axcpt*).** Participants viewed sequential letters and responded only when X followed A. Different trial types vary context and interference (e.g., BX, BY), creating high vs. low cognitive control demands.

**Cued Task-Switching (*task-Cuedts*).** A cue instructed participants to attend either the number or the letter in a letter-digit pair. Incongruent stimuli increase task-switching demands; participants performed odd/even or vowel/consonant judgments depending on the cue.

**Sensorimotor Choice Reaction Tasks Localizer (*task-Localizer*).** Participants completed short blocks of visual and auditory stimuli used to define modality-specific and multimodal regions of interest.

**Sensorimotor Choice Reaction Tasks (*task-ST*).** Participants responded to visual or auditory stimuli with manual or vocal responses. Participants had to respond to the side of the stimuli by either pressing a button with their right or left hand (index finger) and/or by saying the German word for "right" and "left".

**Stroop (*task-Stroop*).** Participants named the color of visually presented words spoken aloud in the scanner. Incongruent stimuli evoke strong interference, engaging conflict resolution and cognitive control.

##### *B.3. Language.*

**Auditory Semantic Judgment, Visual Rhyming, Visual Semantic, Visual Spelling (*task-AudSem*, *task-VisRhyme*, *task-VisSem*, *task-VisSpell*).** Participants made lexical judgments in auditory or visual modalities, determining whether word pairs rhymed, shared spelling patterns, or were semantically related. These tasks engage phonological, orthographic, and semantic language processes.

##### *B.4. Perception.*

**Exposure to Face Morphs (*task-expo*).** Participants passively viewed blended face images across blocks. This task probes visual perception and face-processing networks.

**Task-Specific Localiser (*task-localiser*).** In category localiser scans, participants viewed images of faces, scenes, and phase scrambled versions of the face images. These can be used to define face and scene selective regions of interest.

**Item Matching (*task-matching*).** Participants judged whether pairs of numbers, shapes, or faces matched. The task probes symbolic and non-symbolic quantity processing, visual comparison, and feature matching.

**Parahippocampal Place Area (PPA) Localizer (*task-ppalocalizer*).** Participants completed a spatial detection task involving place-object image pairs following conditioned tones. The participants' task was to quickly indicate whether the high-salience image was on the right or left via a button press. The task is designed to localize parahippocampal place area responses to scene stimuli.

###### B.5. Working Memory.

**Arithmetic (*task-arithmetic*).** Participants evaluated whether single-digit addition problem solutions were correct or incorrect. This task probes basic arithmetic processing, numerical reasoning, and parietal cortex engagement.

**Movie Recall (*task-recall*).** After watching a shortened version of *The Sixth Sense* in a separate scan, participants recalled movie scenes during fMRI. The task engages episodic memory retrieval and high-level narrative reconstruction.

**Sternberg Working Memory (*task-Stern*).** Participants memorized short word lists and judged whether a later probe was present. "Recent negative" probes induce interference, increasing working memory and cognitive control demands.

**Visuospatial Working Memory (*task-vswm*).** Participants tracked the movement of a dot across a grid and judged whether a later probe matched previous positions. Load manipulations test visuospatial working memory capacity and maintenance.

##### C. Negative Valence System.

###### C.1. Acute Threat ("Fear").

**Fear Conditioning (*task-fearlearning*).** Participants learned associations between a conditioned tone and mild shock versus a safe tone. The task engages learning, arousal, and anticipatory threat processing.

**Pain Generation (*task-paingen*).** Participants received thermal or mechanical stimuli across placebo and control runs. This task investigates pain processing, modulation, and expectation effects on brain networks.

##### D. Positive Valence System.

###### D.1. Reward Responsiveness.

**Intertemporal Choice (*task-itc*).** Participants chose between a smaller immediate reward and a larger delayed reward. The task measures delay discounting and subjective valuation of future rewards.

**Risky Choice (*task-risk*).** Participants chose between a guaranteed $20 reward and a larger uncertain reward. The task probes risk preferences and decision-making under uncertainty.

###### D.2. Reward Valuation.

**Fribble Value Construction (*task-fribBids*).** Participants evaluated or chose between "fribbles," multi-attribute artificial objects used to study value construction. The task probes multi-attribute decision making and valuation processes in the ventromedial prefrontal cortex (vmPFC), a region consistently implicated in encoding subjective value.

##### E. Sensorimotor Systems.

###### E.1. Motor Actions.

**Execution (*task-execution*).** Participants squeezed a rubber ball with different hands following visual cues. The task examines motor execution and lateralized motor network engagement.

**IHG: Isometric Handgrip (*task-IHG*).** Participants alternated between rest and strong squeezes of a handgrip ball following visual cues. The task engages sympathetic arousal, motor control, and autonomic regulation.

**Motor Imagery (*task-imagery*).** Participants imagined squeezing a rubber ball with different hands. The task allows comparison of motor imagery versus execution and the influence of handedness on motor representations.

**Motor Execution Mapping (*task-motor*).** Participants performed small targeted movements of body parts (e.g., fingers, toes, jaw, tongue). This broad somatomotor task localizes motor and premotor cortical representations.

#### General Linear Model (GLM) Analysis for a Representative Experiment

This section presents a representative General Linear Model (GLM) analysis to demonstrate the quality and validity of the fMRI data and preprocessing pipeline. The GLM was applied to a single subject's run from `EXP.19` (*task-expo*; Exposure to Face Morphs paradigm) using the `nilearn` implementation of first-level modeling. The design matrix was constructed using the canonical hemodynamic response function (HRF) and task timing extracted from the corresponding `events.tsv` file. The resulting contrast (*active* - *rest*) reveals significant activation primarily in the occipital cortex, reflecting expected visual engagement during face presentation. This analysis provides additional validation of preprocessing quality and confirms the reliability of task-related BOLD fMRI responses in the analysis.

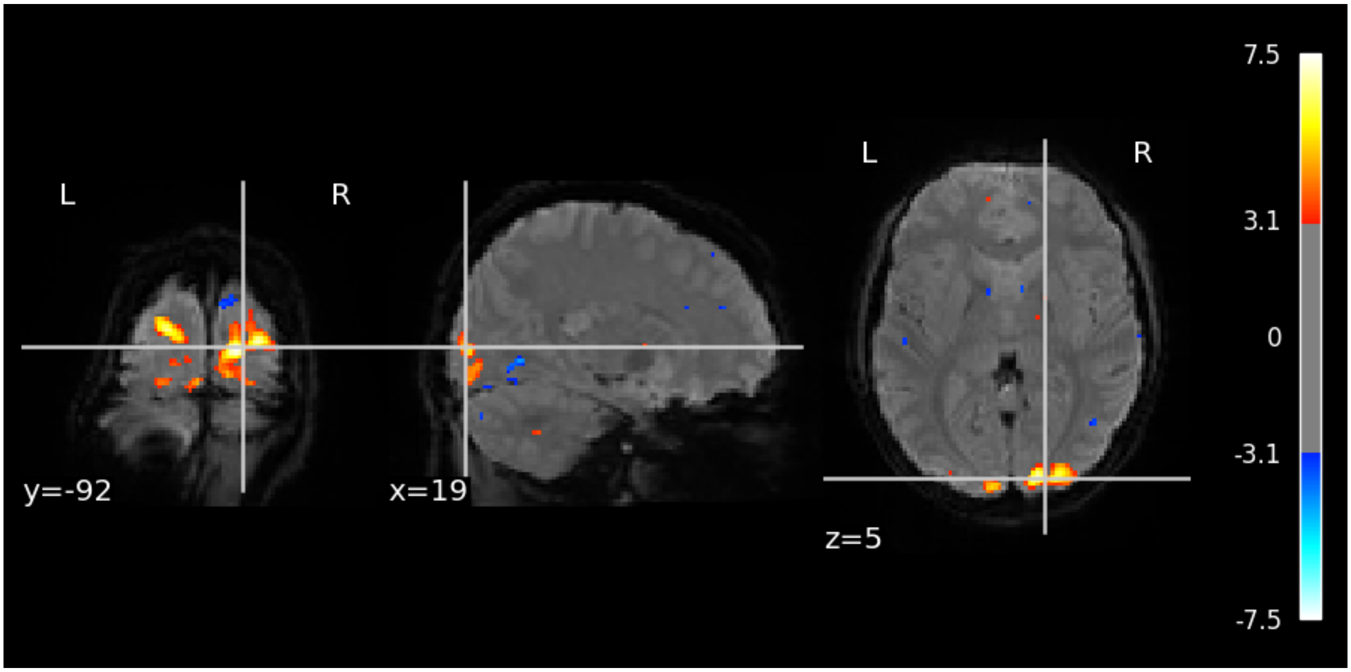


**Fig. 8. Example GLM activation map for `EXP.19` (*task-expo*; Exposure to Face Morphs paradigm).** First-level GLM analysis of one subject's run demonstrates significant task-related activation in visual regions during face presentation, shown on a background mean functional image. The contrast (*active* - *rest*) was estimated using the canonical HRF and task timings derived from `events.tsv`. Thresholded statistical map ($Z > 3.1$) highlights robust activation in the occipital cortex, confirming expected engagement of the visual network and validating the overall data quality.

#### Extended Analysis of Cohen's D Effect Sizes Across Experiments

This section provides additional analyses of the task-rest signal contrast — the Cohen's d of the BOLD difference between task and rest — across brain regions and experiments. These supplementary figures characterize how this contrast varies across regions and experiments.

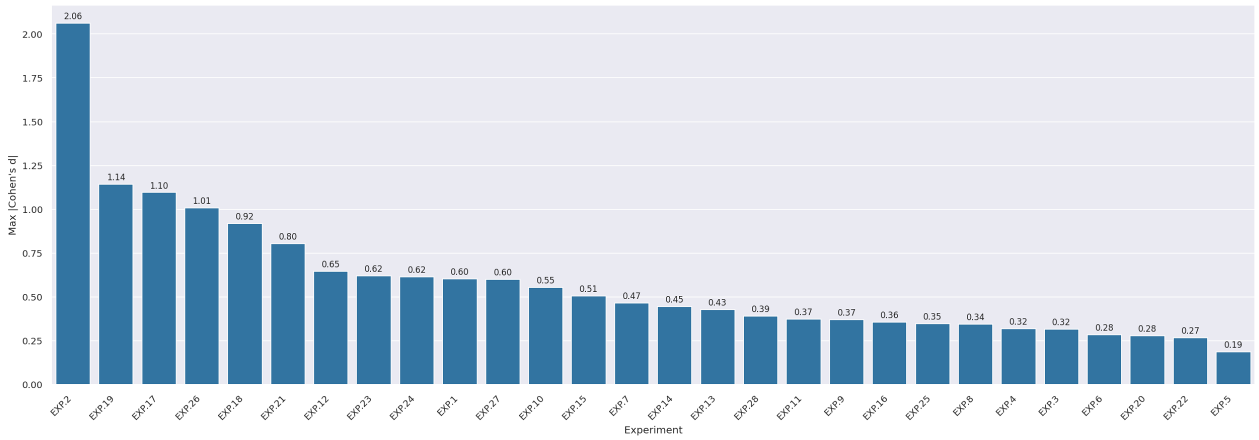


**Fig. 9. Maximum absolute Cohen's D values across experiments.** Each bar represents the largest absolute Cohen's D effect size observed among brain regions for a given experiment, sorted by decreasing maximum value. Experiments such as `EXP.2`, `EXP.19`, `EXP.17`, and `EXP.26` exhibit the strongest regional contrasts between task and rest.

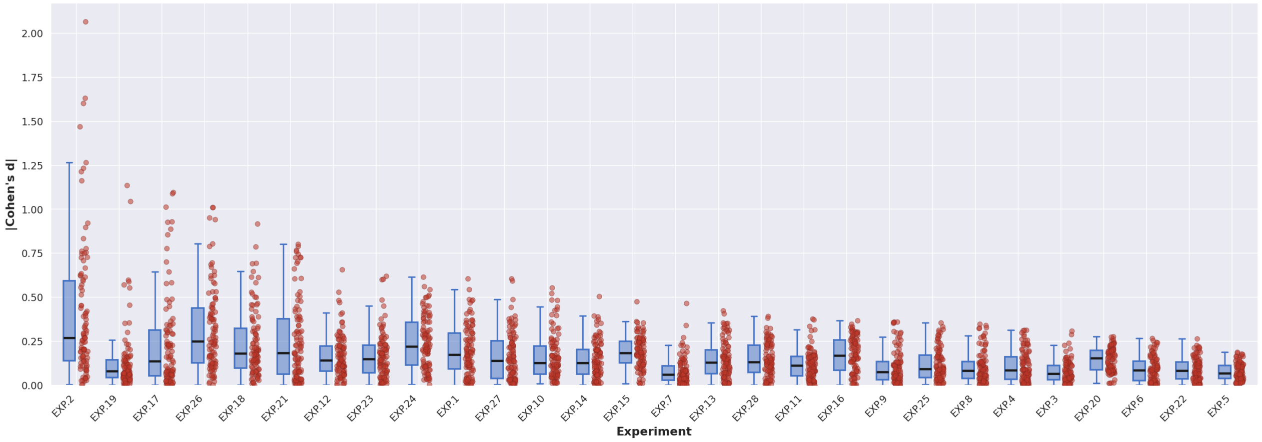


**Fig. 10. Distribution of Cohen's D effect sizes across regions in experiments from experimental datasets.** Boxplots show the distribution of absolute Cohen's D values across all brain regions for each experiment, sorted by maximum effect size (as in Figure9). Experiments such as `EXP.2` and `EXP.26` display the largest median effect sizes, indicating widespread and robust task-related activation.

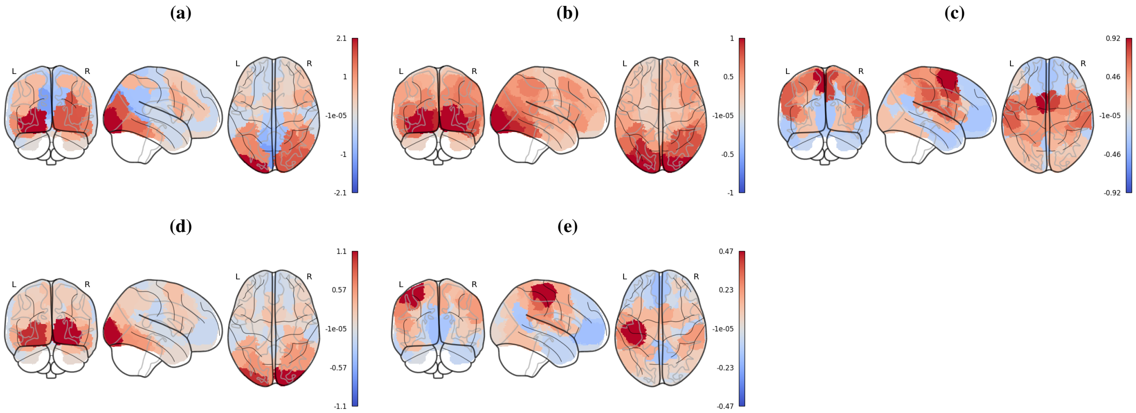


**Fig. 11. Spatial distribution of Cohen's D effect sizes across representative experiments.** Glass brain maps show region-wise Cohen's D values for (a) `EXP.2`, (b) `EXP.26`, (c) `EXP.18`, (d) `EXP.19`, and (e) `EXP.7`. Stronger activations are predominantly observed in expected task-related cortical regions, e.g., visual cortex for `EXP.26` (`task-localiser`) or motor cortex for `EXP.18` (`task-motor`).

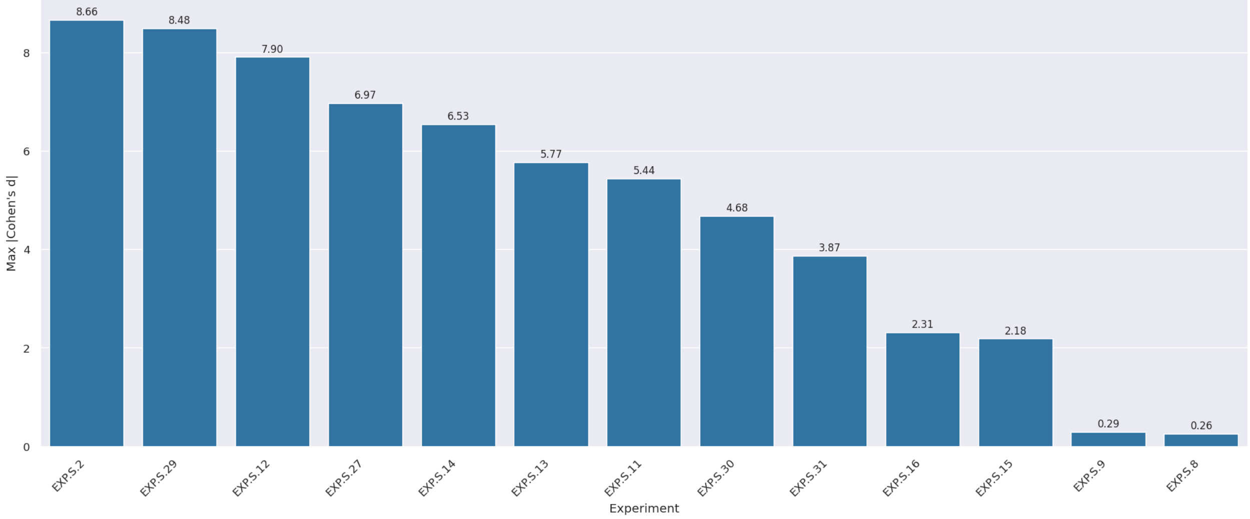


**Fig. 12. Maximum absolute Cohen's D values across simulated experiments.** Each bar represents the largest absolute Cohen's D effect size observed among brain regions for a given experiment, sorted by decreasing maximum value. Experiments such as `EXP.S.2` and `EXP.S.29` exhibit the strongest regional contrasts between task and rest.

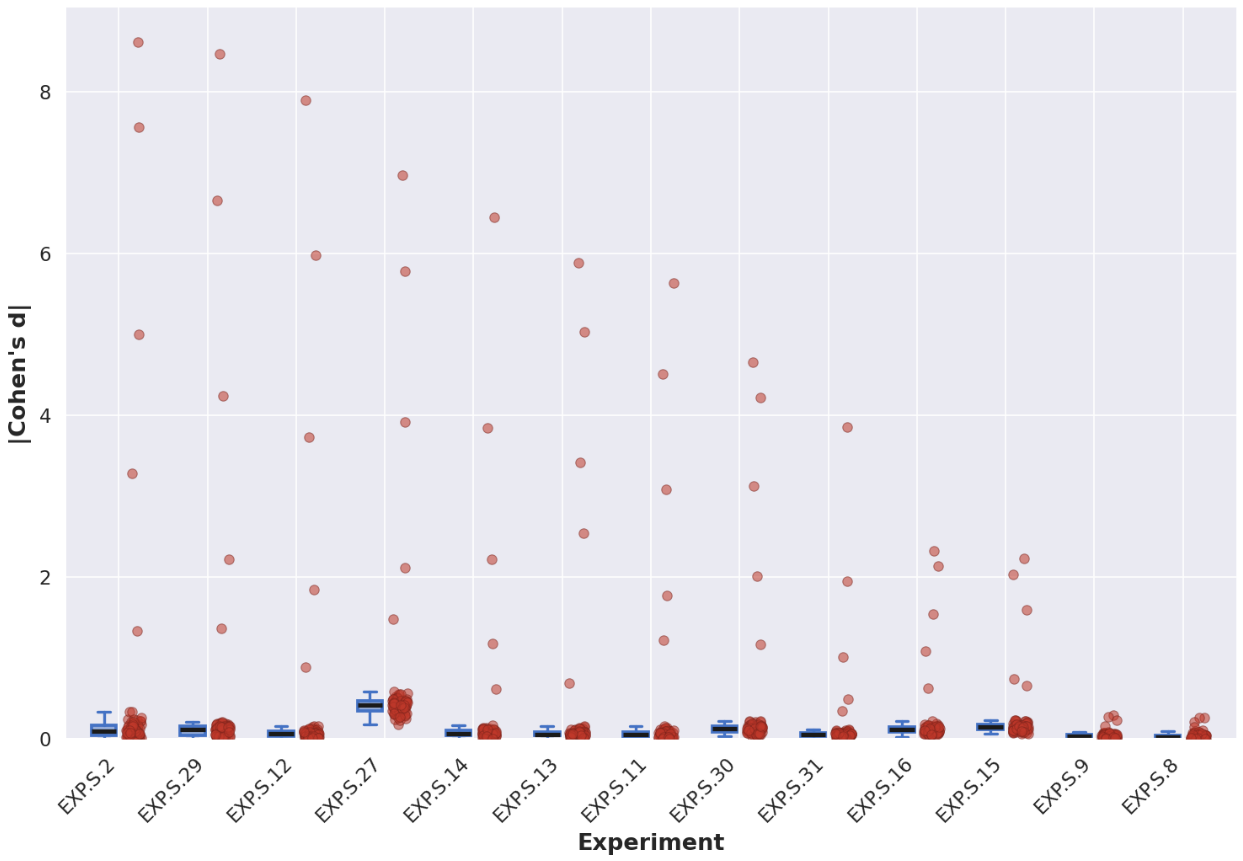


**Fig. 13. Distribution of Cohen's D effect sizes across regions in simulated experiments.** Boxplots show the distribution of absolute Cohen's D values across all brain regions for each experiment, sorted by maximum effect size (as in Figure12). `EXP.S.27` displays the largest median effect sizes, indicating widespread and robust task-related activation.

#### Task Timing Design and Binarization Across Experiments

To illustrate how task design influences the extraction of task presence labels, Figures 14 and 15 visualize the task timing and Gaussian Mixture Model (GMM)-based binarization process for representative experiments. Each panel displays the original task or stimulus timing (golden yellow), its convolution with a canonical hemodynamic response function (black), and the resulting task presence labels (task-present in blue, rest in grey). Vertical red dashed lines correspond to MRI repetition times (TRs). The HRF-convolved signals are plotted at a higher temporal resolution for clarity. Together, these examples highlight how experimental design and temporal structure shape the clarity and balance of task presence labels used for prediction.

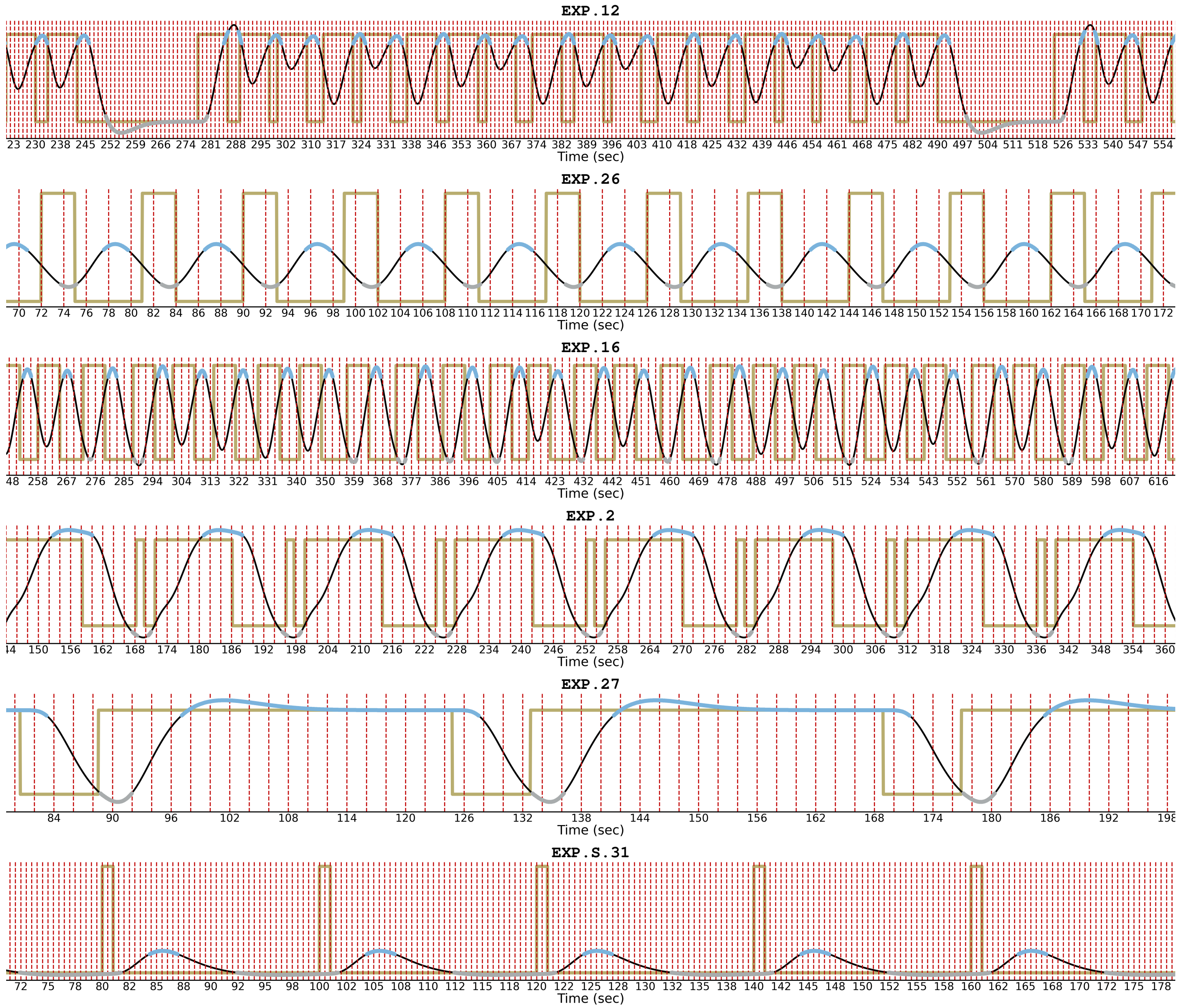


**Fig. 14. Examples of experiments with effective task designs and successful GMM-based binarization.** Panels show experiments with clear task-rest alternations and well-defined HRF-convolved responses: `EXP.12`, `EXP.26`, `EXP.16`, `EXP.2`, `EXP.27`, and `EXP.S.31`. Gold lines show original task/stimulus timings, black lines the HRF-convolved signals, and blue and grey shading denote task-present and rest segments identified by GMM binarization. Vertical dashed red lines mark MRI sampling points (TRs). Experiments such as `EXP.26` exhibit clear periodicity and balanced task-rest durations, leading to clean label separation. Notably, `EXP.12`-despite its more complex, less periodic timing compared to experiments such as `EXP.26`-exemplifies a well-designed paradigm that supports clear and balanced binarization and ultimately high classification performance. Together, these examples demonstrate how thoughtful task design facilitates reliable task-rest separability and robust downstream analysis.

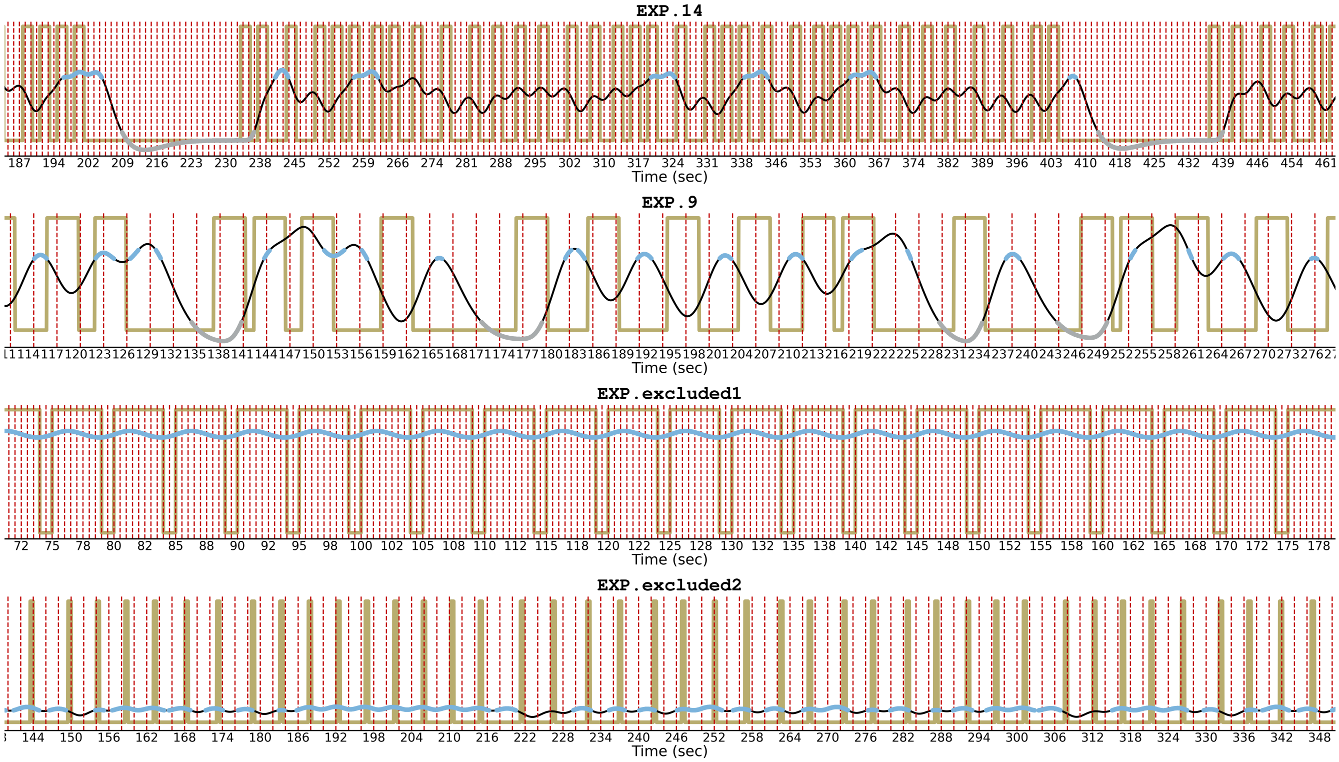


**Fig. 15. Examples of experiments with suboptimal task designs or limited predictive power for task presence labels.** Panels show experiments with irregular timing or extreme class imbalance: `EXP.14`, `EXP.9`, `EXP.excluded1`, and `EXP.excluded2`. Gold lines indicate original task/stimulus timings, black lines represent HRF-convolved signals, and shaded regions correspond to task-present (blue) and rest (grey) intervals identified by GMM binarization. `EXP.excluded1` and `EXP.excluded2` are examples of excluded experiments. In experiments such as `EXP.excluded1`, the rapid alternation between task and rest causes the HRF-convolved signal to remain almost entirely in the task-present range, leading to severe class imbalance and unreliable label boundaries. In experiments such `EXP.excluded2`, the extremely short task duration causes a flat HRF-convolved signal. These are only two examples of suboptimal task design. `EXP.9` exhibited weak decoding performance. `EXP.14` despite its suboptimal design, exhibited strong downstream decoding performance. These examples illustrate how suboptimal temporal design can obscure task presence transitions and limit dFC-based task-rest decoding performance.

#### Task Timing Statistics Across Experiments

This section summarizes the temporal properties of task and rest periods across all experiments, providing quantitative context for the observed classification and separability results. For each experiment, we examined the distribution of individual task and rest block durations, the proportion of total scan time occupied by task-present periods, the relative frequency of rest-task transitions, as well as the distribution of Optimality Index (OI) — a measure that compares the frequency content of the task design to that of an ideal design, with higher values indicating better-aligned, more optimal task timing [16]. Some of the candidate factors used in the top-vs-bottom performance analysis in the main text (Figures 5 and 6) were derived from these properties; here we report their full distributions across experiments to provide descriptive context. Together, these statistics capture important aspects of experiment design that might affect dFC analysis.

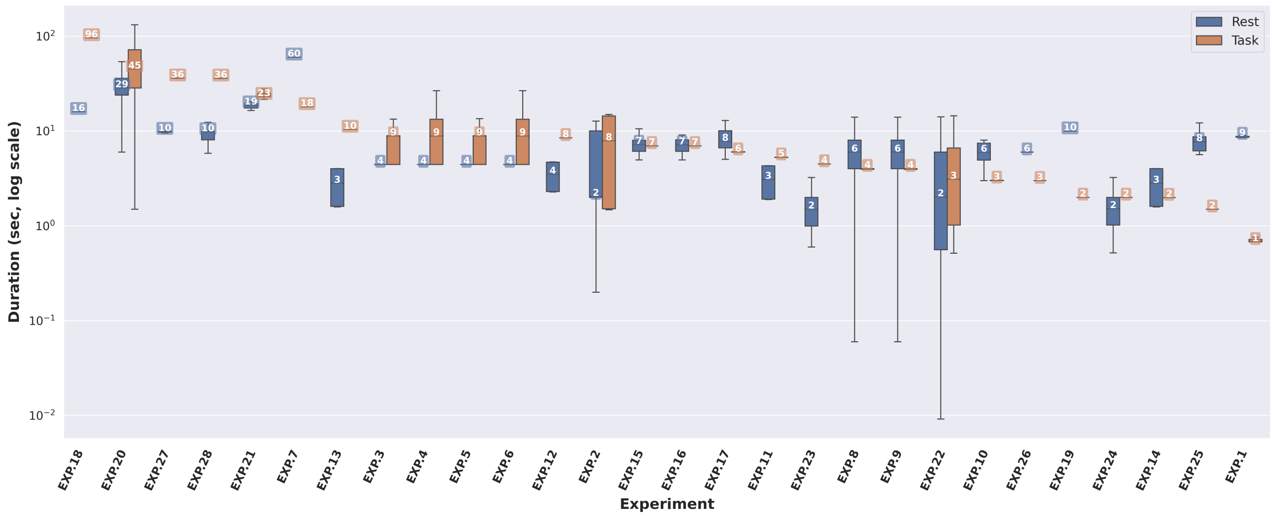


**Fig. 16. Distribution of task and rest block durations across experiments from experimental datasets.** Each pair of boxplots represents the distribution of individual task (orange) and rest (blue) block durations, aggregated across all subjects and scans for each experiment. Durations are shown in seconds on a logarithmic scale for better visualization. Median values are annotated above the median lines. These durations correspond to individual task or rest blocks, not total task or rest time within a scan, and are derived directly from the original task timing (`events.tsv`) files.

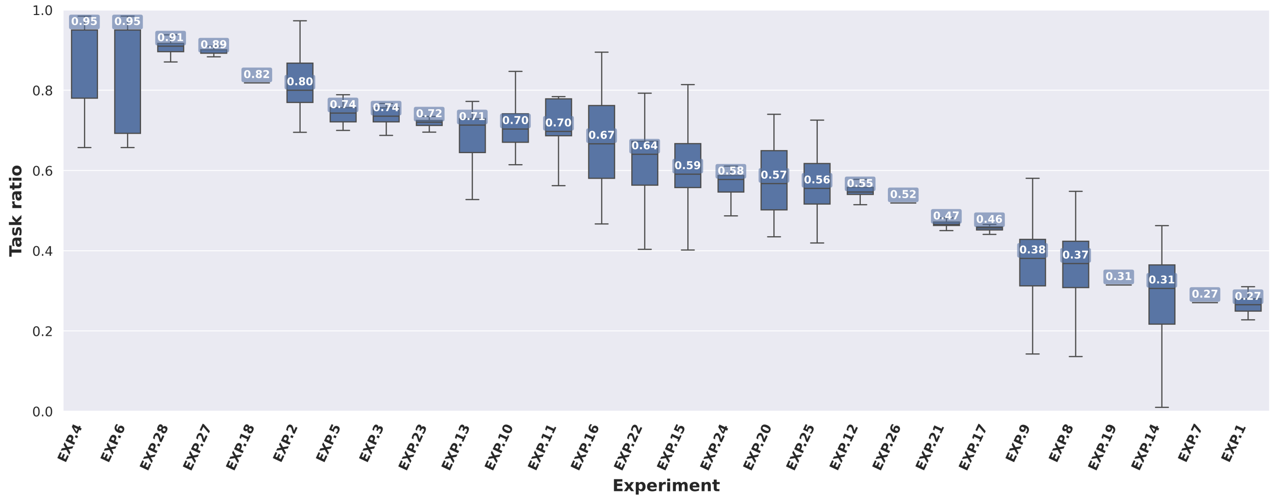


**Fig. 17. Proportion of scan time spent in task-present condition across experiments from experimental datasets.** Boxplots represent the distribution of the ratio of task-present duration to total scan duration across all subjects and scans. Ratios are computed from the task presence time courses after Gaussian Mixture Model (GMM) binarization and removal of transition samples. Median ratio values are indicated above the median lines. Experiments with ratios near 0.5 have well-balanced task and rest periods, whereas extreme ratios indicate imbalance that can affect classification performance.

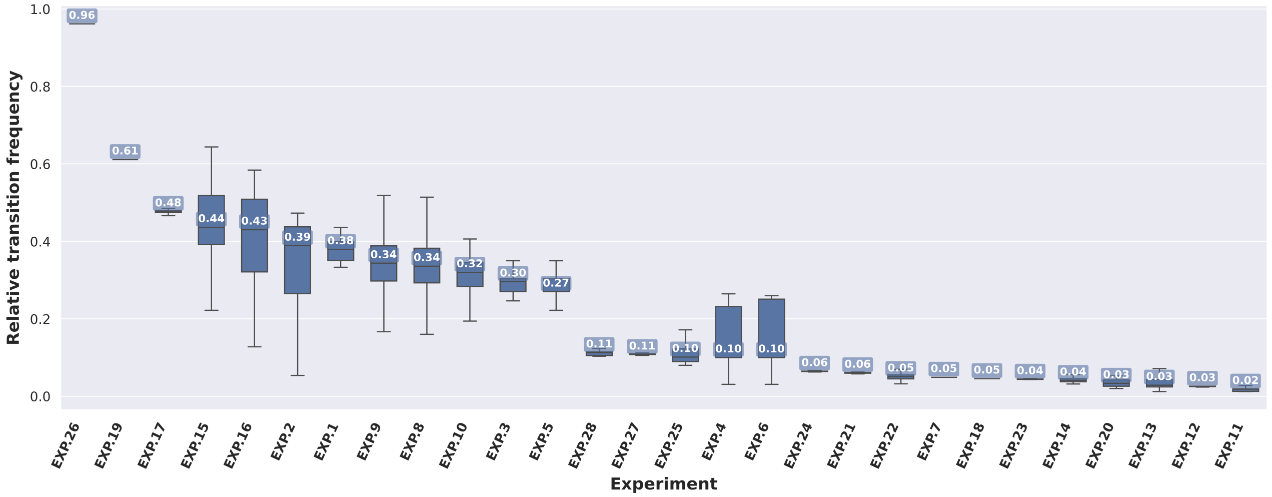


**Fig. 18. Relative transition frequency across experiments from experimental datasets.** Boxplots show the distribution of relative transition frequency (number of task-to-rest or rest-to-task transitions divided by total time points) across all scans and subjects. Computations are based on GMM-binarized task presence time courses after discarding transition samples. Median values are annotated above the median lines. This metric reflects how frequently changes in task presence occur within each experiment, influencing both dFC estimation and classifier performance.

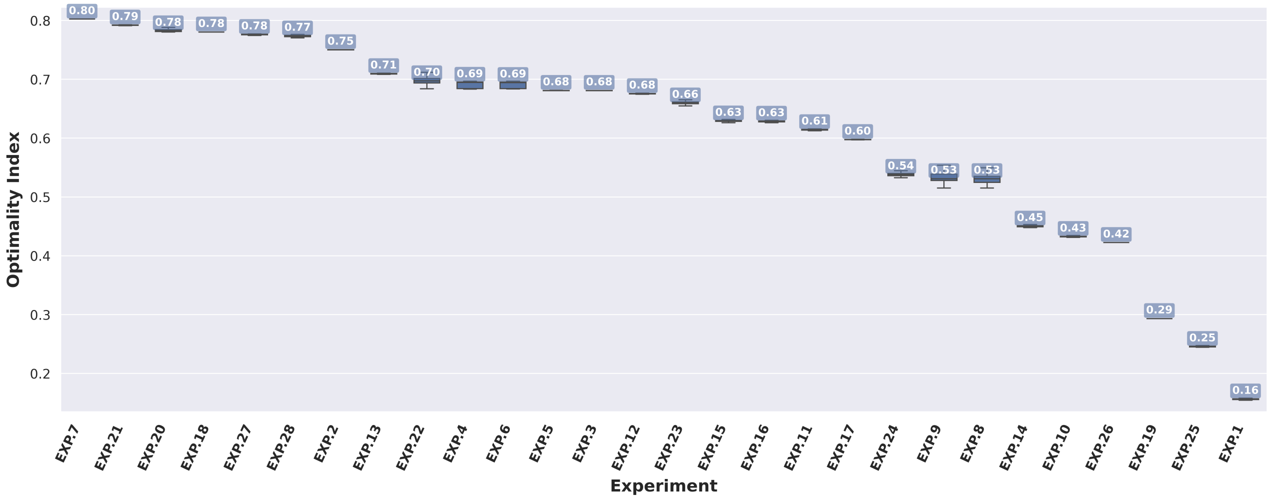


**Fig. 19. Optimality Index (OI) across experiments from experimental datasets.** Boxplots show the distribution of OI across all scans and subjects. Computations are based on original task timing (from `events.tsv`). Median values are annotated on the median lines. The OI quantifies the optimality of task timing by comparing the frequency content of the task design to that of an ideal design, with higher values indicating better alignment and therefore more optimal task structure.

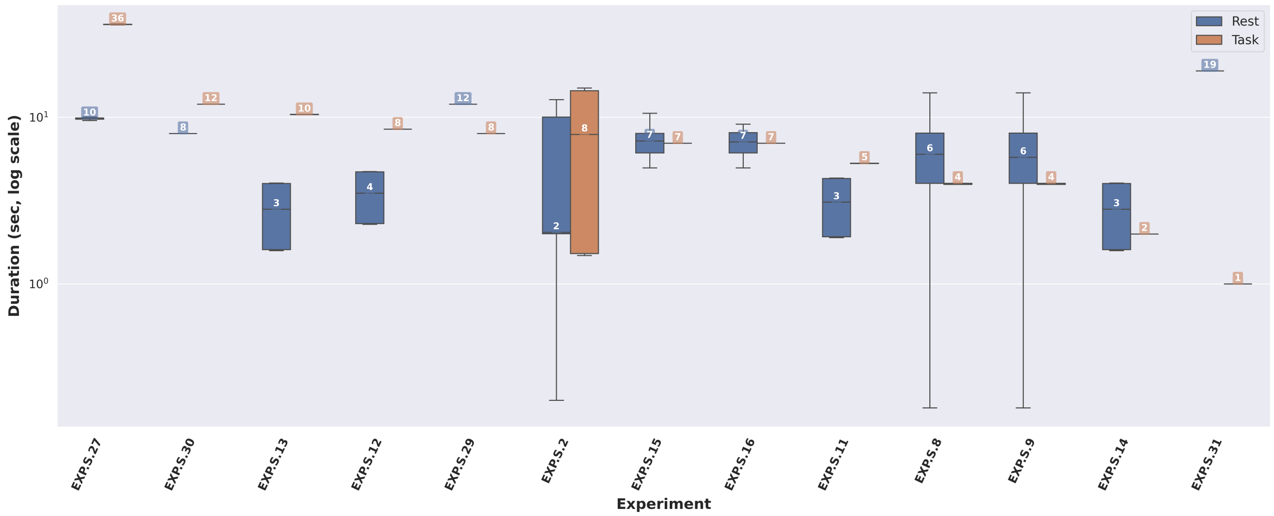


**Fig. 20. Distribution of task and rest block durations across simulated experiments.** Similar to Figure16, this figure shows task (orange) and rest (blue) block duration distributions for simulated experiments. Durations are in seconds and shown on a logarithmic scale.

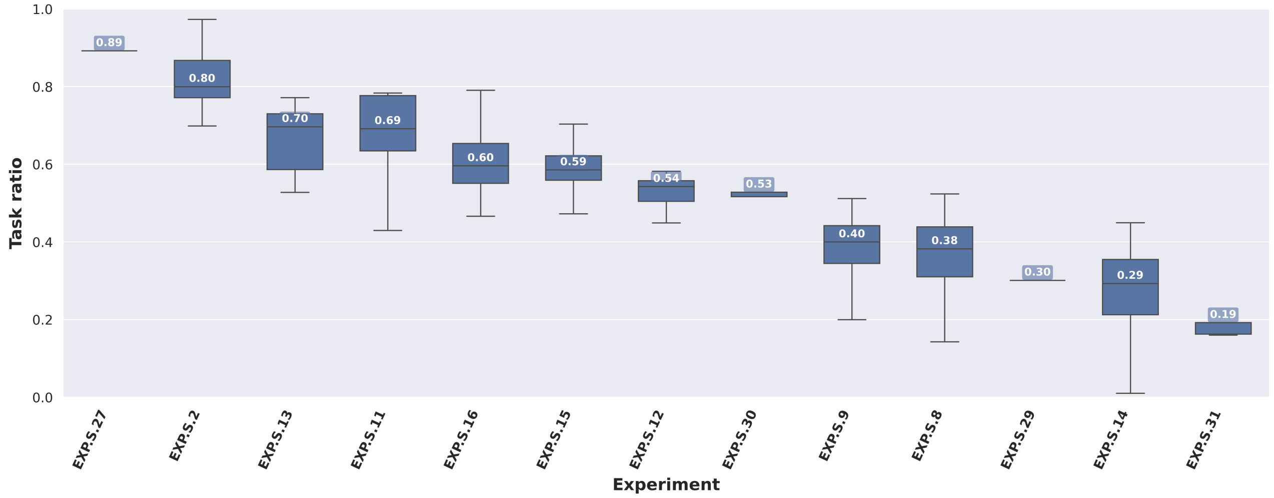


**Fig. 21. Proportion of scan time spent in task-present condition across simulated experiments.** The ratio of total task-present duration to total scan length is shown for simulated experiments, based on task presence time courses after GMM binarization and transition removal. Median values are annotated on the median lines.

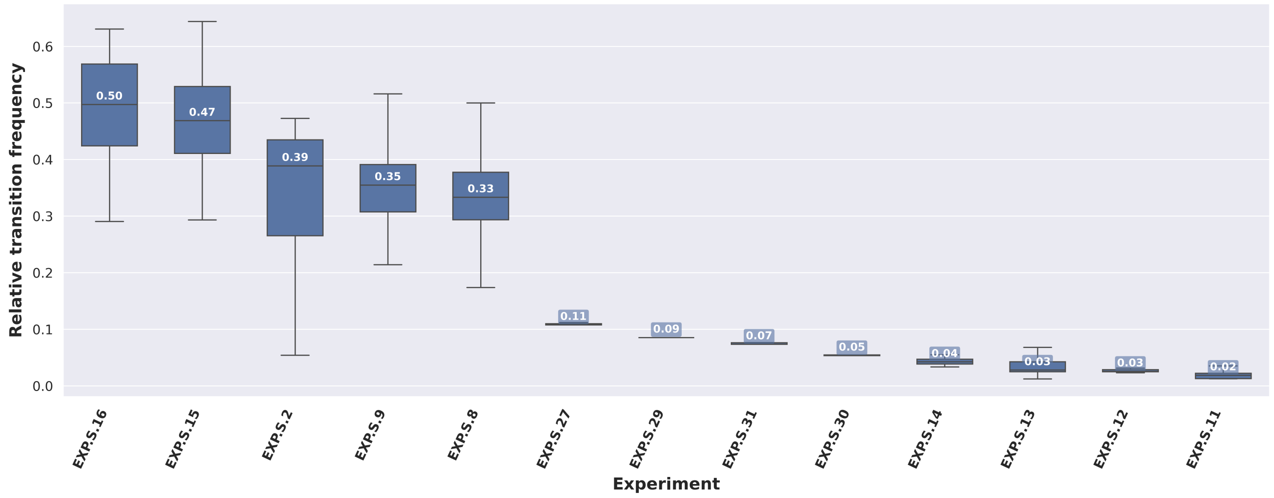


**Fig. 22. Relative transition frequency across simulated experiments.** Boxplots depict the frequency of task-rest transitions normalized by scan length, for simulated experiments. Transition frequencies are derived from post-GMM task presence time courses.

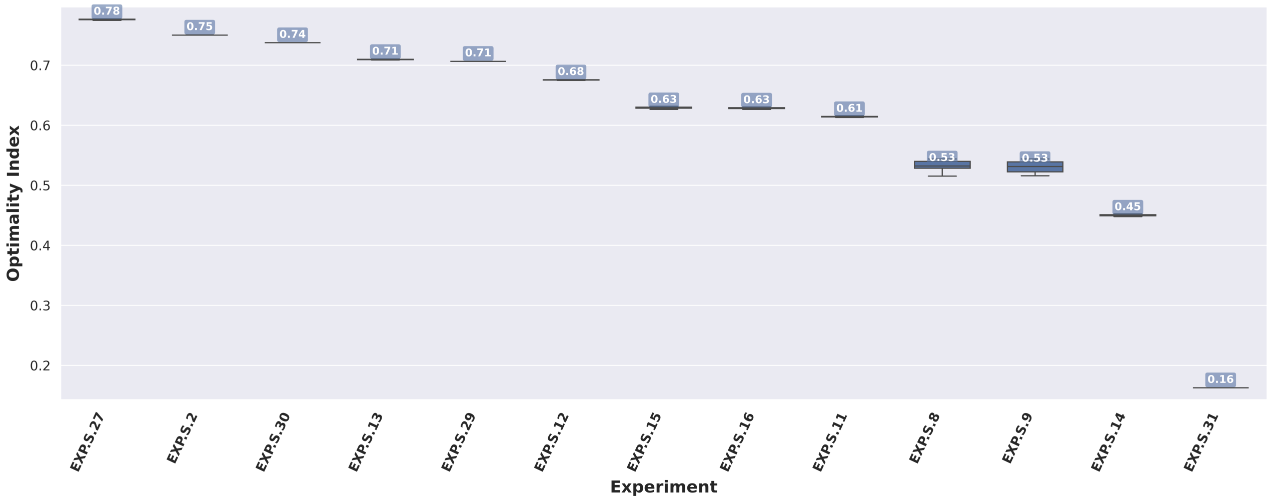


**Fig. 23. Optimality Index (OI) across simulated experiments.** Boxplots show the distribution of OI across all scans and subjects. Computations are based on original task timing (from `events.tsv`). Median values are annotated on the median lines. The OI quantifies the optimality of task timing by comparing the frequency content of the task design to that of an ideal design, with higher values indicating better alignment and therefore more optimal task structure.

#### Distribution of Temporal Signal-to-Noise Ratio Across Experiments

We quantified the temporal signal-to-noise ratio (tSNR) for each fMRI run to characterize differences in signal quality across datasets. For each 4D BOLD image, voxel-wise tSNR was computed as the temporal mean divided by the temporal standard deviation. A run-specific EPI brain mask was extracted, and tSNR values were summarized within this mask using the median, yielding a single scan-level tSNR value per run.

Figure 24 summarizes the distribution of scan-level median tSNR values across all runs and subjects for each experiment. Each point corresponds to one fMRI run, and boxplots therefore reflect between-scan variability within each experiment.

Substantial variability in tSNR is observed both across and within experiments. Median tSNR values vary widely across experiments, ranging from approximately 26 to 87, indicating differences in overall signal quality between datasets. In addition, several experiments—particularly those with higher median tSNR—exhibit large within-experiment variability across runs, with values spanning a broad range.

These results highlight that signal quality is not uniform across experiments and may contribute to variability in decoding performance. However, as shown in the main analyses, differences in tSNR alone do not fully explain performance variability (see Figure5).

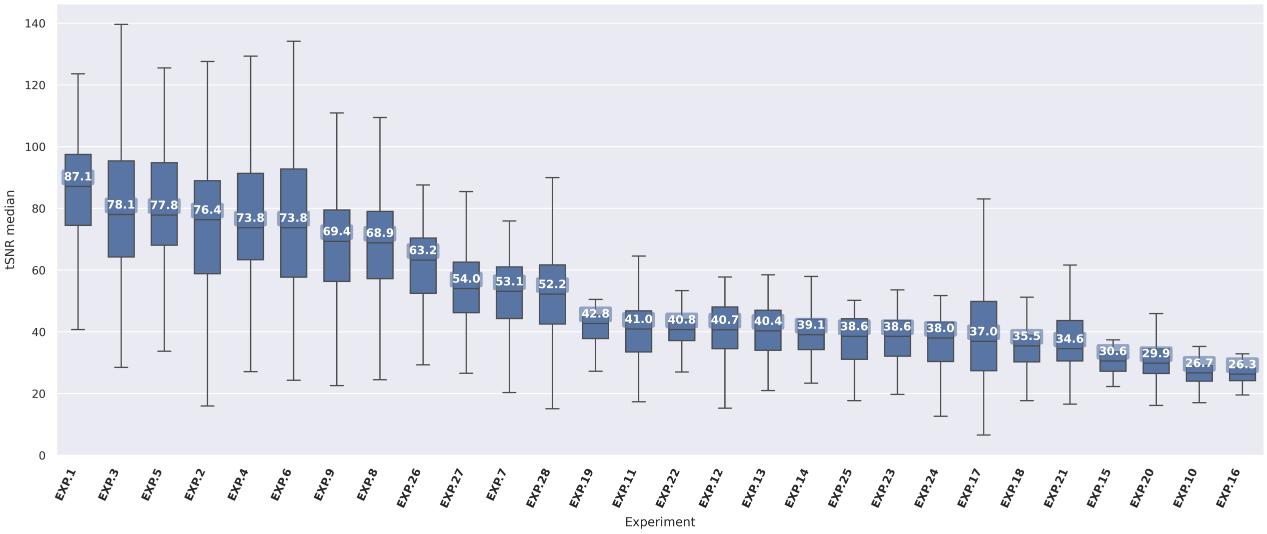


**Fig. 24. Distribution of scan-level median temporal signal-to-noise ratio (tSNR) across experiments.** For each fMRI run, voxel-wise tSNR was computed as the temporal mean divided by the temporal standard deviation, and summarized within a run-specific brain mask using the median, yielding a single tSNR value per scan. Each point represents one run from one subject. Boxplots therefore reflect the distribution of scan-level median tSNR values across all runs and subjects within each experiment. Experiments are ordered by median tSNR. Substantial variability is observed both across experiments (median range: ∼26–87) and within experiments, with some showing wide dispersion across runs, indicating heterogeneity in signal stability.

#### Example Dynamic Functional Connectivity (dFC) Patterns Across Methods

To illustrate how different dFC methodologies capture transient changes in whole-brain connectivity, Figure25 and Figure26 visualize sample dFC patterns obtained from experimental and simulated experiments. For each experiment, a short 10-time-point segment is extracted at the midpoint of a representative scan. Each row corresponds to one of the seven dFC assessment methods, and each column corresponds to a single time point. All dFC matrices are rank-normalized per scan and method for visual comparability across methods.

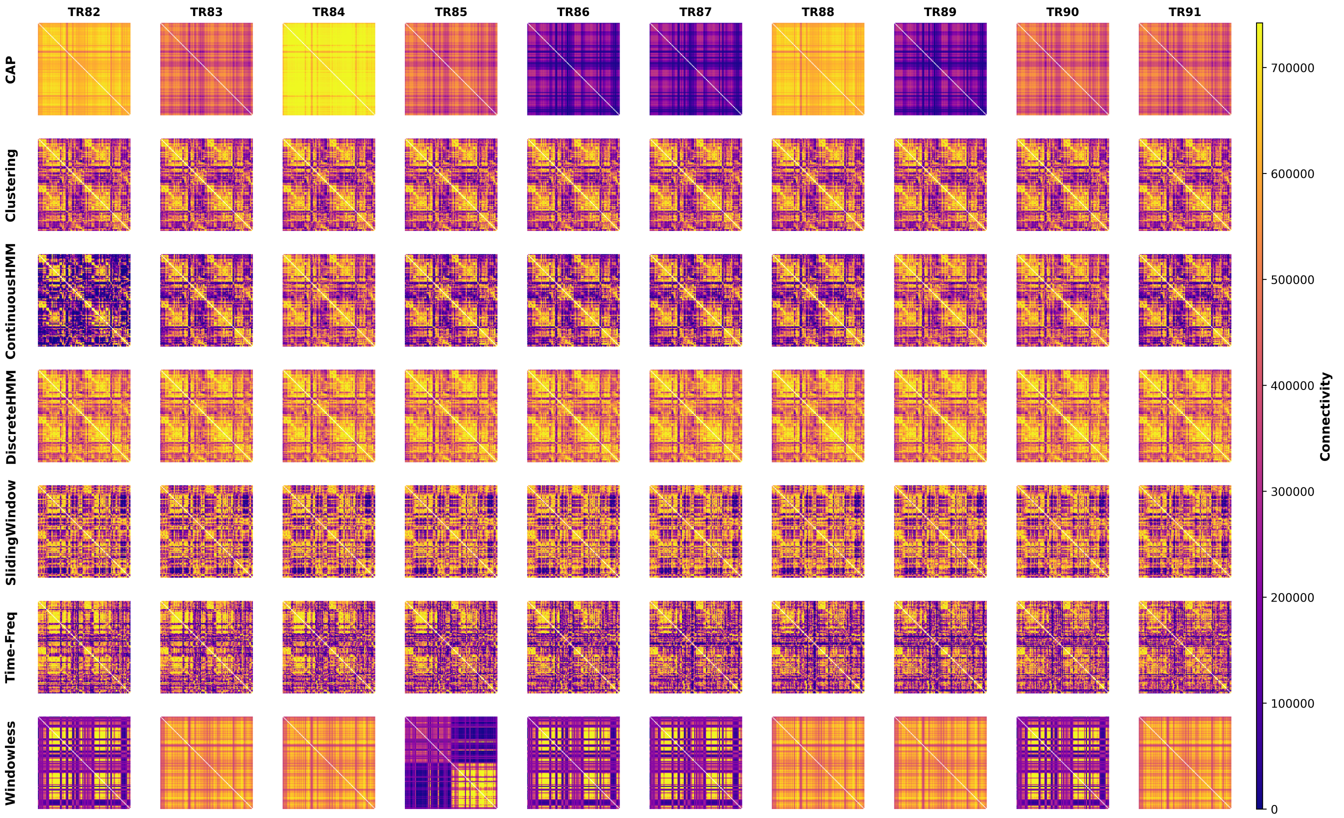


**Fig. 25. Representative dFC patterns from `EXP.1` (experimental data).** Each row shows dFC matrices derived from one of the seven examined dFC assessment methods, and each column represents a consecutive time point within a 10-TR segment from a single subject's scan. The matrices depict pairwise regional connectivity patterns computed at each time point, rank-normalized within method for visualization. This visualization highlights the diversity of temporal dynamics expressed by different methodological families when applied to the same empirical dataset.

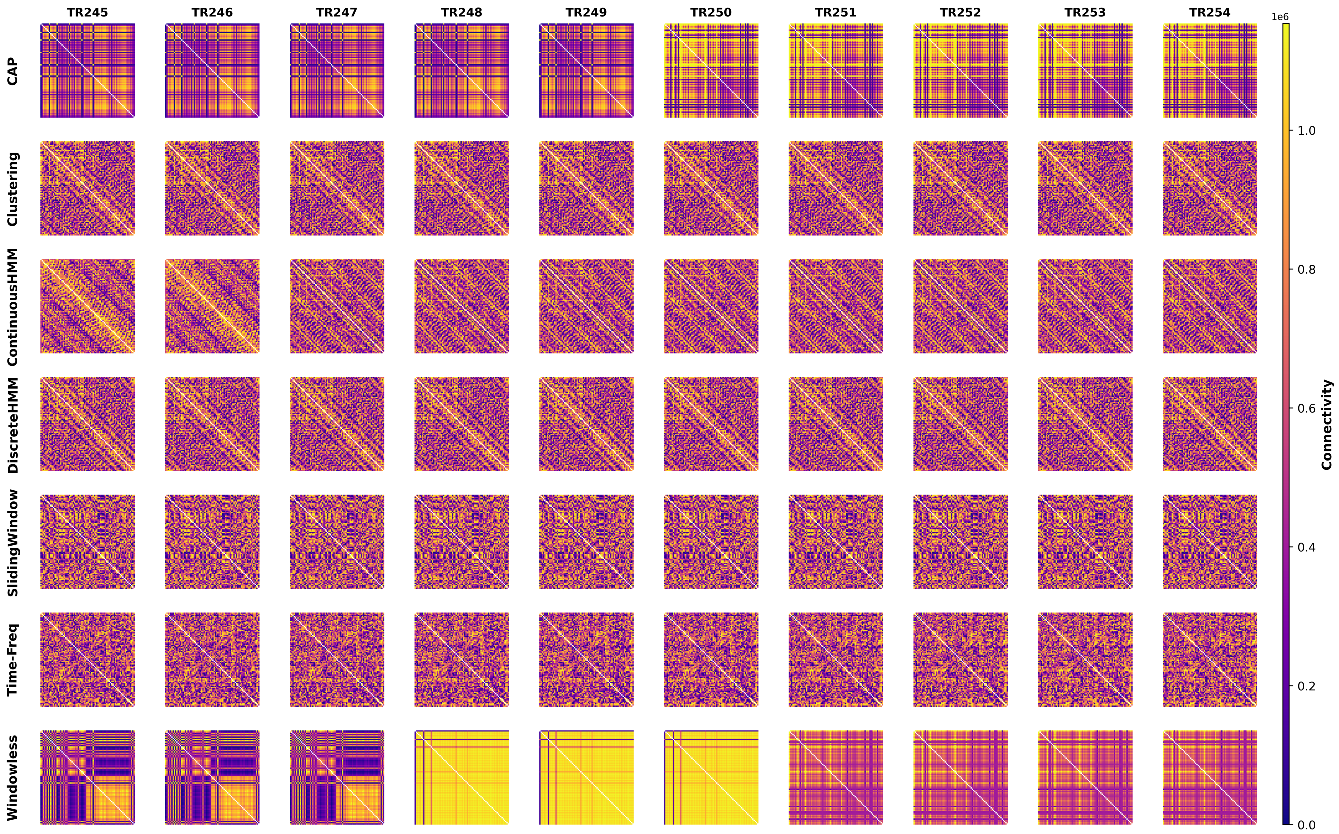


**Fig. 26. Representative dFC patterns from `EXP.S.29` (simulated data).** Same visualization format as Figure25, showing 10 consecutive time points from one simulated subject. Patterns were generated using The Virtual Brain framework under idealized conditions with known task timing.

#### Cognitive Domain and Decoding Performance (Exploratory Analysis)

As an exploratory analysis, we examined whether decoding performance varies across cognitive domains using the Research Domain Criteria (RDoC) framework (Figure 27).

Broadly, experiments categorized under Cognitive Systems and Sensorimotor Systems exhibited higher average performance and a greater proportion of high-performing experiments compared to other domains. In contrast, experiments within the Negative Valence Systems domain tended to show lower decoding performance.

However, these patterns should be interpreted with caution. The number of experiments is not evenly distributed across domains, and substantial variability remains within each domain. In addition, domain-level differences are confounded by variations in task design, dataset characteristics, and signal properties, which are not controlled for in this analysis.

Overall, these results suggest that cognitive domain may influence decoding performance, but do not support strong or generalizable conclusions. More targeted and balanced experimental designs will be required to isolate the contribution of cognitive domain from other factors.

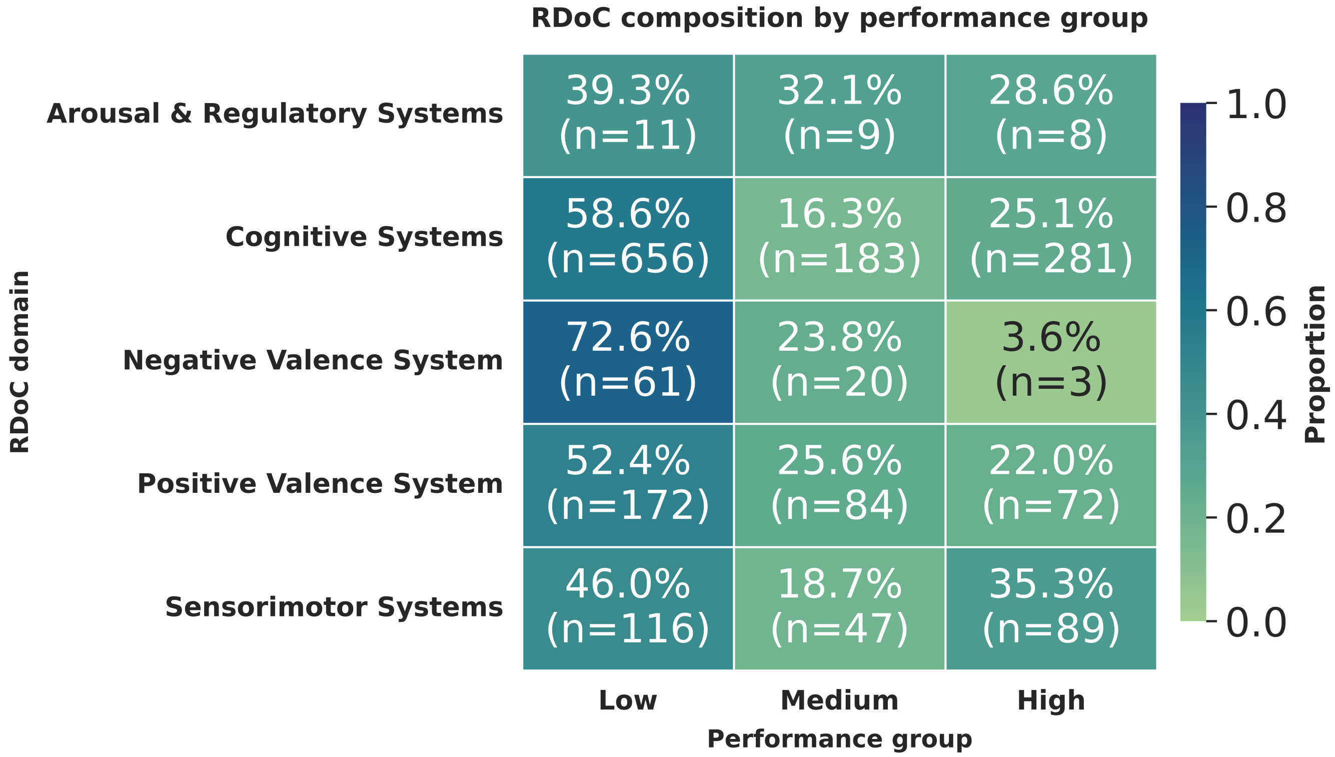


**Fig. 27. Performance-group composition across RDoC domains.** Composition of performance groups across RDoC domains, showing the proportion of runs classified as [High/Medium/Low] performance for each domain. Bars (or heatmap cells) indicate percentages within domain after normalization to 100%; overlaid annotations report corresponding sample counts (n). Group thresholds were defined using quantile-based cutoffs – the top quartile and bottom quartile as high performance and low performance and the samples between as medium – applied consistently across all runs. Domains with small n should be interpreted cautiously because proportion estimates are less stable.

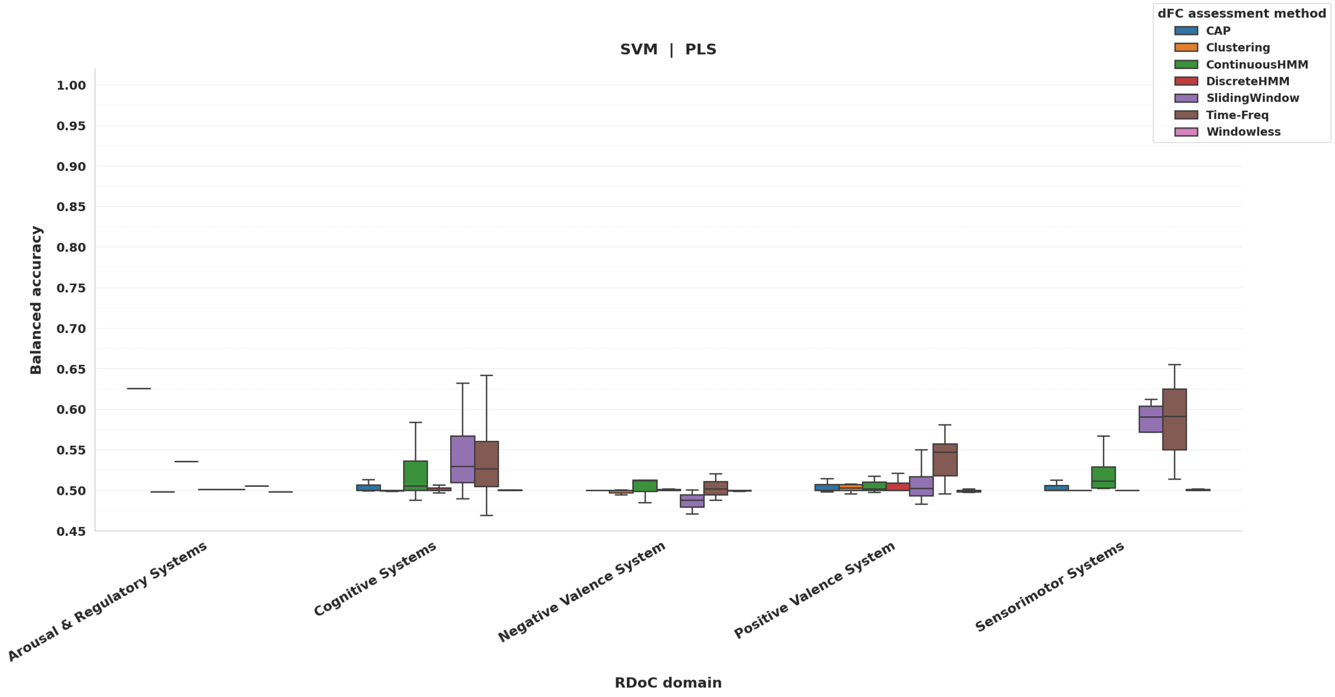


**Fig. 28. Domain-wise performance distributions with method-level grouping (using SVM on PLS-transformed features).** Balanced accuracy distribution is shown across RDoC domains, with dFC assessment methods displayed as grouped boxplots (color-coded hue) within each domain. Boxes indicate median and interquartile range, whiskers represent distribution spread (outliers not shown).

#### Three-Dimensional Visualization of Embedded dFC Features for Representative Scans

To illustrate the representational geometry of dFC features, we visualized the Principal Component Analysis (PCA), Partial Least Square (PLS), and Laplacian Eigenmaps (LE) embeddings for each dFC methodology using one representative scan from `EXP.12`, and simulated `EXP.S.29`. Each plot displays the first three components from embedded dFC features of the subject, with samples colored by condition (rest vs. task). For state-based methods, the embedding is applied to state probability features, while for state-free methods the embedding is applied to dFC values.

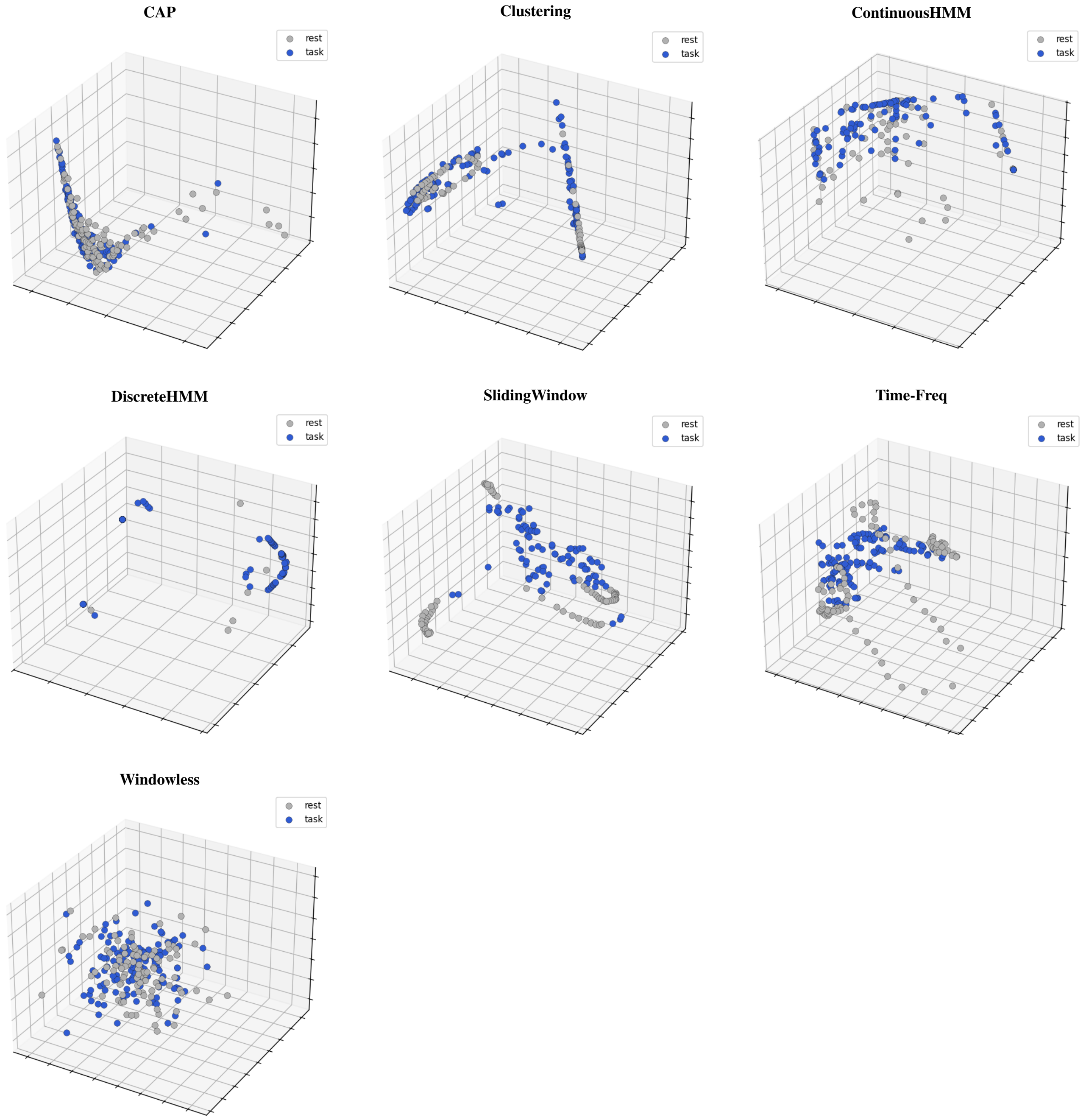


**Fig. 29. Three-dimensional visualization of PCA embeddings from `EXP.12`.** Three-dimensional visualization of PCA-transformed dFC features for a representative subject from `EXP.12`. Each panel corresponds to one dFC methodology. Rest and task-present time points are color-coded, illustrating task-rest separability within the reduced feature space. For state-based methods, the PCA is applied to state probability features, allowing visualization of samples in 3D space. These examples highlight inter-method variability in the degree to which dFC features in 3D PCA embedding detect task engagement at single-TR resolution.

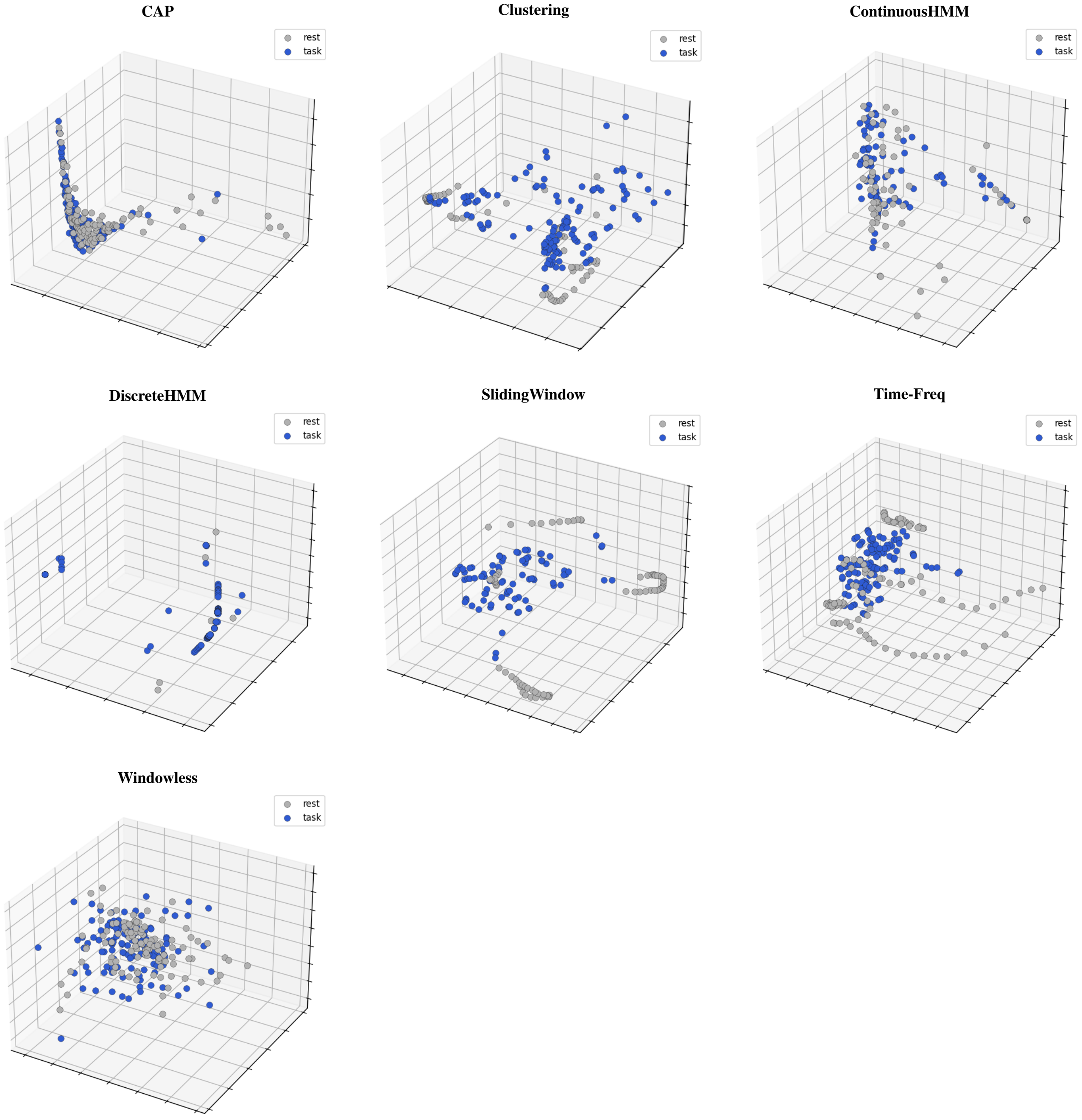


**Fig. 30. Three-dimensional visualization of PLS embeddings from `EXP.12`.** Three-dimensional visualization of PLS-transformed dFC features for a representative subject from `EXP.12`. Each panel corresponds to one dFC methodology. Rest and task-present time points are color-coded, illustrating task-rest separability within the reduced feature space. For state-based methods, the PLS is applied to state probability features, allowing visualization of samples in 3D space. These examples highlight inter-method variability in the degree to which dFC features in 3D PLS embedding detect task engagement at single-TR resolution.

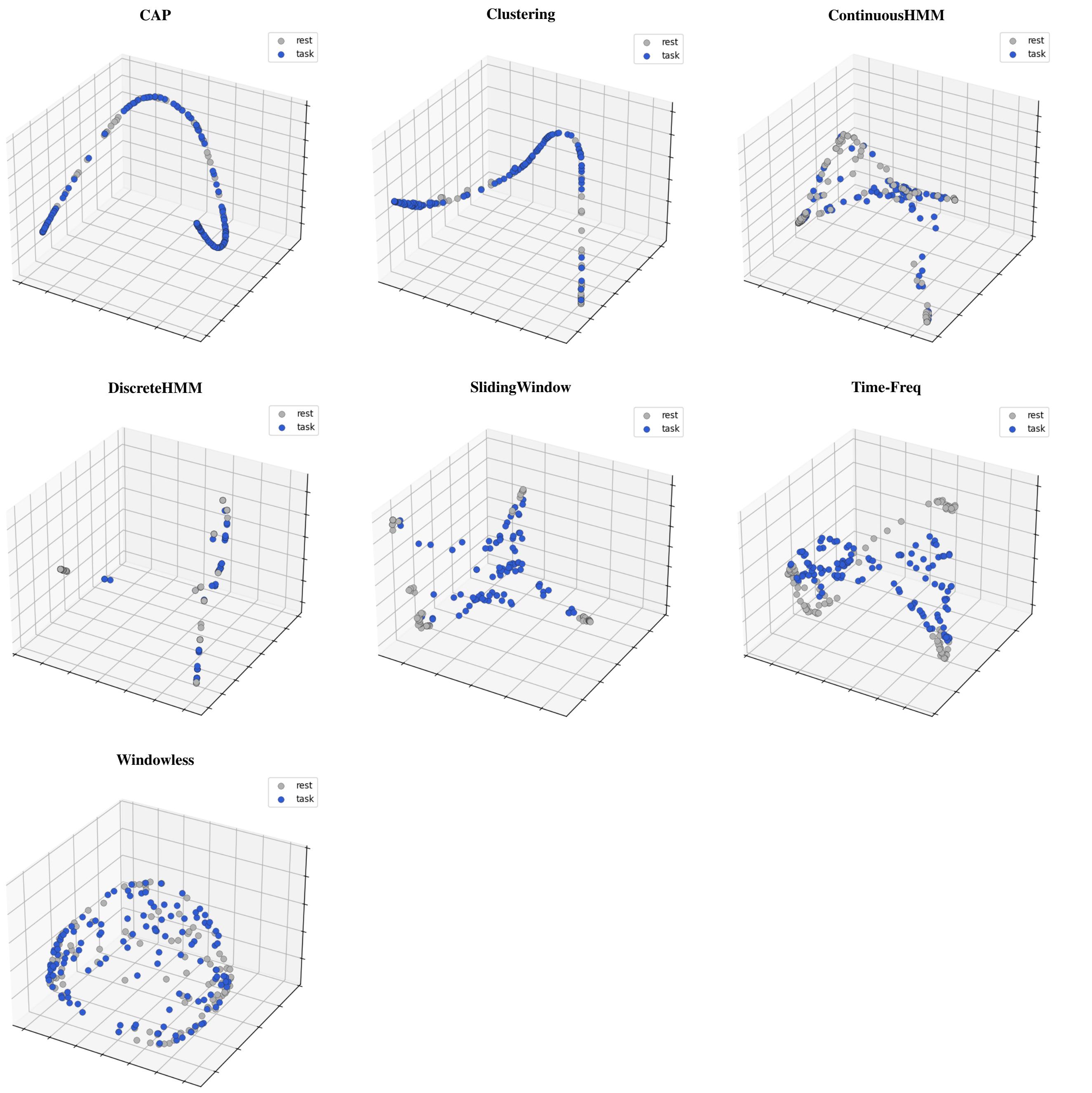


**Fig. 31. Three-dimensional visualization of LE embeddings from `EXP.12`.** Three-dimensional visualization of LE-transformed dFC features for a representative subject from `EXP.12`. Each panel corresponds to one dFC methodology. Rest and task-present time points are color-coded, illustrating task-rest separability within the reduced feature space. For state-based methods, the LE is applied to state probability features, allowing visualization of samples in 3D space. These examples highlight inter-method variability in the degree to which dFC features in 3D LE embedding detect task engagement at single-TR resolution.

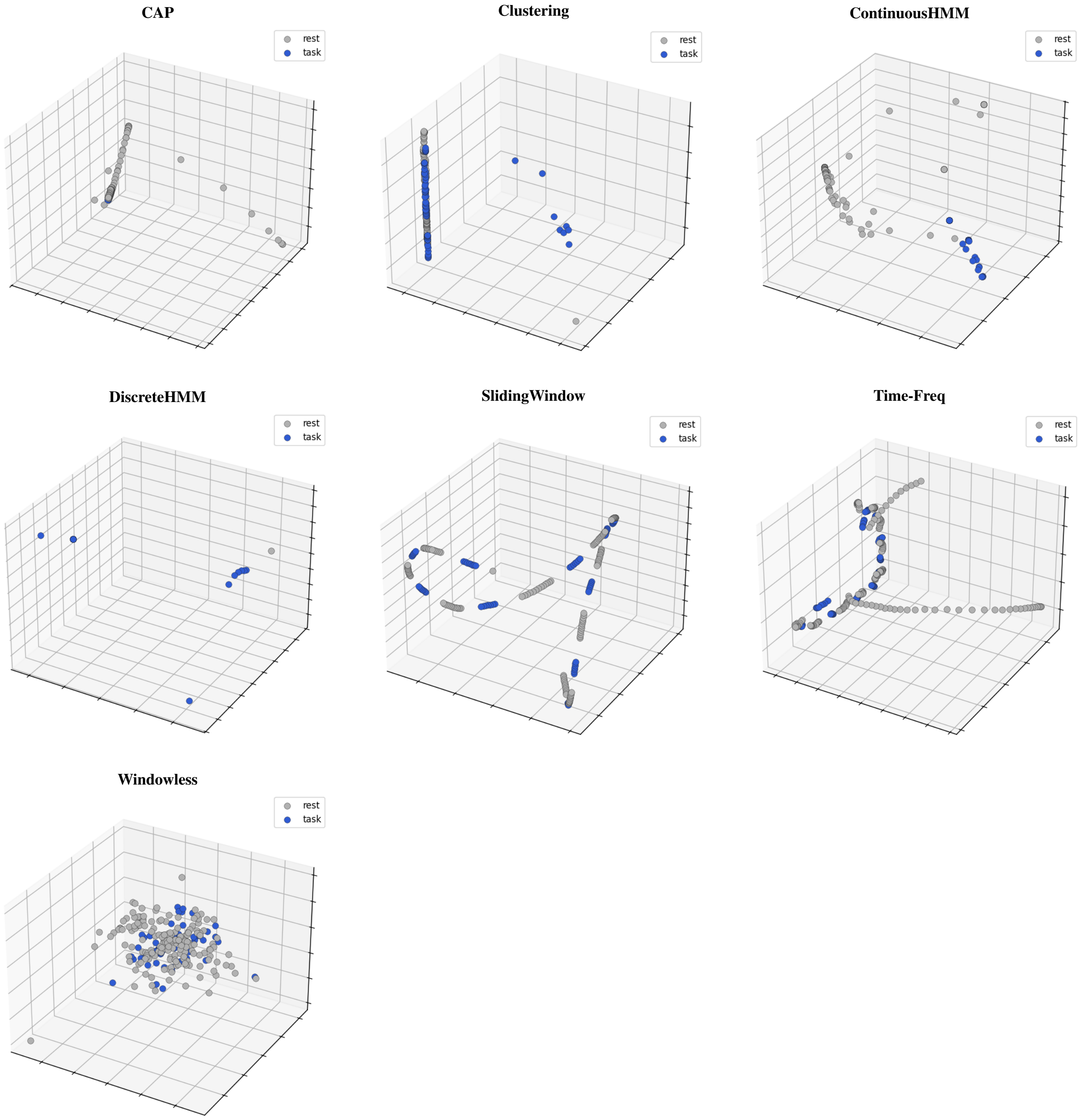


**Fig. 32. Three-dimensional visualization of PCA embeddings from simulated experiment `EXP.S.29`.** Three-dimensional visualization of PCA-transformed dFC features for a representative subject from `EXP.S.29`. Each panel corresponds to one dFC methodology. Rest and task-present time points are color-coded, illustrating the separability of rest and task-present states within the reduced feature space.

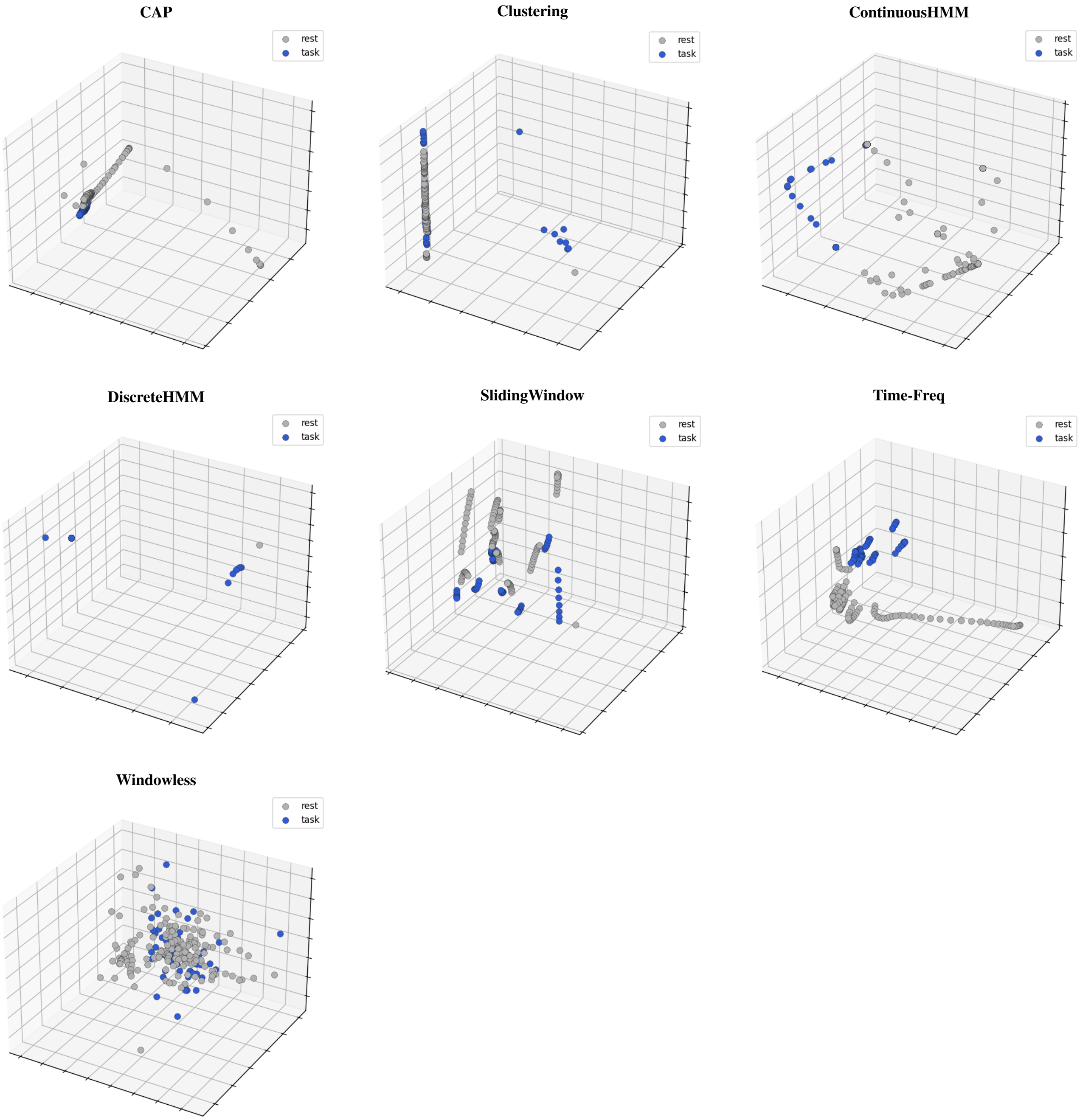


**Fig. 33. Three-dimensional visualization of PLS embeddings from simulated experiment `EXP.S.29`.** Three-dimensional visualization of PLS-transformed dFC features for a representative subject from `EXP.S.29`. Each panel corresponds to one dFC methodology. Rest and task-present time points are color-coded, illustrating the separability of rest and task-present states within the reduced feature space.

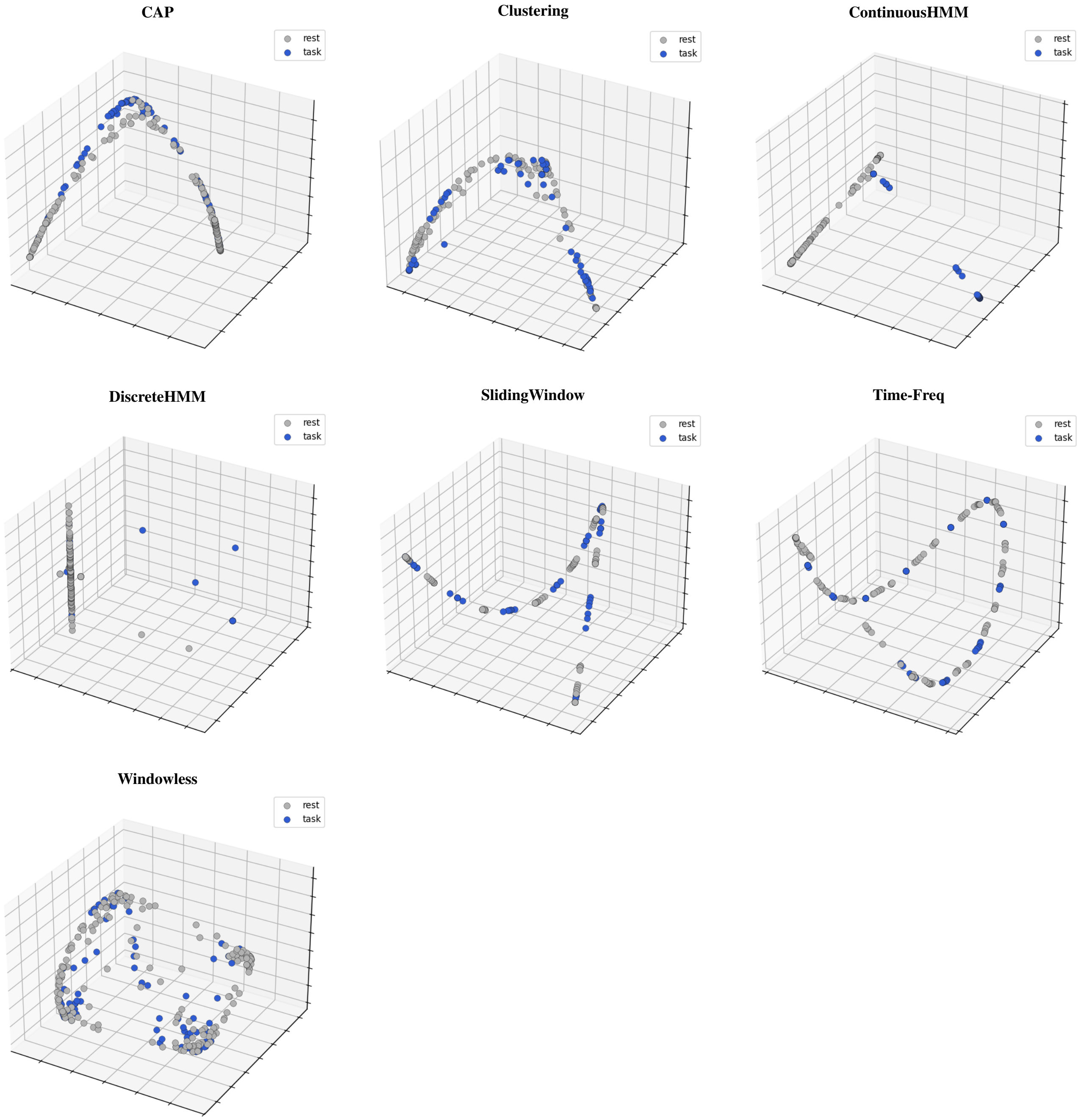


**Fig. 34. Three-dimensional visualization of LE embeddings from simulated experiment `EXP.S.29`.** Three-dimensional visualization of LE-transformed dFC features for a representative subject from `EXP.S.29`. Each panel corresponds to one dFC methodology. Rest and task-present time points are color-coded, illustrating the separability of rest and task-present states within the reduced feature space.

#### Feature-Sample Matrices Across Experimental and Simulated Data

To illustrate how dFC features reflect task-related variations across experiments, we present representative sample–feature matrices from two experimental datasets: one with strong decoding performance (`EXP.12` with Sliding Window) and one with weak decoding performance (`EXP.7` with Sliding Window), as well as a simulated experiment (`EXP.S.29`). For each experiment, training and test sets are visualized to highlight differences in feature organization.

In `EXP.12`, task and rest samples exhibit distinct and structured patterns, indicating that dFC captures task-related variability. In contrast, in `EXP.7`, task and rest samples show similar and overlapping patterns, consistent with weaker decoding performance.

In the simulated experiment, a clear separation between task and rest samples emerges, demonstrating that under idealized conditions with high signal-to-noise ratio and controlled task design, dFC can robustly capture task-related dynamics.

Together, these visualizations reinforce that the detectability of task-related transitions in dFC depends jointly on methodology, experimental design, and data quality.

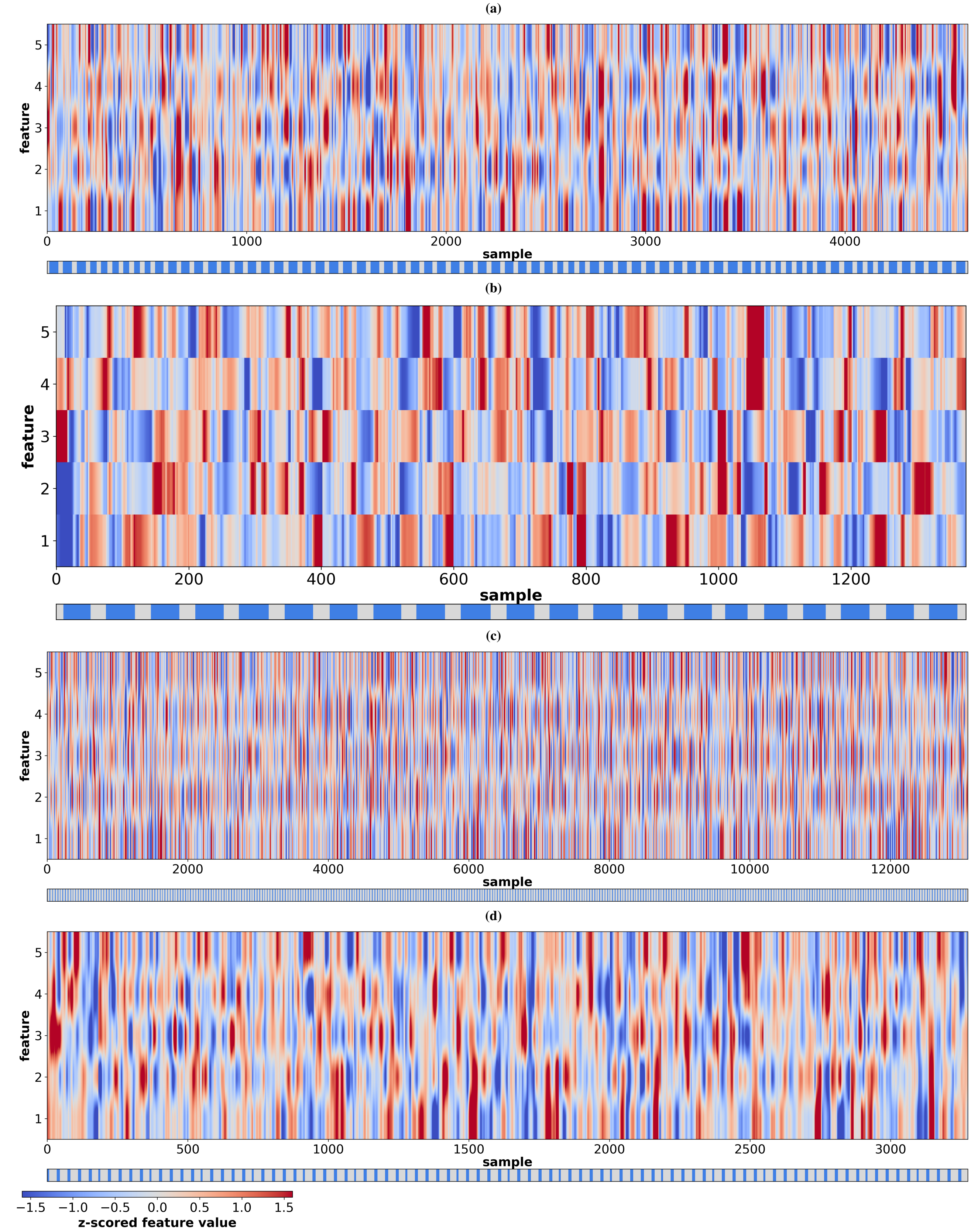


**Fig. 35. Dynamic functional connectivity feature matrices for representative experiments (`EXP.12` and `EXP.7`).** Sample-by-feature heatmaps of dFC features obtained using the Sliding Window method and transformed using PLS. **(a–b)** Training and test samples from `EXP.12`. **(c–d)** Training and test samples from `EXP.7`. Columns represent samples (time points across all subjects, shown in their original temporal order), and rows correspond to features. The color bar beneath each matrix indicates ground-truth labels (task-present in blue, rest in grey).

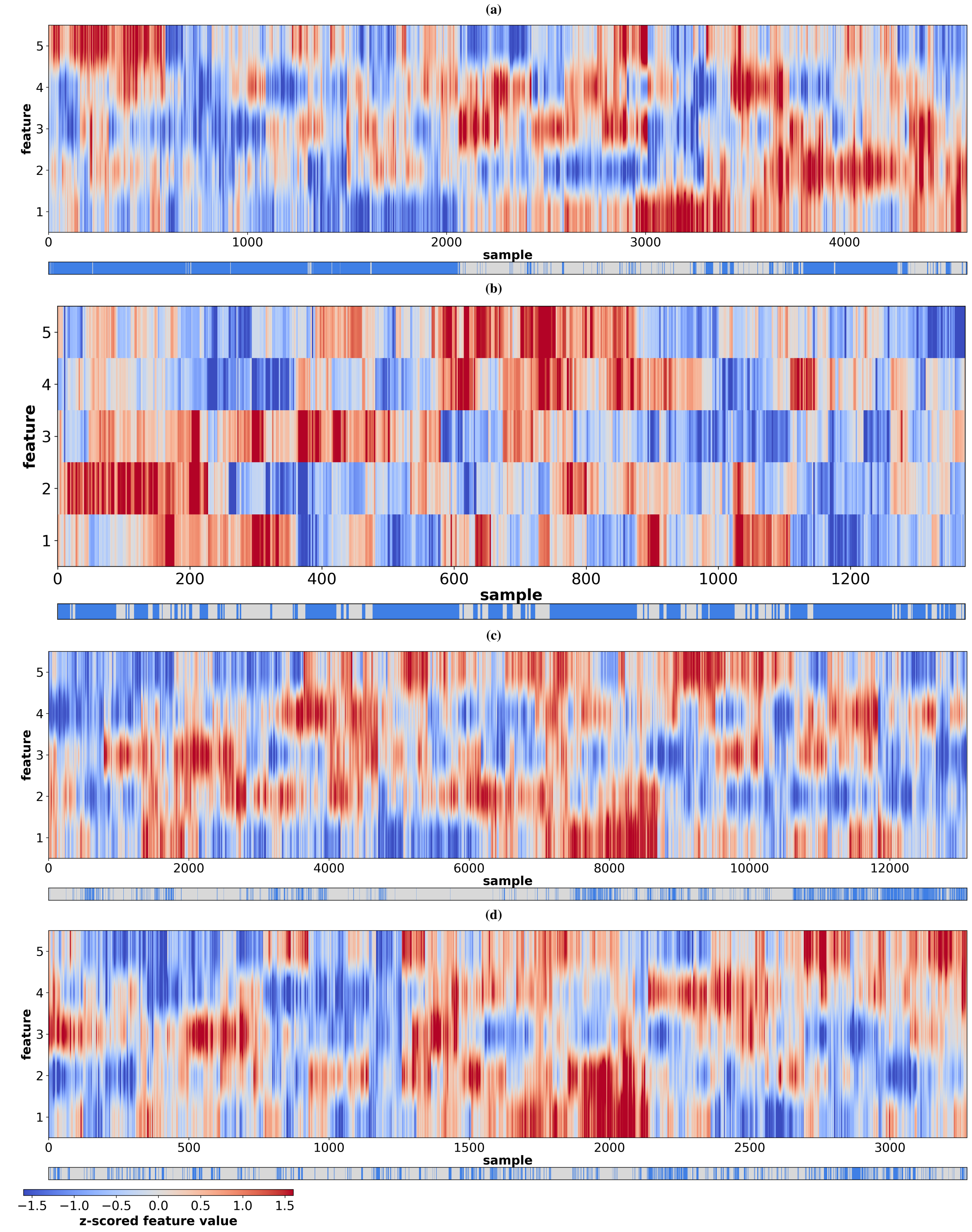


**Fig. 36. Dynamic functional connectivity feature matrices for representative high- and low-performing experiments.** This figure provides a qualitative comparison of how dFC features differ between task and rest across experiment–method combinations. **(a–b)** PLS-transformed Sliding Window features for training and test samples from `EXP.12` (high decoding performance). **(c–d)** PLS-transformed Sliding Window features for training and test samples from `EXP.7` (low decoding performance). Columns correspond to samples (time points across all subjects), and rows correspond to features. Samples are ordered using hierarchical clustering based on cosine similarity. The color bar beneath each matrix indicates ground-truth labels (task-present in blue, rest in grey). In `EXP.12`, samples with similar features tend to belong to the same class, forming coherent groupings that reflect distinct task-related patterns, facilitating task-rest detectability. In contrast, in `EXP.7`, samples with similar features are more intermixed across classes, indicating weaker differentiation between task and rest states.

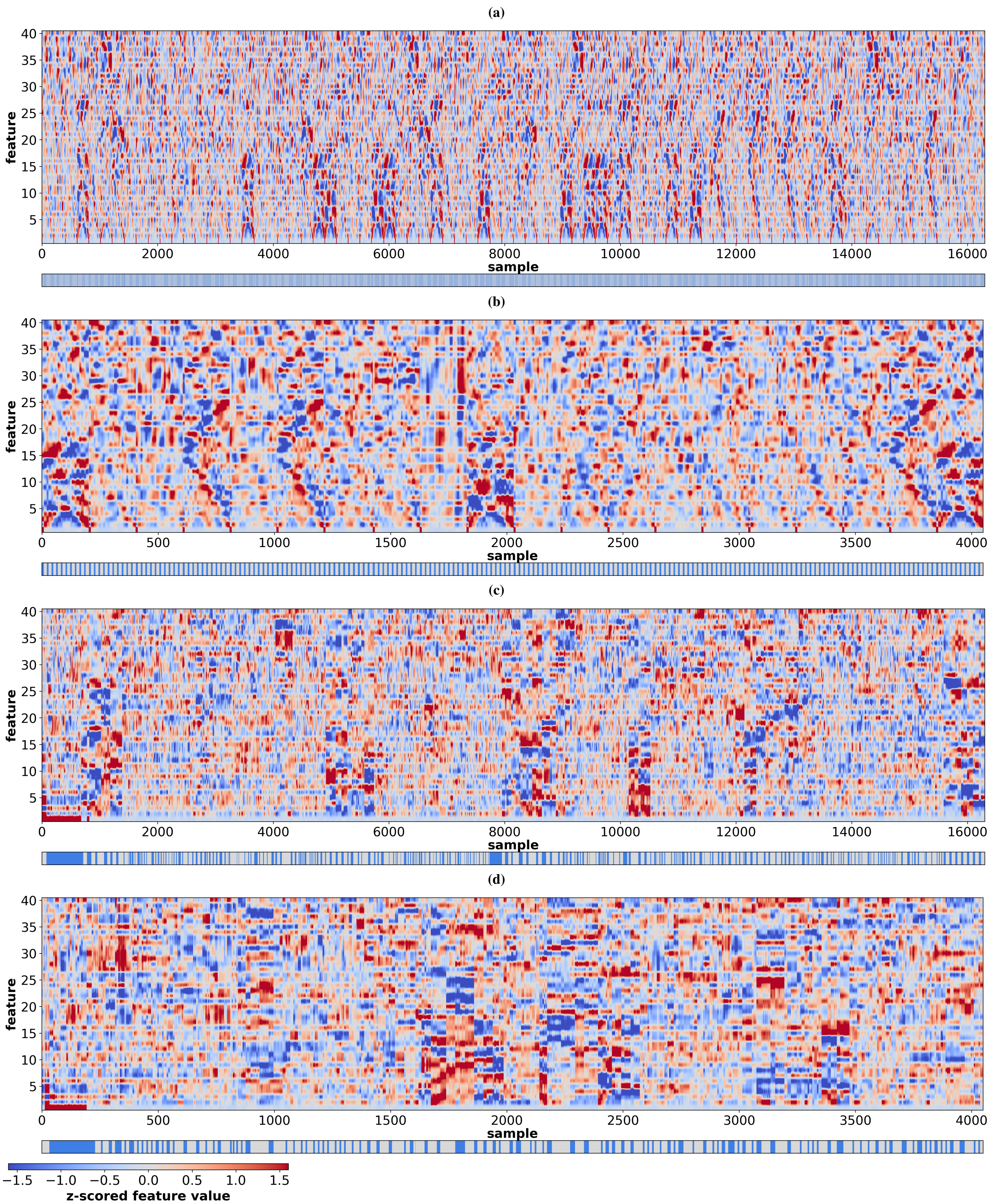


**Fig. 37. Dynamic functional connectivity feature matrices for a representative simulated experiment (`EXP.S.29`).** Sample-by-feature heatmaps of dFC features obtained using the Sliding Window method and transformed using PLS. Columns represent samples (time points across all subjects), and rows correspond to features. The color bar beneath each matrix indicates ground-truth labels (task-present in blue, rest in grey). **(a–b)** Training and test samples in their original temporal order. **(c–d)** The same samples reordered so that those with similar feature patterns are placed next to one another, rather than in temporal order.

#### Experimental Data Classification Performance Results

This section provides extended quantitative results for the classification analyses presented in the main text. The goal is to offer detailed experiment-by-method values and complementary comparisons across different dimensionality reduction and classifier configurations. While the main paper highlights overall trends in dFC performance, the figures below allow direct inspection of exact accuracies for each experiment-method pair. Together, these analyses further confirm the main finding that state-free methodologies such as Sliding Window and Time-Frequency generally outperform state-based ones, and show that performance strongly depends on both the chosen dimensionality reduction technique and the classifier type. *Note: For experiments with multiple runs, all reported values correspond to the best-performing run, unless stated otherwise.*

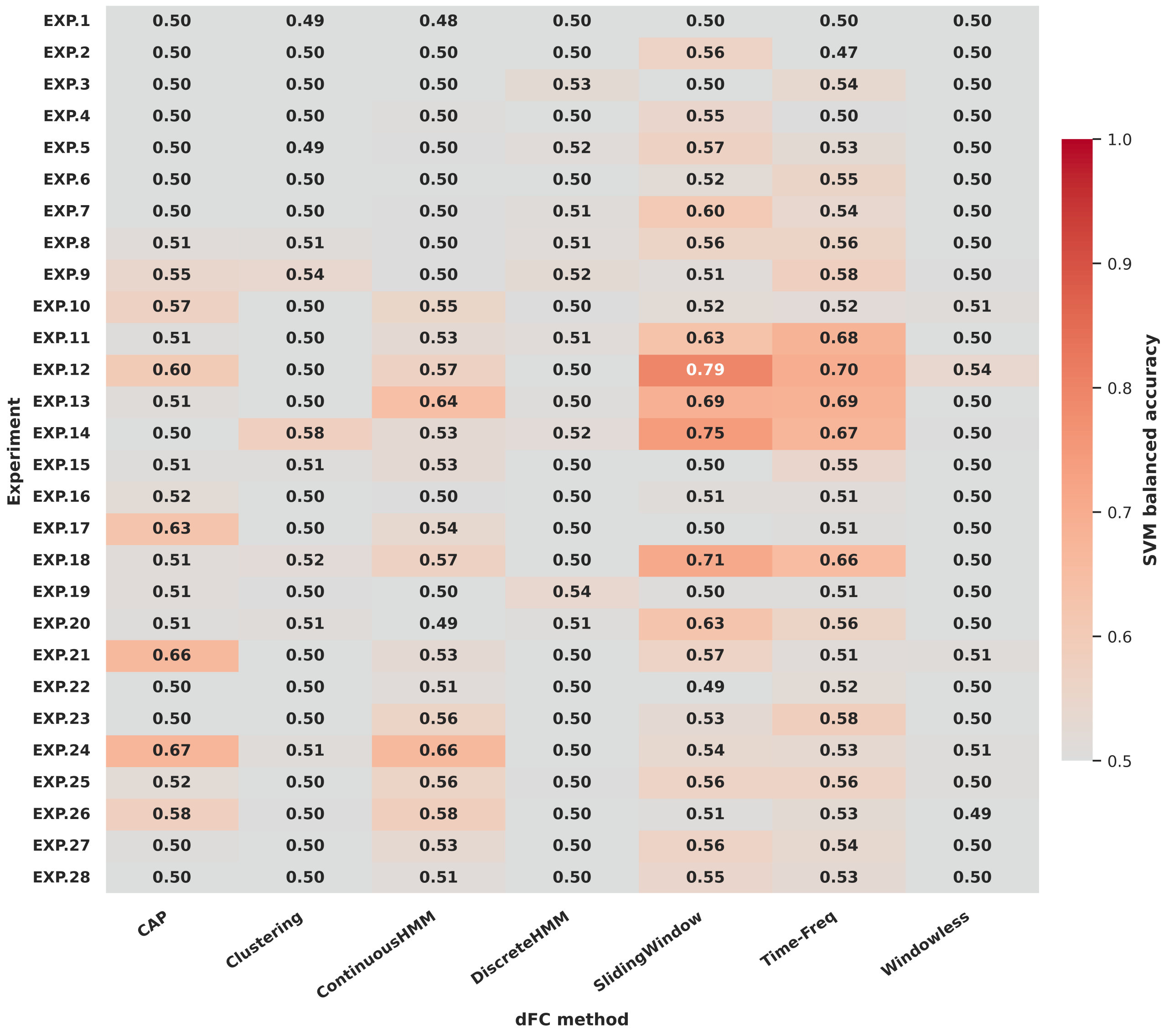


**Fig. 38. Balanced accuracy across experiments and dFC methods (PLS-SVM configuration).** Heatmap showing classification balanced accuracy for each experiment-method pair using PLS-reduced features and a SVM classifier. Values are annotated on the heatmap to facilitate direct comparison across methods and experiments.

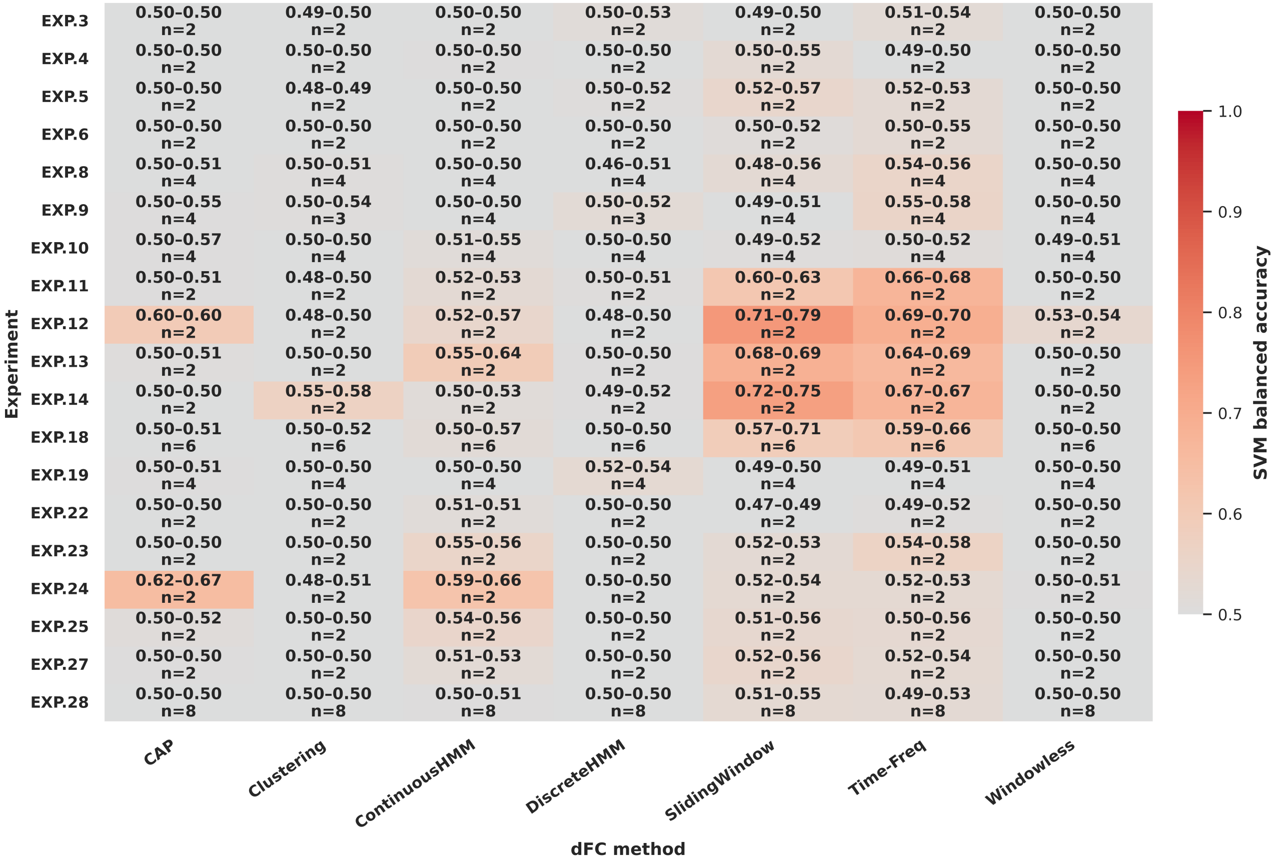


| Experiment | CAP | Clustering | ContinuousHMM | DiscreteHMM | SlidingWindow | Time-Freq | Windowless |
|---|---|---|---|---|---|---|---|
| EXP.3 | 0.50-0.50 n=2 | 0.49-0.50 n=2 | 0.50-0.50 n=2 | 0.50-0.53 n=2 | 0.49-0.50 n=2 | 0.51-0.54 n=2 | 0.50-0.50 n=2 |
| EXP.4 | 0.50-0.50 n=2 | 0.50-0.50 n=2 | 0.50-0.50 n=2 | 0.50-0.50 n=2 | 0.50-0.55 n=2 | 0.49-0.50 n=2 | 0.50-0.50 n=2 |
| EXP.5 | 0.50-0.50 n=2 | 0.48-0.49 n=2 | 0.50-0.50 n=2 | 0.50-0.52 n=2 | 0.52-0.57 n=2 | 0.52-0.53 n=2 | 0.50-0.50 n=2 |
| EXP.6 | 0.50-0.50 n=2 | 0.50-0.50 n=2 | 0.50-0.50 n=2 | 0.50-0.50 n=2 | 0.50-0.52 n=2 | 0.50-0.55 n=2 | 0.50-0.50 n=2 |
| EXP.8 | 0.50-0.51 n=4 | 0.50-0.51 n=4 | 0.50-0.50 n=4 | 0.46-0.51 n=4 | 0.48-0.56 n=4 | 0.54-0.56 n=4 | 0.50-0.50 n=4 |
| EXP.9 | 0.50-0.55 n=4 | 0.50-0.54 n=3 | 0.50-0.50 n=4 | 0.50-0.52 n=3 | 0.49-0.51 n=4 | 0.55-0.58 n=4 | 0.50-0.50 n=4 |
| EXP.10 | 0.50-0.57 n=4 | 0.50-0.50 n=4 | 0.51-0.55 n=4 | 0.50-0.50 n=4 | 0.49-0.52 n=4 | 0.50-0.52 n=4 | 0.49-0.51 n=4 |
| EXP.11 | 0.50-0.51 n=2 | 0.48-0.50 n=2 | 0.52-0.53 n=2 | 0.50-0.51 n=2 | 0.60-0.63 n=2 | 0.66-0.68 n=2 | 0.50-0.50 n=2 |
| EXP.12 | 0.60-0.60 n=2 | 0.48-0.50 n=2 | 0.52-0.57 n=2 | 0.48-0.50 n=2 | 0.71-0.79 n=2 | 0.69-0.70 n=2 | 0.53-0.54 n=2 |
| EXP.13 | 0.50-0.51 n=2 | 0.50-0.50 n=2 | 0.55-0.64 n=2 | 0.50-0.50 n=2 | 0.68-0.69 n=2 | 0.64-0.69 n=2 | 0.50-0.50 n=2 |
| EXP.14 | 0.50-0.50 n=2 | 0.55-0.58 n=2 | 0.50-0.53 n=2 | 0.49-0.52 n=2 | 0.72-0.75 n=2 | 0.67-0.67 n=2 | 0.50-0.50 n=2 |
| EXP.18 | 0.50-0.51 n=6 | 0.50-0.52 n=6 | 0.50-0.57 n=6 | 0.50-0.50 n=6 | 0.57-0.71 n=6 | 0.59-0.66 n=6 | 0.50-0.50 n=6 |
| EXP.19 | 0.50-0.51 n=4 | 0.50-0.50 n=4 | 0.50-0.50 n=4 | 0.52-0.54 n=4 | 0.49-0.50 n=4 | 0.49-0.51 n=4 | 0.50-0.50 n=4 |
| EXP.22 | 0.50-0.50 n=2 | 0.50-0.50 n=2 | 0.51-0.51 n=2 | 0.50-0.50 n=2 | 0.47-0.49 n=2 | 0.49-0.52 n=2 | 0.50-0.50 n=2 |
| EXP.23 | 0.50-0.50 n=2 | 0.50-0.50 n=2 | 0.55-0.56 n=2 | 0.50-0.50 n=2 | 0.52-0.53 n=2 | 0.54-0.58 n=2 | 0.50-0.50 n=2 |
| EXP.24 | 0.62-0.67 n=2 | 0.48-0.51 n=2 | 0.59-0.66 n=2 | 0.50-0.50 n=2 | 0.52-0.54 n=2 | 0.52-0.53 n=2 | 0.50-0.51 n=2 |
| EXP.25 | 0.50-0.52 n=2 | 0.50-0.50 n=2 | 0.54-0.56 n=2 | 0.50-0.50 n=2 | 0.51-0.56 n=2 | 0.50-0.56 n=2 | 0.50-0.50 n=2 |
| EXP.27 | 0.50-0.50 n=2 | 0.50-0.50 n=2 | 0.51-0.53 n=2 | 0.50-0.50 n=2 | 0.52-0.56 n=2 | 0.52-0.54 n=2 | 0.50-0.50 n=2 |
| EXP.28 | 0.50-0.50 n=8 | 0.50-0.50 n=8 | 0.50-0.51 n=8 | 0.50-0.50 n=8 | 0.51-0.55 n=8 | 0.49-0.53 n=8 | 0.50-0.50 n=8 |

**Fig. 39. Across-run classification performance.** Heatmap summarizing SVM-based test balanced accuracies using PLS-transformed dFC features for experiments with multiple runs. Each cell corresponds to an experiment-method pair, with color representing the median accuracy across runs, and in-cell annotations showing the minimum and maximum values (min-max) and the number of available runs (n). This dual encoding visualizes both central tendency and run-to-run stability: narrow min-max ranges indicate consistent performance, while wider ranges highlight variability across runs. High, warm-colored cells with tight min-max ranges (and reasonable n) indicate robust, high performance across runs; warm colors with wide ranges flag instability; cool colors indicate near-chance medians regardless of spread.

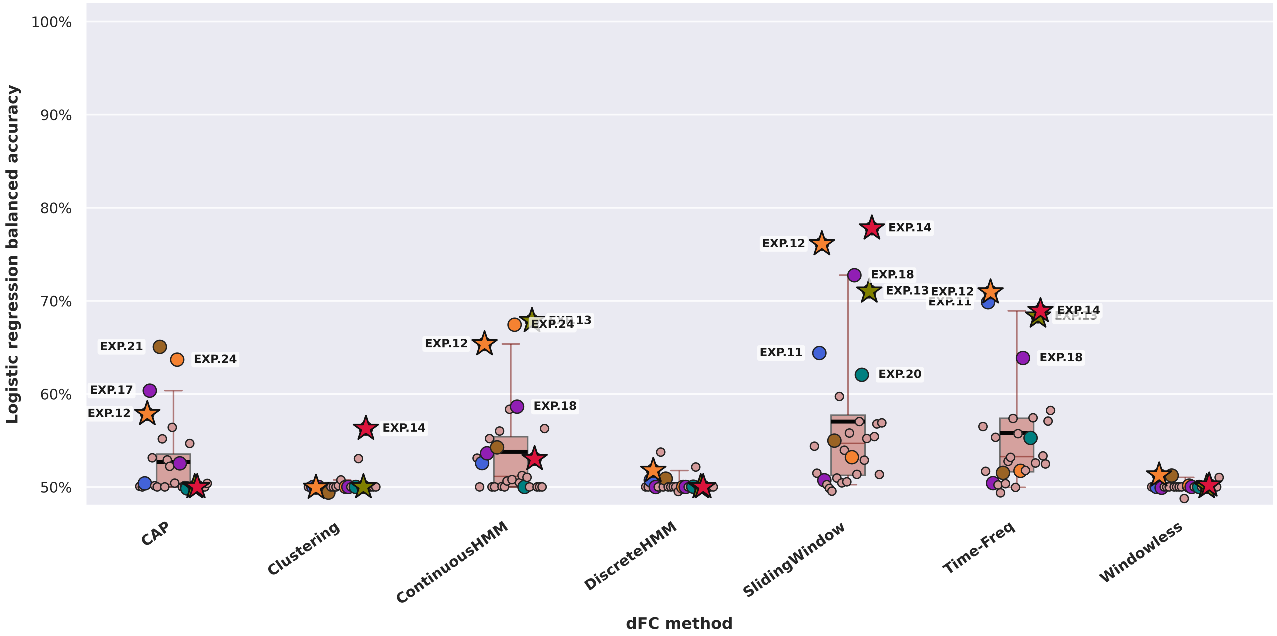


**Fig. 40. Classification performance using PLS features with logistic regression.** Strip and box plots show balanced accuracies across experiments and dFC methods using PLS-reduced features and logistic regression as classifier. Performance is overall very similar to PLS-SVM; however, some experiment rankings are different.

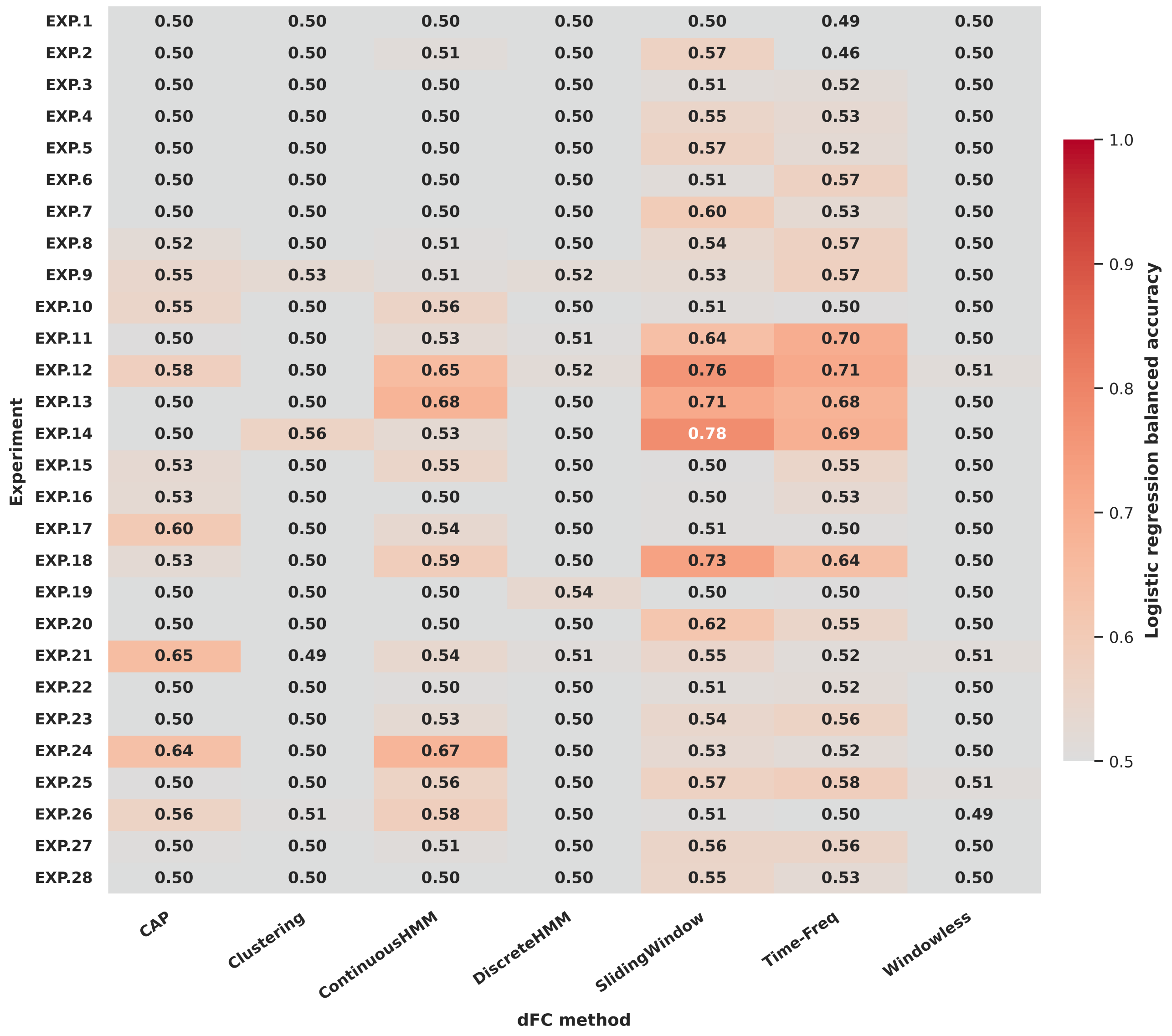


**Fig. 41. Balanced accuracy across experiments and methods (PLS-logistic regression configuration).** Same configuration as Figure40 but presented as a heatmap for clarity.

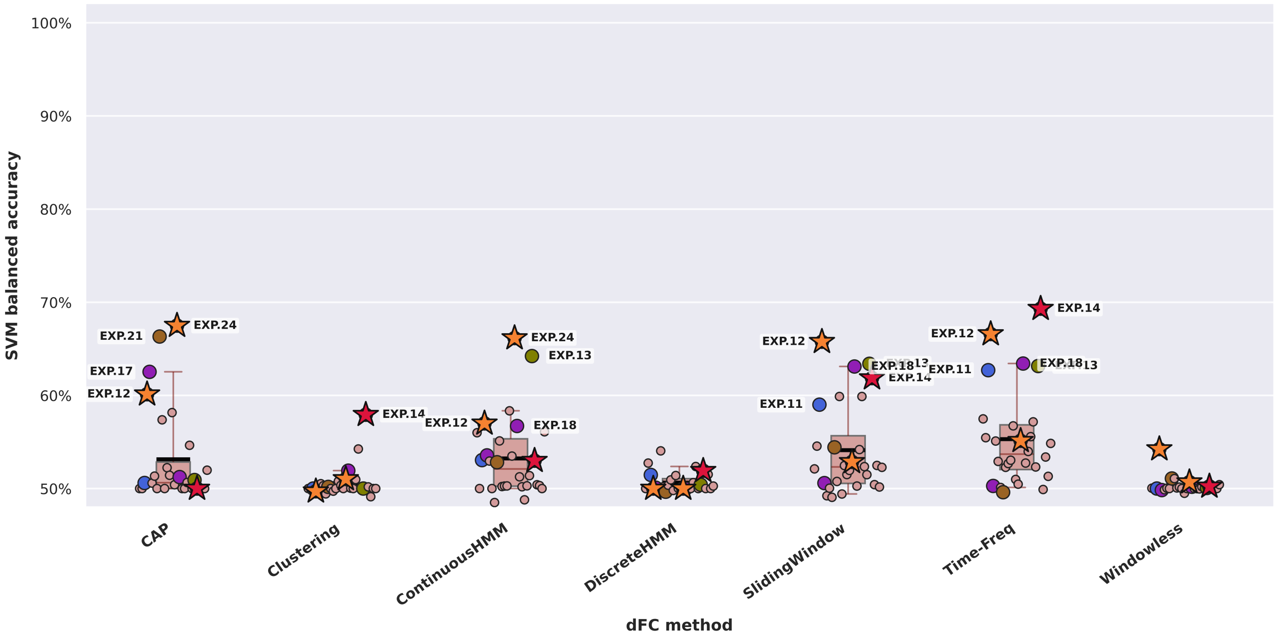


**Fig. 42. Classification performance using PCA-reduced features with SVM.** Balanced accuracy distributions obtained using PCA for dimensionality reduction and an SVM classifier. Overall performance is lower compared to PLS-based analyses, with no values going above 70%. Note: State-based methods retain the same values as in the PLS-based analyses since their four-dimensional feature spaces are not subjected to dimensionality reduction.

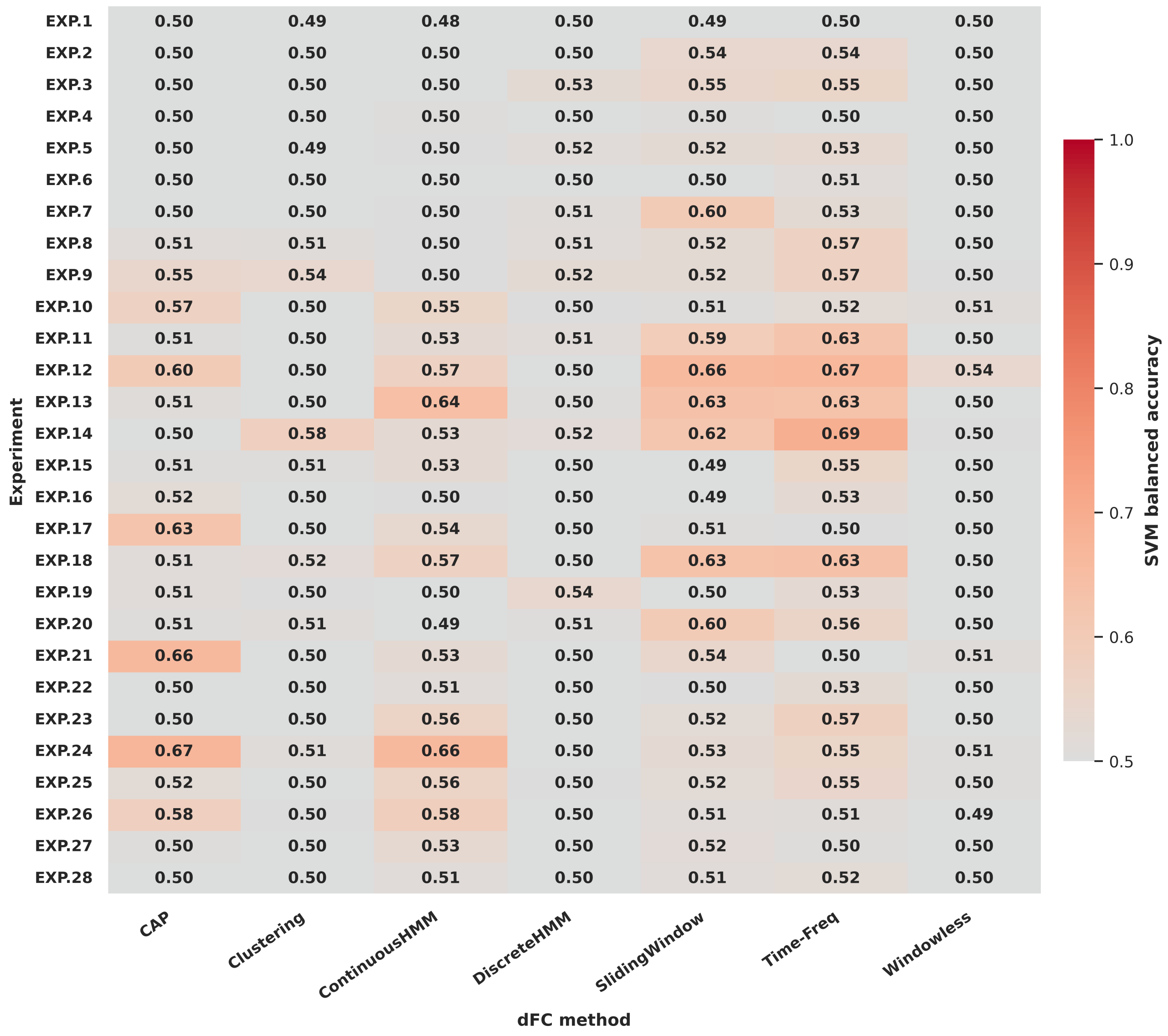


**Fig. 43. Balanced accuracy across experiments and methods (PCA-SVM configuration).** Heatmap version of the PCA-SVM results.

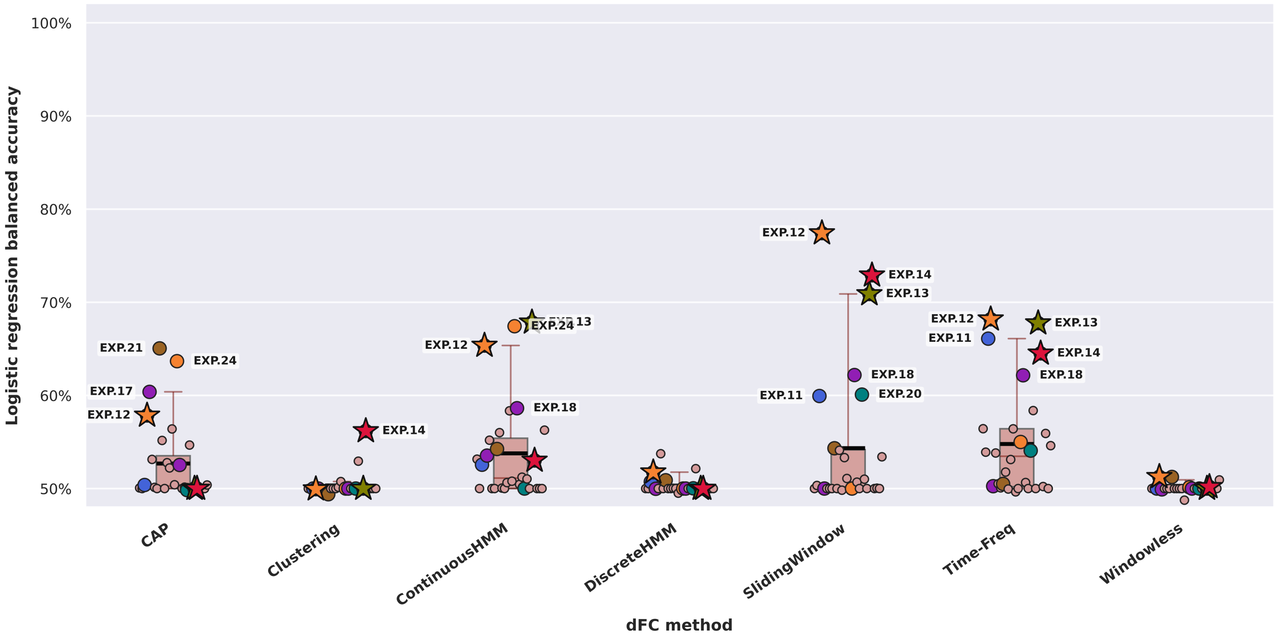


**Fig. 44. Classification performance using PCA features with logistic regression.** Balanced accuracies across experiments and methods using PCA-reduced features and a logistic regression classifier. The performance was overall slightly lower but very close to PLS-SVM results. This configuration outperforms PCA-SVM (e.g., `EXP.13`: 0.77 with SW). Note: State-based methods retain the same values as in the PLS-based analyses since their four-dimensional feature spaces are not subjected to dimensionality reduction.

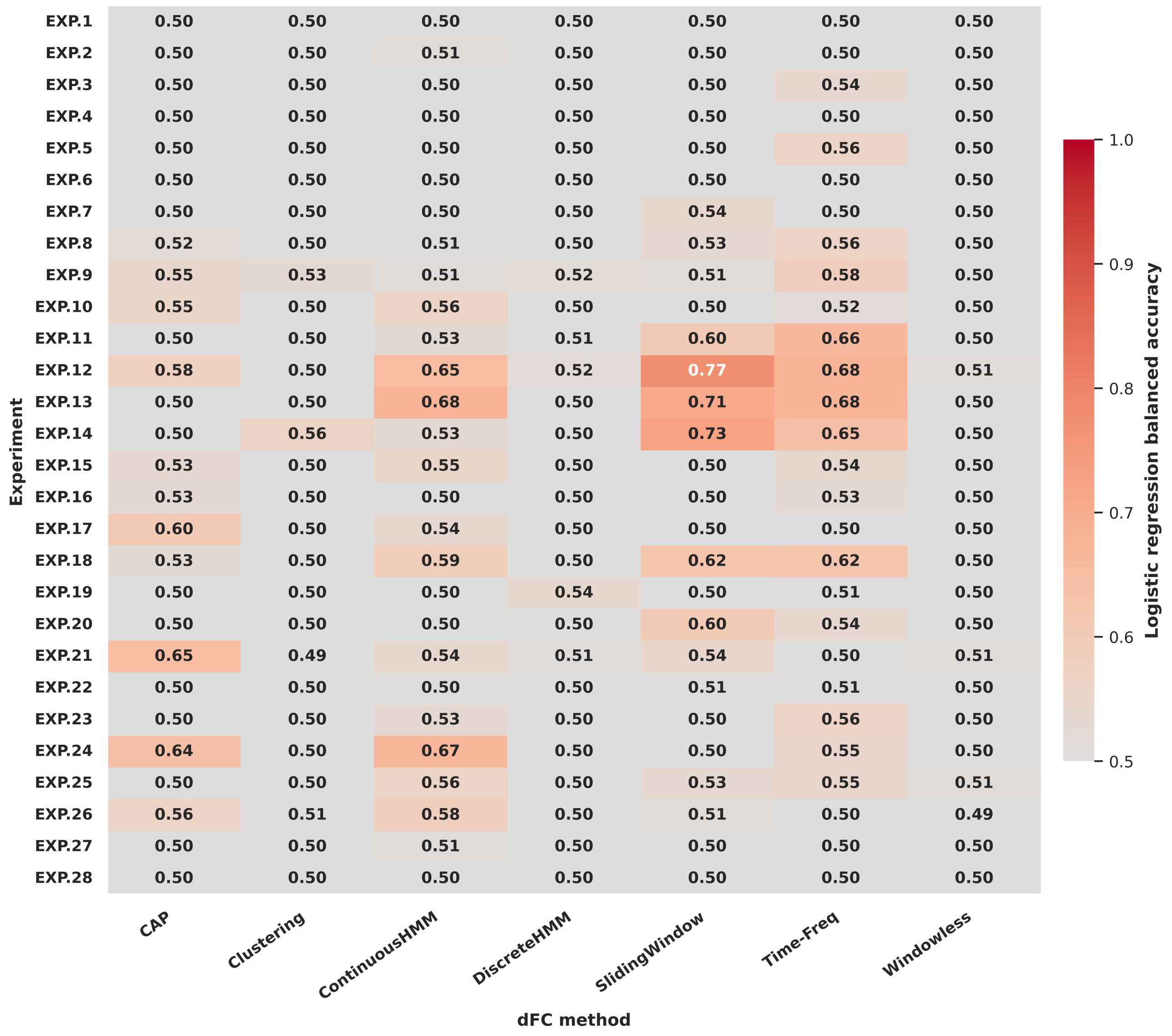


**Fig. 45. Balanced accuracy across experiments and methods (PCA-logistic regression configuration).** Heatmap complementing Figure44.

#### Simulated Data Classification Performance Results

This section presents classification results for simulated experiments, complementing the experimental-data analyses shown in the main text. By examining performance across combinations of dimensionality-reduction and classifier models, we assess how methodological choices affect dFC-based cognitive-state prediction under idealized conditions. Unlike experimental data, simulated experiments feature precisely controlled experimental designs, enabling clearer interpretation of method sensitivity. Across all configurations, Co-Activation Patterns (CAP) and Continuous Hidden Markov Models (CHMM) achieve the highest balanced accuracies, often approaching ceiling performance, while Sliding Window and Time-Frequency perform comparably lower. PCA sometimes outperforms PLS for state-free methods with the SVM classifier, while performing much lower than PLS when applied with logistic regression. Similar to experimental data, SVM classification generally yields higher accuracy than logistic regression.

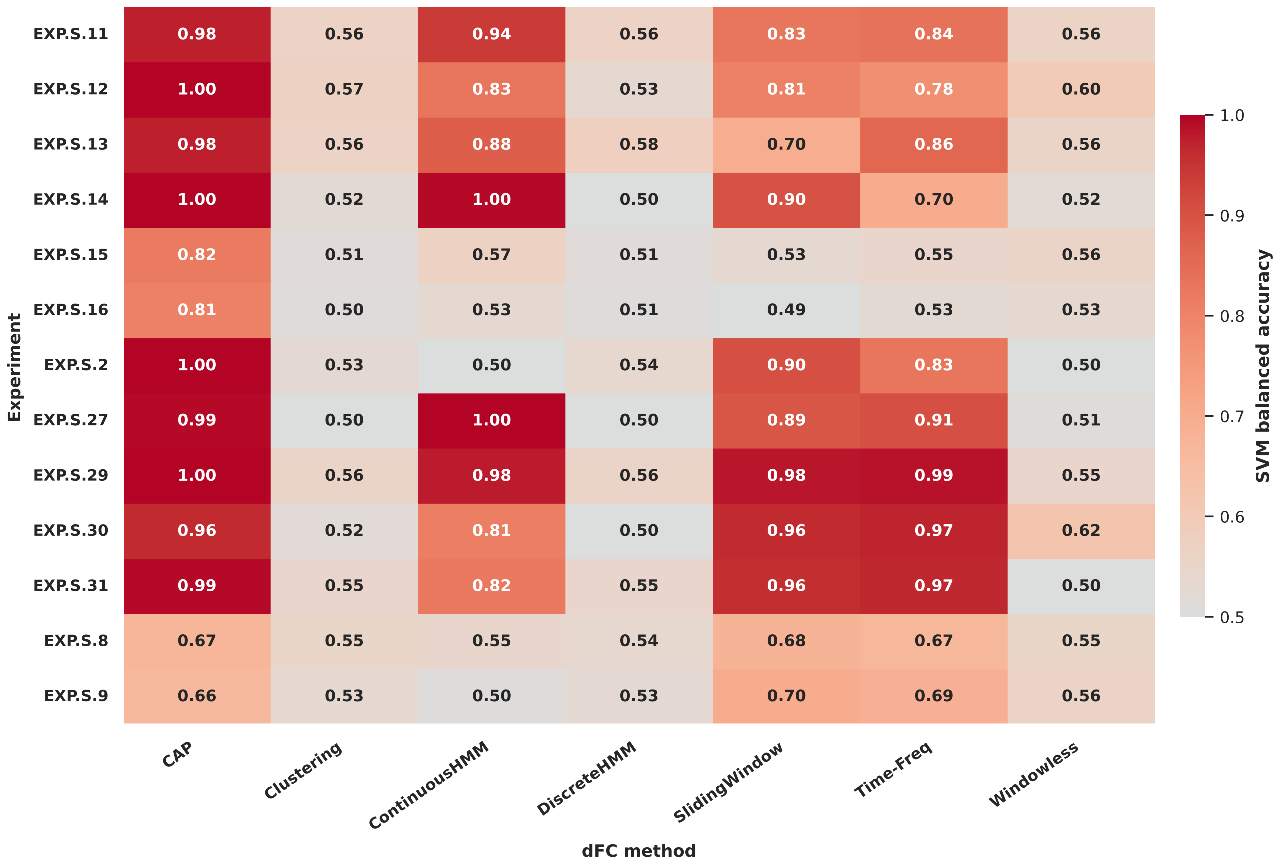


**Fig. 46. PLS-SVM classification accuracies across simulated experiments.** Heatmap of balanced accuracies for each experiment-method pair using PLS reduction and SVM classification.

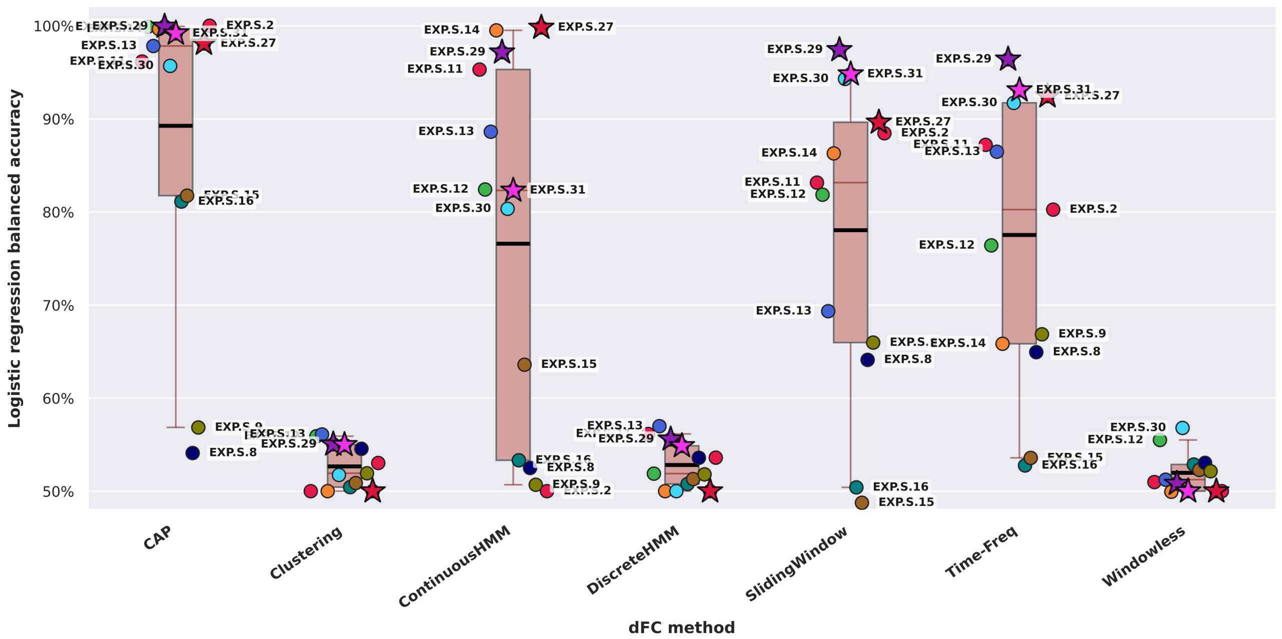


**Fig. 47. PLS-Logistic Regression classification performance for simulated experiments.** Boxplots and stripplots showing balanced accuracies for all dFC methods.

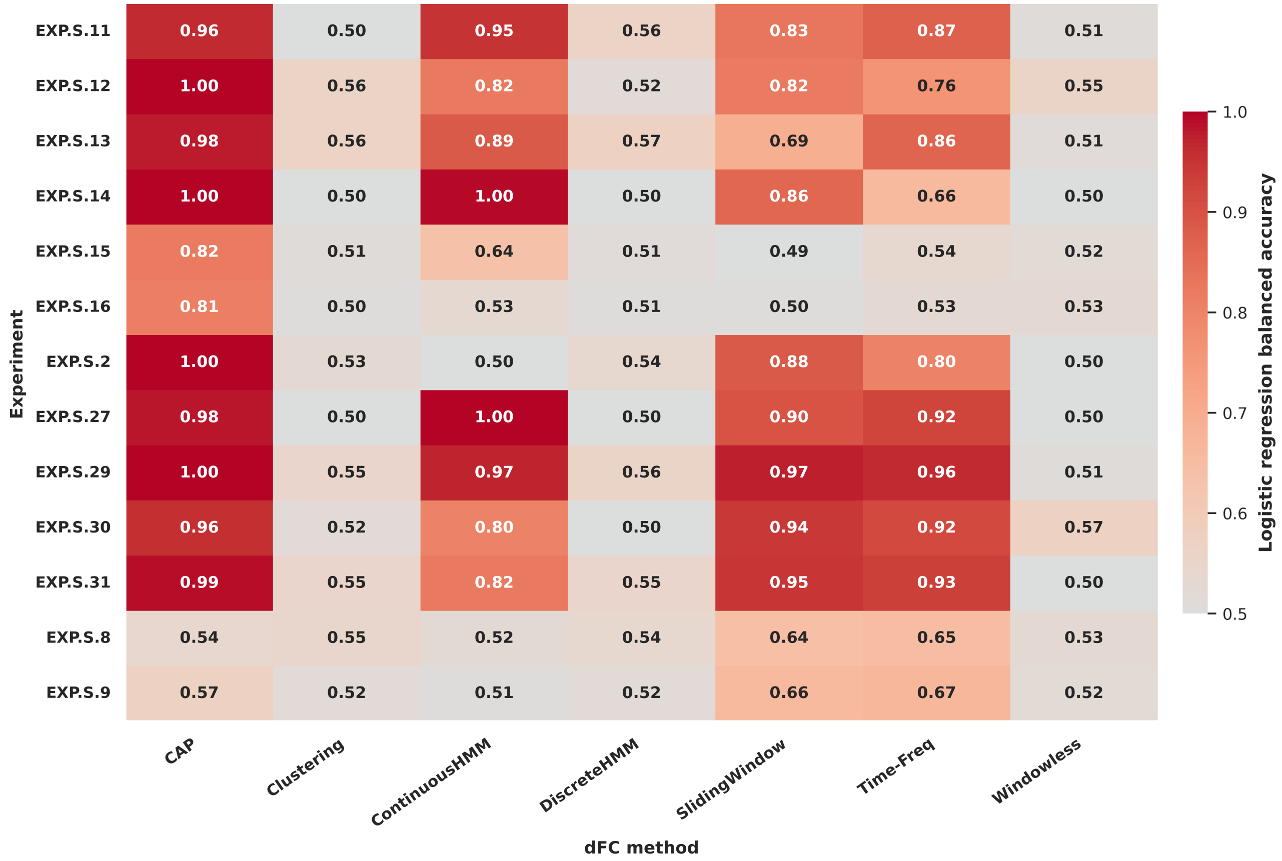


**Fig. 48. PLS-Logistic Regression heatmap for simulated experiments.** Heatmap representation of the results in Figure47.

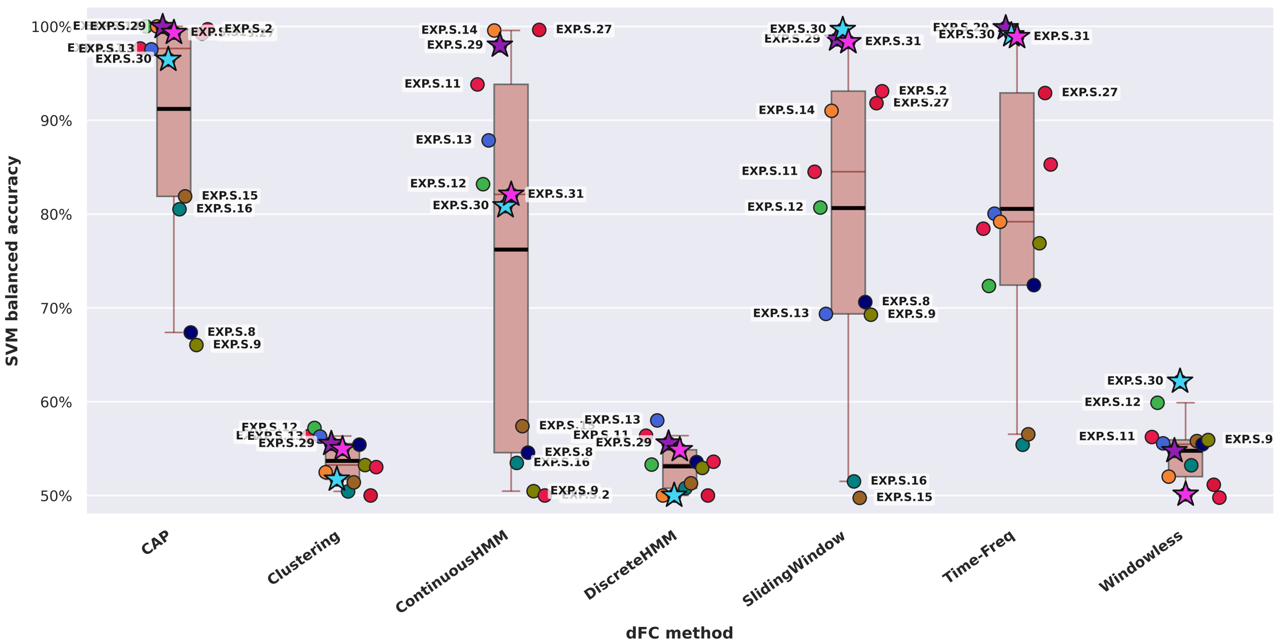


**Fig. 49. PCA-SVM classification performance for simulated experiments.** Contrary to experimental-data results, PCA yields higher accuracies than PLS for Sliding Window and Time-Frequency methods. Note: State-based methods retain the same values as in the PLS analyses since their four-dimensional feature spaces are not subjected to dimensionality reduction.

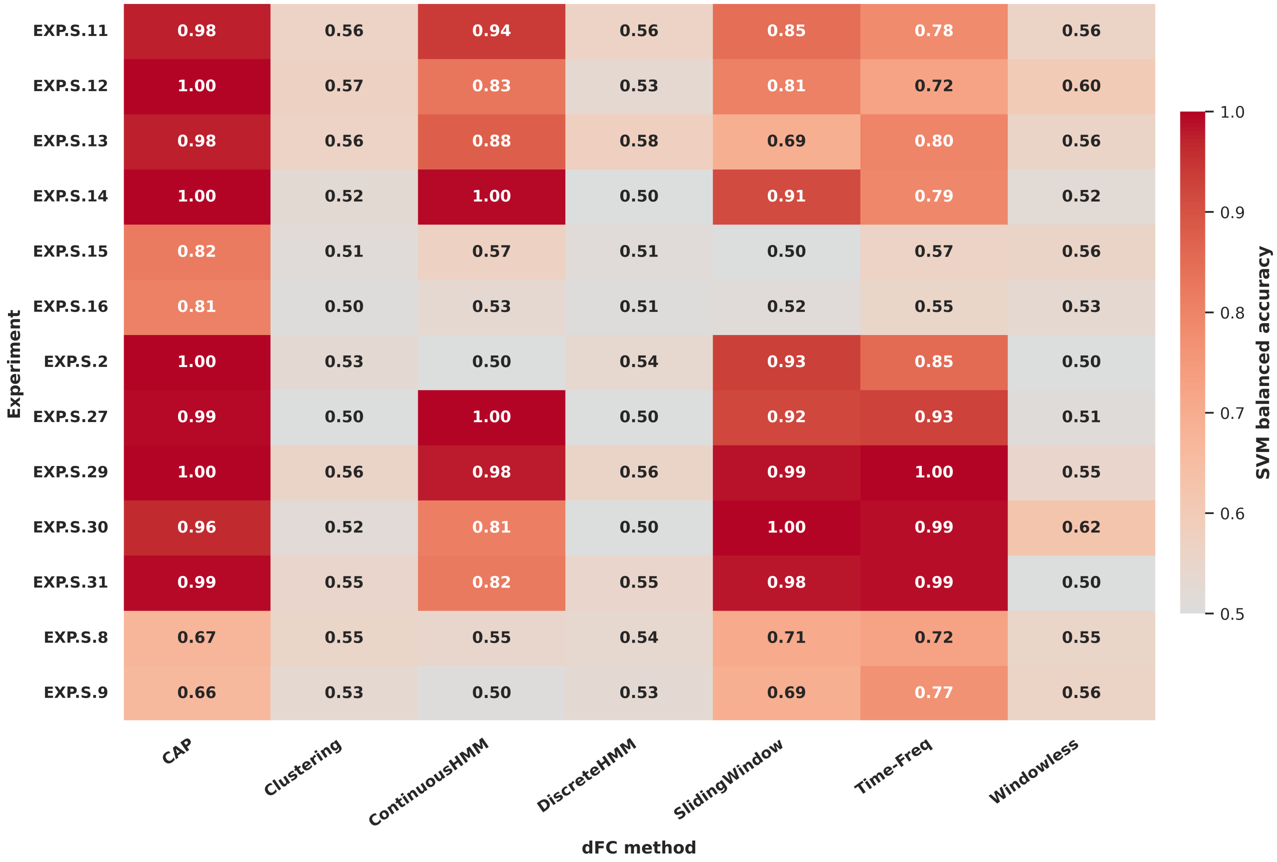


**Fig. 50. PCA-SVM heatmap for simulated experiments.** Heatmap visualization corresponding to Figure49.

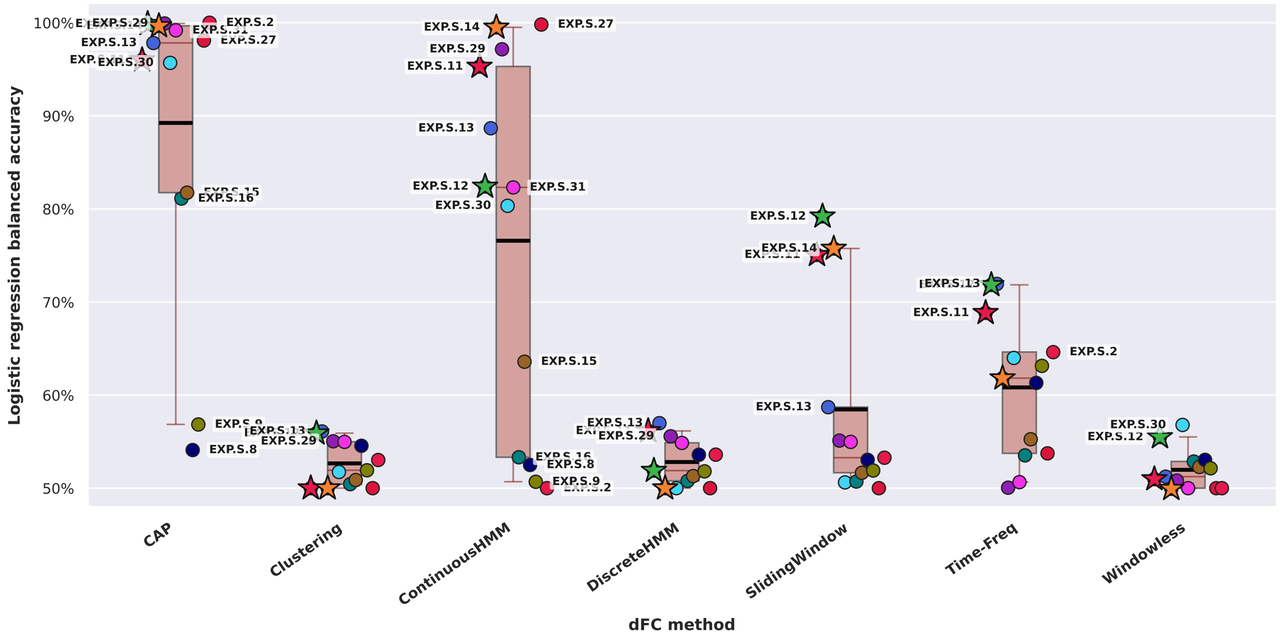


**Fig. 51. PCA-Logistic Regression classification performance for simulated experiments.** Sliding Window and Time-Frequency show reduced performance. Overall, logistic regression yields lower accuracies than SVM. Note: State-based methods retain the same values as in the PLS analyses since their four-dimensional feature spaces are not subjected to dimensionality reduction.

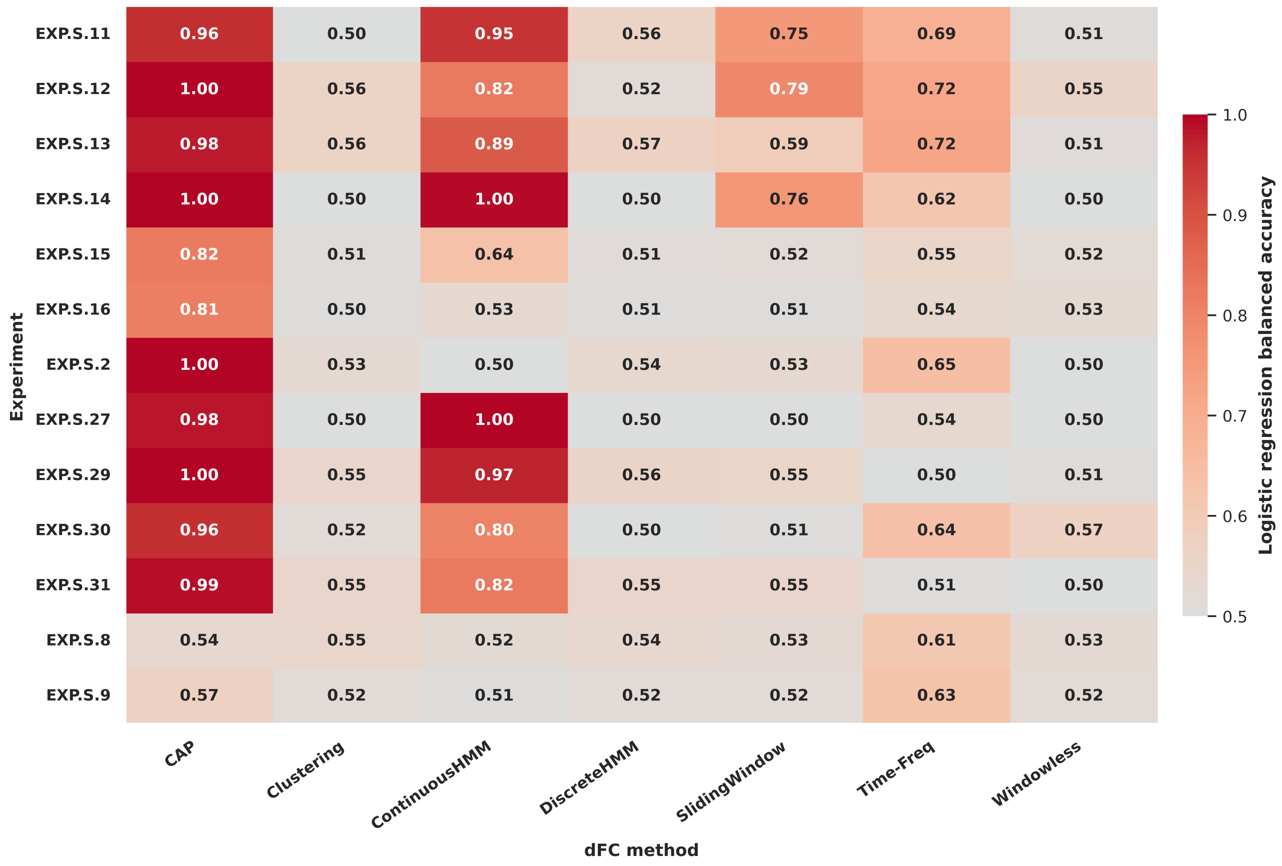


**Fig. 52. PCA-Logistic Regression heatmap for simulated experiments.** Heatmap representation corresponding to Figure51.

#### Silhouette Index (SI) Results for Experimental Data

To complement the classification analyses, this section evaluates how well dynamic functional connectivity (dFC) features separate task and rest samples without supervision. The Silhouette Index (SI) values quantify the intrinsic separability of cognitive states in the feature space, offering an unsupervised perspective on discriminative structure. Across experiments, SI values remain near zero for most methods and experiments. The highest separability appears for `EXP.18`, reaching SI = 0.35 with the Time-Frequency method. These results highlight that, although supervised models can decode task presence effectively, intrinsic class separation in raw dFC features is limited-especially in experimental, noisy data. *Note: For experiments with multiple runs, all reported values correspond to the best-performing run, unless stated otherwise.*

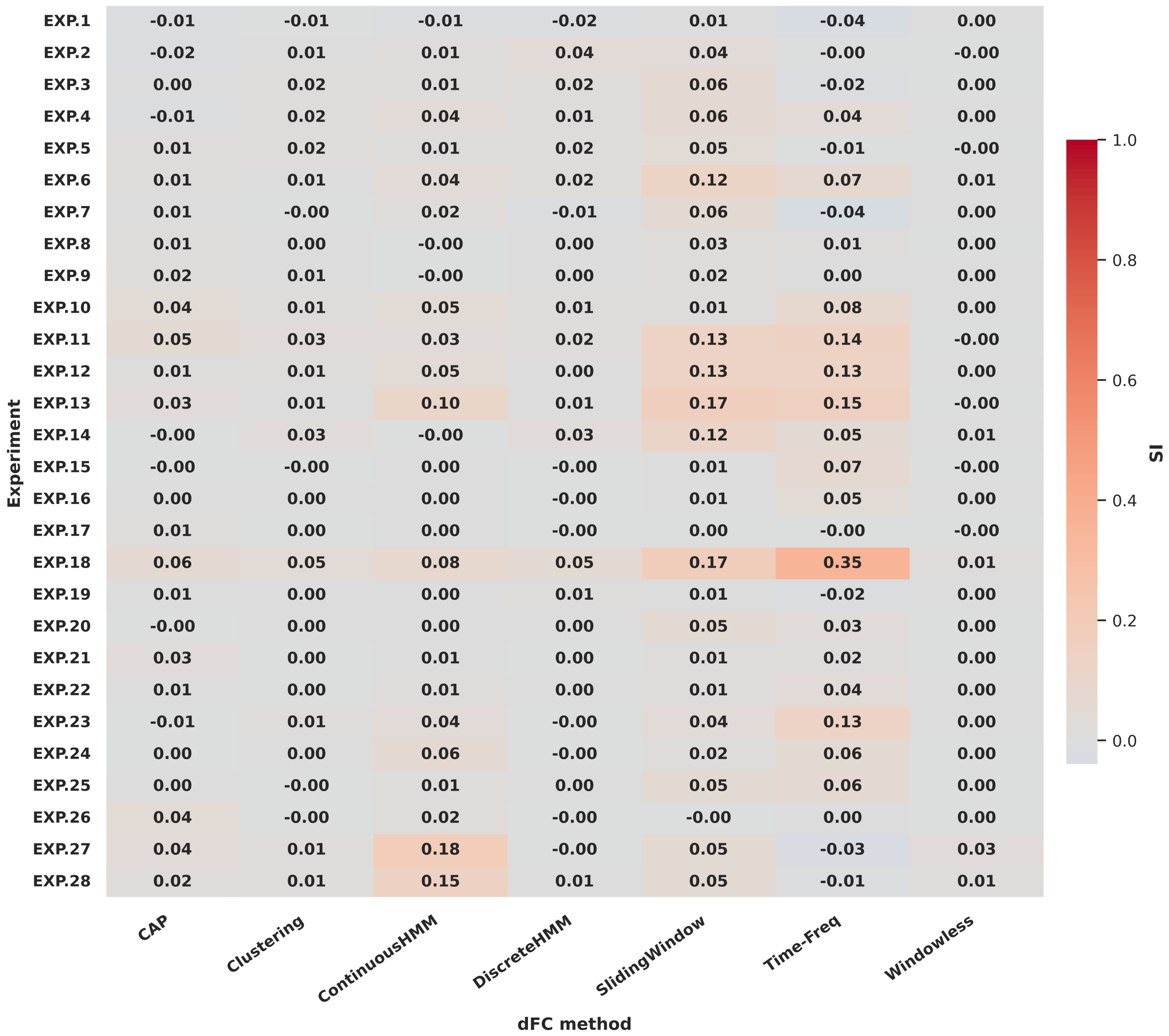


**Fig. 53. PLS-based SI values across experiments from experimental datasets.** Heatmap of SI scores for each dFC method and experiment, using PLS reduction. Most values are close to zero, indicating weak unsupervised separability between rest and task conditions. `EXP.18` shows the strongest separation, with SI = 0.35 with the Time-Frequency method.

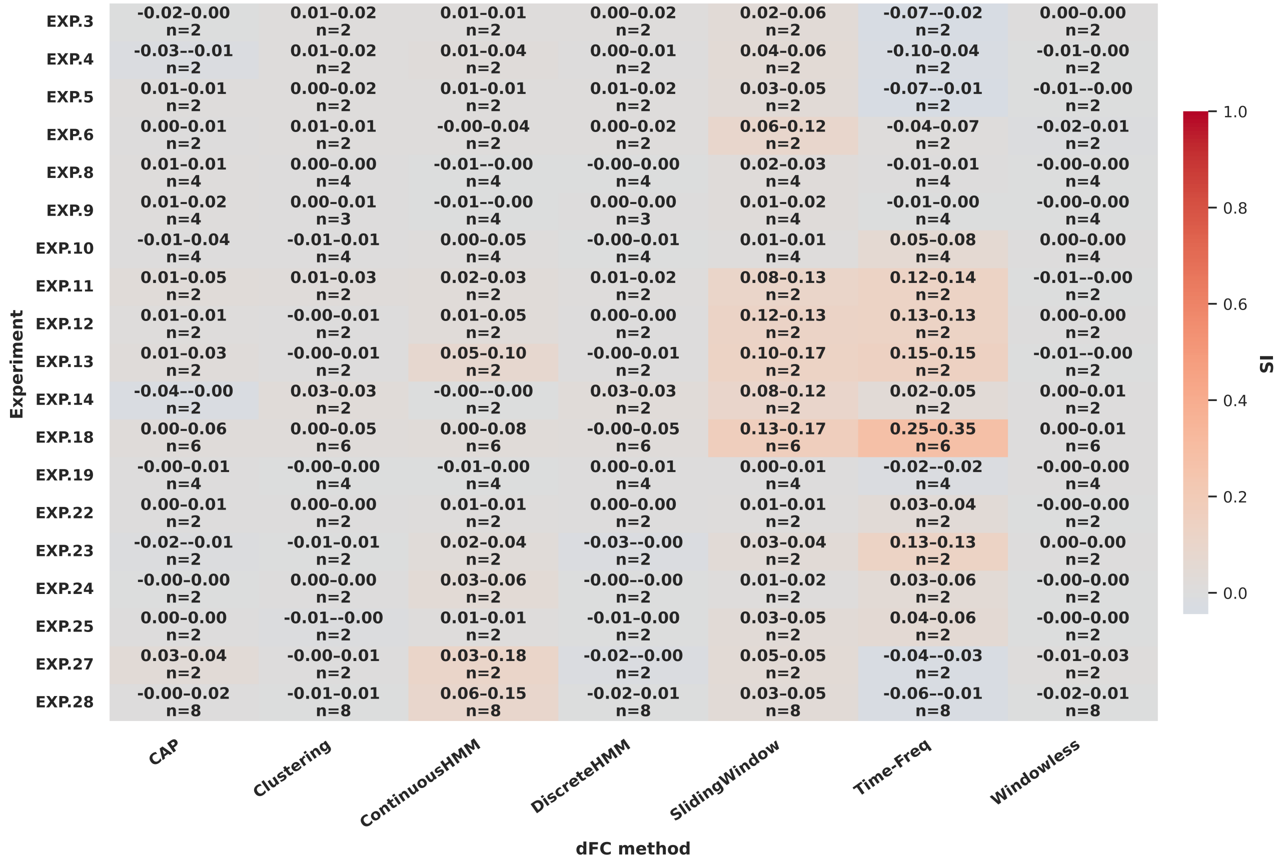


**Fig. 54. Across-run stability of SI.** Heatmap summarizing task-rest separability using SI score based on PLS-transformed dFC features. Each cell corresponds to an experiment-method pair, with color representing the median SI across runs, and in-cell annotations showing the minimum and maximum values (min-max) and the number of available runs (n). Narrow min-max ranges indicate high run-to-run stability, whereas wide ranges signal variability even when the median SI is high. Robust high separability corresponds to warm hues with tight ranges, reflecting consistent task-rest separability across runs.

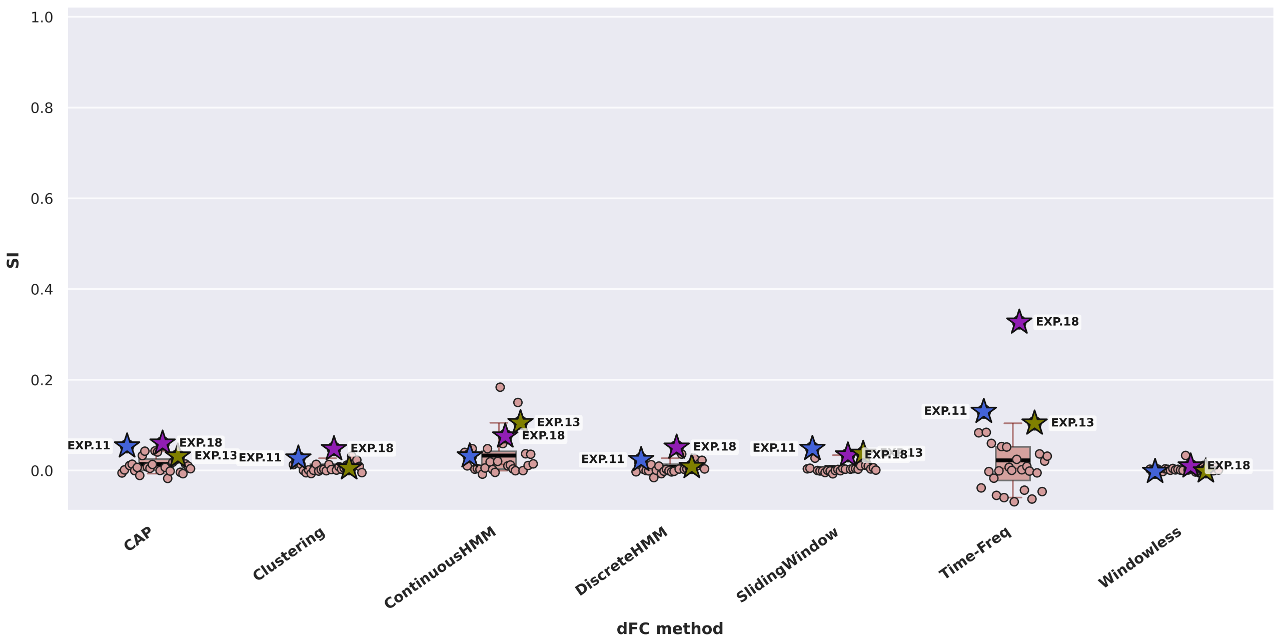


**Fig. 55. PCA-based Silhouette Index (SI) distribution across experiments from experimental datasets.** Boxplots and stripplots of SI values for all dFC methods after PCA reduction. Values remain very low overall, often near zero, showing minimal class separation in the linear subspace. For `EXP.18`, the Time-Frequency method yields a modest SI of 0.33, similar to PLS-based SI. Note: State-based methods retain the same values as in the PLS analyses since their four-dimensional feature spaces are not subjected to dimensionality reduction.

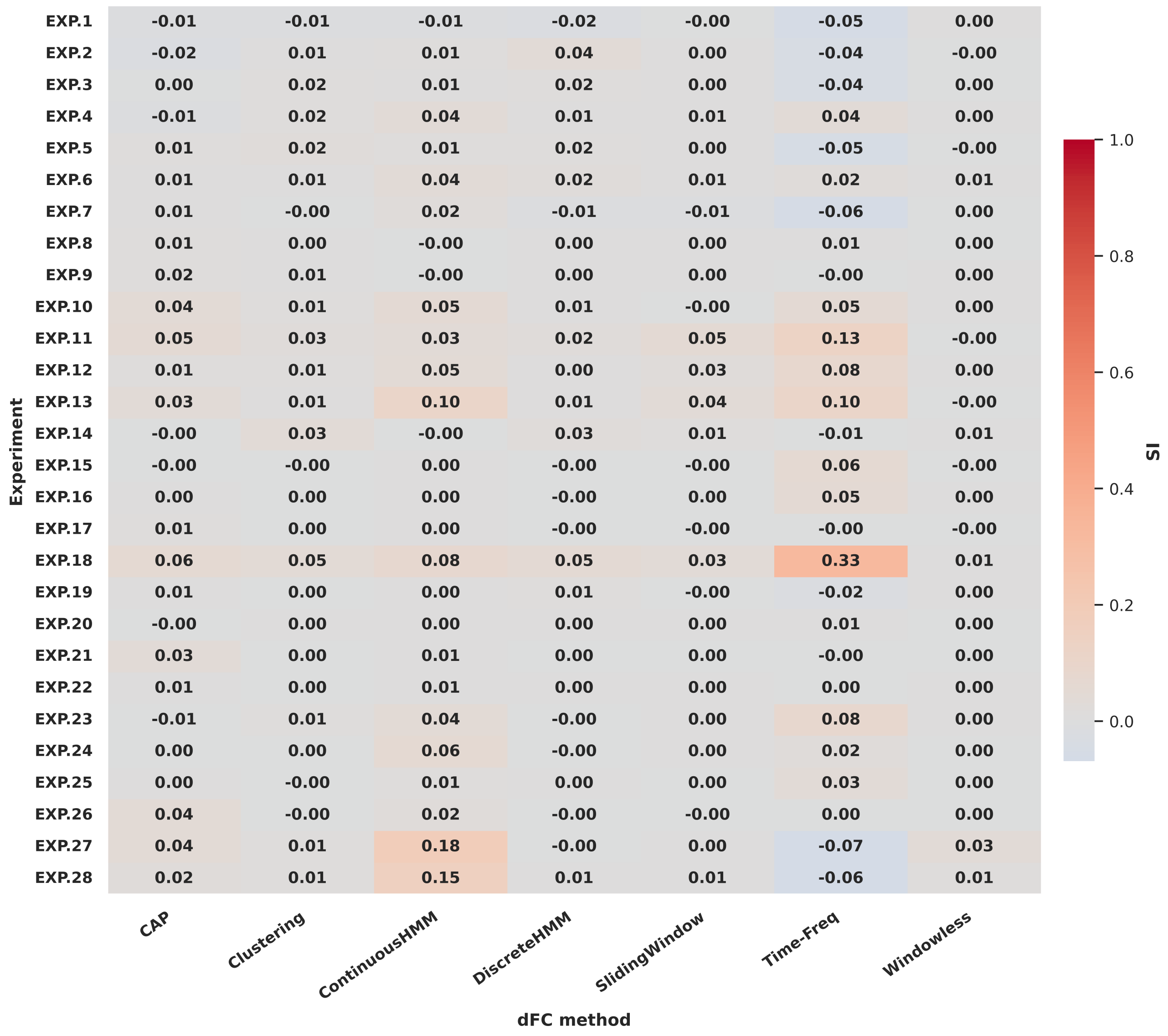


**Fig. 56. PCA-based Silhouette Index (SI) heatmap for experiments from experimental datasets.** Heatmap corresponding to Figure55, visualizing SI values per method and experiment.

#### Silhouette Index (SI) Results for Simulated Data

To complement the classification analyses, this section examines task-rest separability in simulated data using the Silhouette Index (SI), which quantifies the intrinsic distinction between task and rest samples in the feature space. Compared to real-world experiments, simulated experiments yield markedly higher SI values, reflecting idealized conditions with clearer neural responses and higher signal-to-noise ratio. `EXP.S.27` achieves the strongest separability overall, reaching SI = 0.61 with Continuous HMM and SI = 0.60 with CAP.

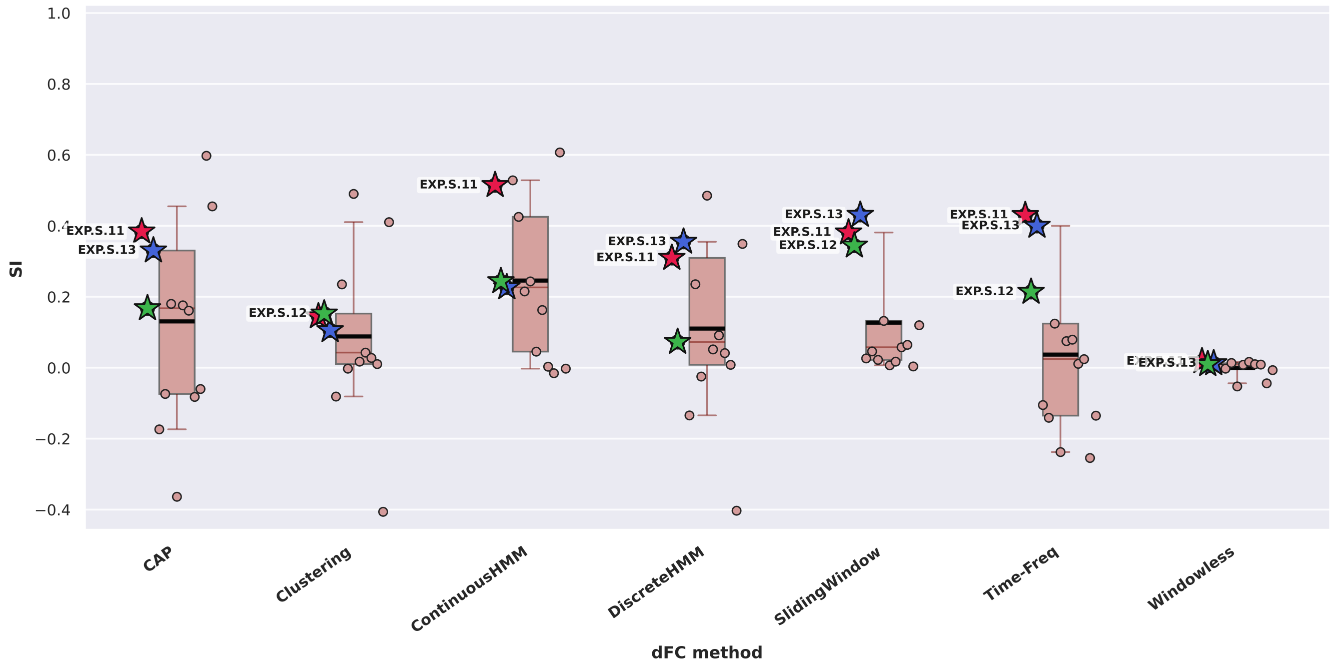


**Fig. 57. Task-rest separability across dFC methods in simulated datasets, quantified by SI using PLS-transformed dFC features.** Each boxplot summarizes the distribution of SI scores measured using PLS-transformed dFC features across simulated experiments for a given dFC method, with median (red line) and mean (black line) indicated. Individual points correspond to different experiments. The three experiments with the highest separability overall are marked with a star.

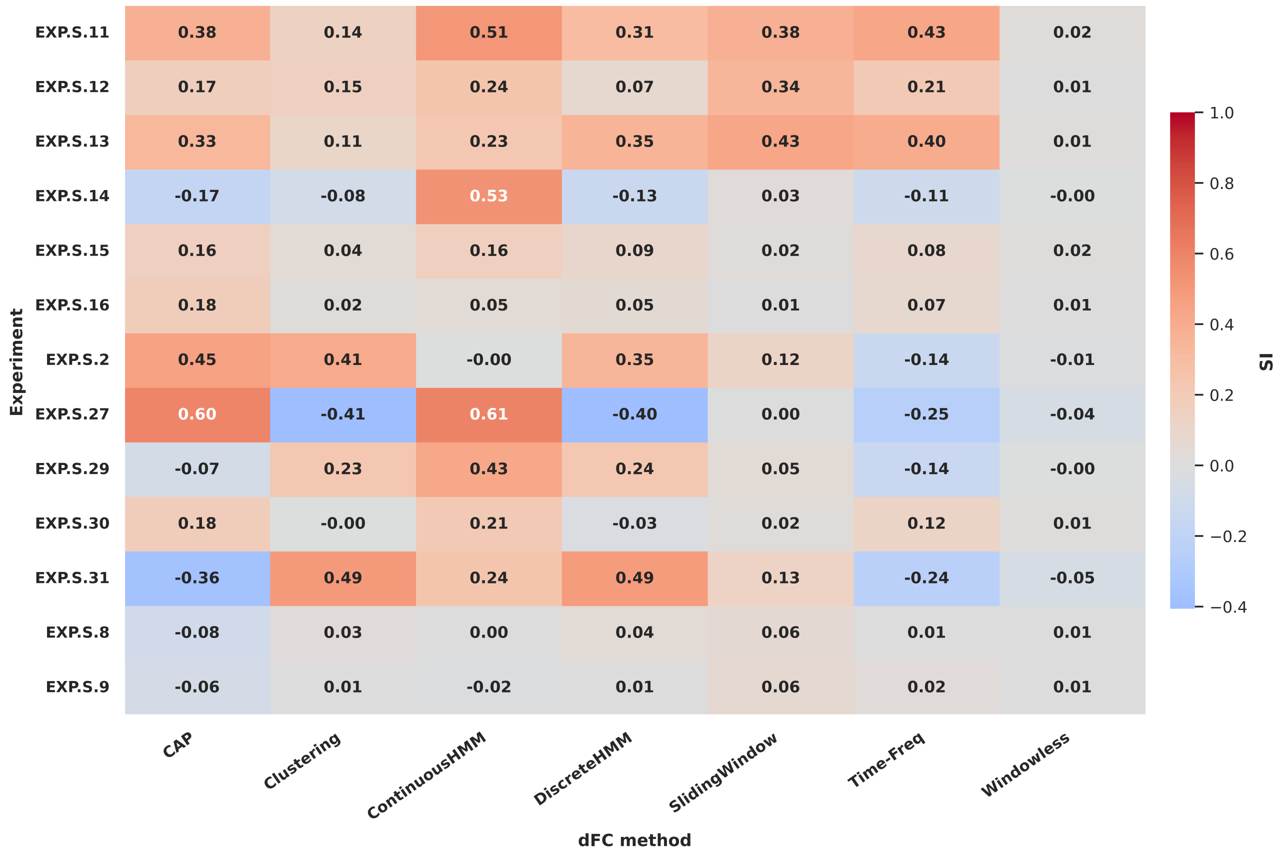


**Fig. 58. PLS-based Silhouette Index (SI) heatmap across simulated experiments.** Heatmap corresponding to Figure57, visualizing SI values per method and experiment.

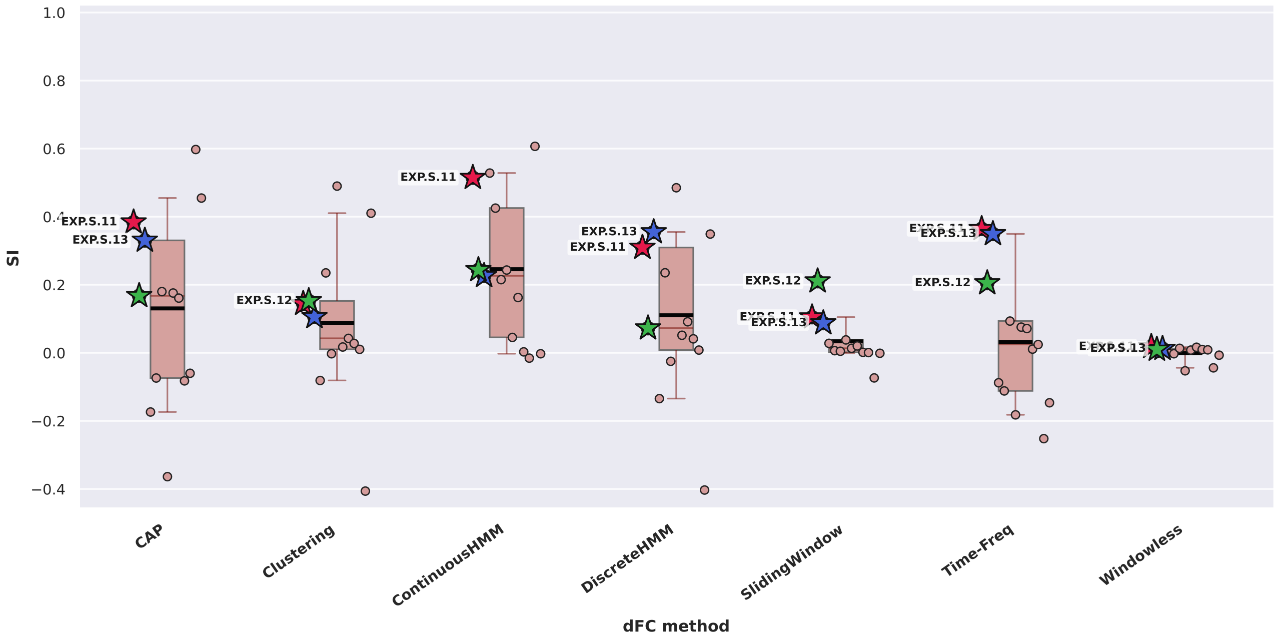


**Fig. 59. Task-rest separability across dFC methods in simulated datasets, quantified by SI using PCA-transformed dFC features.** Each boxplot summarizes the distribution of SI scores measured using PCA-transformed dFC features across simulated experiments for a given dFC method, with median (red line) and mean (black line) indicated. Individual points correspond to different experiments. The three experiments with the highest separability overall are marked with a star. Note: State-based methods retain the same values as in Figure57 since their four-dimensional feature spaces are not subjected to dimensionality reduction.

| Experiment | CAP | Clustering | ContinuousHMM | DiscreteHMM | SlidingWindow | Time-Freq | Windowless |
|---|---|---|---|---|---|---|---|
| EXP.S.11 | 0.38 | 0.14 | 0.51 | 0.31 | 0.10 | 0.36 | 0.02 |
| EXP.S.12 | 0.17 | 0.15 | 0.24 | 0.07 | 0.21 | 0.21 | 0.01 |
| EXP.S.13 | 0.33 | 0.11 | 0.23 | 0.35 | 0.09 | 0.35 | 0.01 |
| EXP.S.14 | -0.17 | -0.08 | 0.53 | -0.13 | 0.03 | -0.09 | -0.00 |
| EXP.S.15 | 0.16 | 0.04 | 0.16 | 0.09 | 0.02 | 0.07 | 0.02 |
| EXP.S.16 | 0.18 | 0.02 | 0.05 | 0.05 | 0.01 | 0.08 | 0.01 |
| EXP.S.2 | 0.45 | 0.41 | -0.00 | 0.35 | -0.00 | -0.15 | -0.01 |
| EXP.S.27 | 0.60 | -0.41 | 0.61 | -0.40 | -0.07 | -0.25 | -0.04 |
| EXP.S.29 | -0.07 | 0.23 | 0.43 | 0.24 | 0.01 | -0.11 | -0.00 |
| EXP.S.30 | 0.18 | -0.00 | 0.21 | -0.03 | 0.00 | 0.09 | 0.01 |
| EXP.S.31 | -0.36 | 0.49 | 0.24 | 0.49 | 0.04 | -0.18 | -0.05 |
| EXP.S.8 | -0.08 | 0.03 | 0.00 | 0.04 | 0.00 | 0.01 | 0.01 |
| EXP.S.9 | -0.06 | 0.01 | -0.02 | 0.01 | 0.00 | 0.02 | 0.01 |

**Fig. 60. PCA-based Silhouette Index (SI) heatmap across simulated experiments.** Heatmap corresponding to Figure59, visualizing SI values per method and experiment.
